# A massive 7T fMRI dataset to bridge cognitive and computational neuroscience

**DOI:** 10.1101/2021.02.22.432340

**Authors:** Emily J. Allen, Ghislain St-Yves, Yihan Wu, Jesse L. Breedlove, Logan T. Dowdle, Brad Caron, Franco Pestilli, Ian Charest, J. Benjamin Hutchinson, Thomas Naselaris, Kendrick Kay

**Author notes:** Co-senior author (Thomas Naselaris, Kendrick Kay). Department of Neuroscience, University of Minnesota, Minneapolis, Minnesota, USA. Department of Psychology, University of Minnesota, Minneapolis, Minnesota, USA.

## Abstract

Extensive sampling of neural activity during rich cognitive phenomena is critical for robust understanding of brain function. We present the Natural Scenes Dataset (NSD), in which high-resolution fMRI responses to tens of thousands of richly annotated natural scenes are measured while participants perform a continuous recognition task. To optimize data quality, we develop and apply novel estimation and denoising techniques. Simple visual inspections of the NSD data reveal clear representational transformations along the ventral visual pathway. Further exemplifying the inferential power of the dataset, we use NSD to build and train deep neural network models that predict brain activity more accurately than state-of-the-art models from computer vision. NSD also includes substantial resting-state and diffusion data, enabling network neuroscience perspectives to constrain and enhance models of perception and memory. Given its unprecedented scale, quality, and breadth, NSD opens new avenues of inquiry in cognitive and computational neuroscience.

## Introduction

Neuroscience has an insatiable appetite for data. Many ongoing efforts to extensively sample brain activity (de Vries et al., 2020; Siegle et al., 2021; Stringer et al., 2019) and structure (Markram et al., 2015; Van Essen et al., 2013; Zheng et al., 2018) are motivated, in part, by the availability of new computational methods that make analysis of massive datasets feasible (Jonathan Pillow and Sahani, 2019). Equally as important is the growing desire to understand how the brain coordinates complex sensory and motor behaviors and the realization that the neural networks supporting such behaviors span multiple scales, from single neurons to local circuits to whole systems (Bassett and Sporns, 2017). Understanding massive, complex networks will inevitably require commensurately massive amounts of data.

The need for massive data is especially acute in visual neuroscience, a model system for understanding brain function. The network that mediates our ability to flexibly and efficiently perceive the visual world occupies approximately one-third of human cerebral cortex (Van Essen et al., 2001) and interconnects brain areas with profoundly different functional properties (Grill-Spector and Malach, 2004). This network both encodes visual stimuli and interfaces visual representations into a cognitive context, including information about what one has already seen (Wheeler et al., 2000), might see (Breedlove et al., 2020), or is selectively attending (Kay et al., 2015). Understanding vision thus means interrogating a high-dimensional, context-dependent neural network.

Given these considerations, it is clear that extensive experimental data providing access to whole-brain responses to complex stimuli are critical in the quest to understand the human visual system. The ideal dataset should include naturalistic stimuli: the visual system is distributed widely across the brain, and natural scenes, in addition to being ecologically relevant (Geisler, 2008), are effective activators of the entire system (Huth et al., 2012). Moreover, the ideal dataset should be large: in order to take full advantage of powerful data analysis and machine learning (ML) techniques that have recently become available (Vu et al., 2018), we need considerably more data than is currently available. How much? Modern ML methods used in computer vision to process natural scenes (e.g. deep convolutional neural networks) require tens to hundreds of thousands of image samples for training (Krizhevsky, 2009; Lin et al., 2014). A dataset that sampled brain activity at these scales would raise the exciting possibility of exploiting these methods to develop better models of how the brain processes natural scenes (Güçlü and van Gerven, 2015; Han et al., 2019; Khaligh-Razavi and Kriegeskorte, 2014; Seeliger et al., 2021; Stansbury et al., 2013; St-Yves and Naselaris, 2017; Yamins et al., 2014; Zhuang et al., 2021), and would accelerate efforts to bridge computational and cognitive neuroscience (Naselaris et al., 2018).

In this paper, we present the first dataset that achieves sampling at this ambitious scale. The Natural Scenes Dataset (NSD) consists of high-resolution (1.8 mm) whole-brain 7T fMRI of 8 carefully screened human participants who each viewed 9,000–10,000 color natural scenes (22,000–30,000 trials) during 30–40 scan sessions distributed over the course of a year. Aggregated across participants, NSD includes responses to 70,566 distinct natural scene images—this is more than an order of magnitude larger than comparable datasets involving fMRI sampling of many images (Chang et al., 2019; Horikawa and Kamitani, 2017; Kay et al., 2008). Moreover, as we show, the high quality of the NSD dataset (afforded, in part, by the use of ultra-high magnetic field strength) makes it possible to leverage the full power of modern ML methods for developing better models of visual representation.

NSD incorporates several innovations in addition to its unprecedented scale and quality. To reconcile extensive sampling with a practical time commitment, we used an aggressive rapid event-related design. This drove the development of new analysis techniques that accurately compensate for the overlap of hemodynamic responses across successive trials. To ensure participant engagement and control cognitive state, we incorporated a continuous recognition task (Brady et al., 2008) in which participants were instructed to indicate whether they have seen each presented image at any point in the past. In addition to making the experiment tolerable (and even somewhat interesting) for participants, inclusion of this task makes NSD, to our knowledge, the longest-term continuous recognition memory fMRI study in history and, thus, a likely source of new insights into long-term memory formation and the cognitive context of vision. Finally, to ensure the broad reach of the NSD dataset, we incorporated design input from a large network of collaborators with diverse scientific interests (e.g., low-level vision, high-level vision, memory, connectivity, neuroanatomy) and technical expertise (e.g., mapping, multivariate pattern analysis, encoding models, representational similarity analysis, neural network modeling). This input helped precipitate a carefully curated dataset with extensive auxiliary measures, thereby increasing the likelihood that NSD will enjoy widespread application in cognitive and computational neuroscience.

## Results

### Sampling thousands of images during a continuous recognition task

We obtained 73,000 color natural scenes from the richly annotated Microsoft Common Objects in Context (COCO) image dataset (Lin et al., 2014), a dataset that is heavily used in the computer vision and machine learning communities. Our experimental design specified that each of 8 subjects would view 10,000 distinct images and a special set of 1,000 images would be shared across subjects (8 subjects × 9,000 unique images + 1,000 shared images = 73,000 images). This sampling strategy was chosen to maximize the number of distinct images in NSD. Each image would be presented 3 times to a given subject. While this is a low number, we reasoned that 3 trials would be sufficient to produce robust responses given our use of ultra-high field (7T) fMRI. Furthermore, images would be presented using a rapid event-related design consisting of 4-s trials (**Figure 1A**). This was done to maximize statistical power and to create an engaging experience for the subjects. In addition, the continuous nature of task engagement helps avoid unwanted arousal-related fMRI signals (Roth et al., 2020).

**Figure 1.**
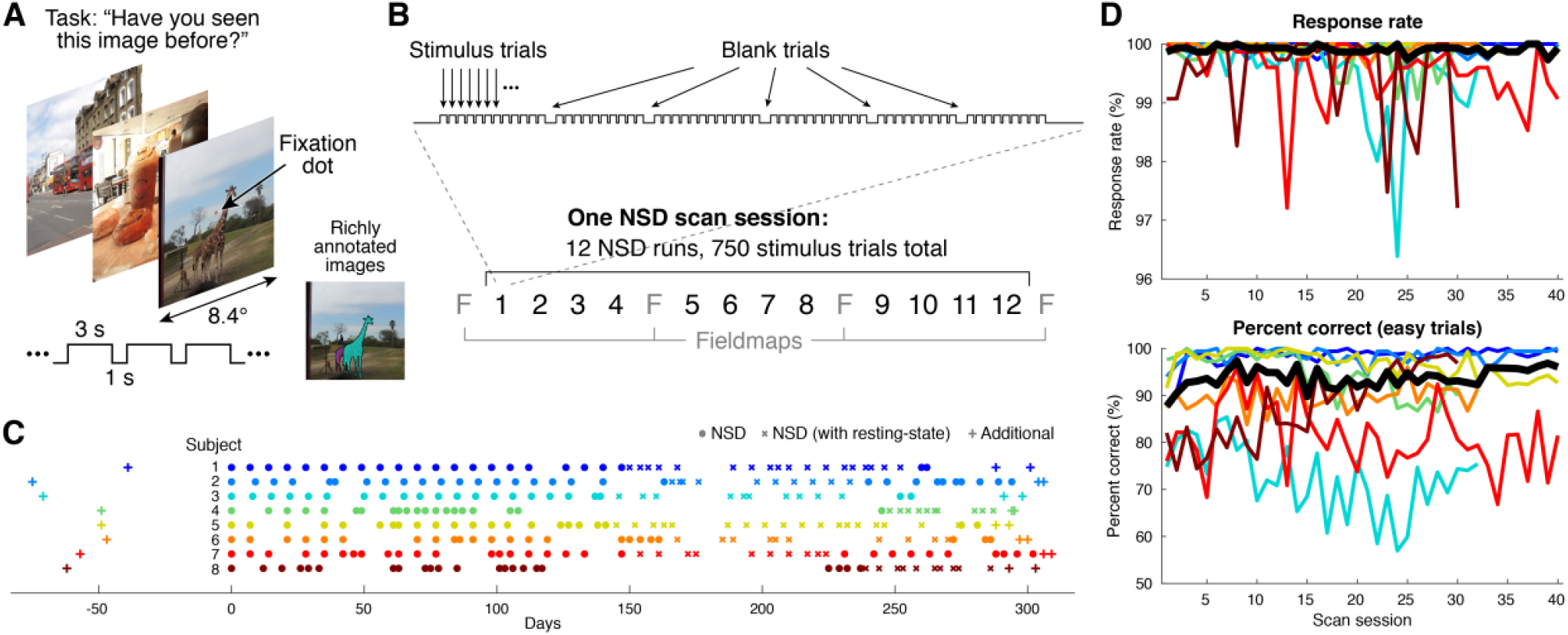
Design of the NSD experiment. *A*, Trial design. While maintaining central fixation, participants viewed sequences of color natural scenes and judged whether each image had been previously shown at any point in the past. The scenes, taken from Microsoft’s COCO (Lin et al., 2014), are richly annotated with object information (as depicted). *B*, Run and session design. Each run lasted 5 minutes and consisted of 62 or 63 stimulus trials with occasional interspersed blank trials. Each scan session consisted of 12 runs (750 stimulus trials). *C*, Timeline of 7T fMRI scan sessions. Each subject participated in an initial screening session (prffloc), 30–40 NSD core sessions, and two final sessions (nsdsynthetic, nsdimagery). The first NSD core session corresponds to day 0. *D*, Behavioral compliance. Results for individual subjects (thin colored lines) and the median across subjects (thick black line) are shown. Easy trials are defined as trials that involved the presentation of an image that had occurred earlier in the same scan session.

The NSD experiment was split across 40 scan sessions for each subject (**Figure 1B**). To control cognitive state and encourage deep processing of the images, subjects were instructed to perform a continuous recognition task in which they reported whether the current image had been presented at any previous point in the experiment. We controlled the distributions of image presentations such that both short-term and long-term repetitions were probed (**Supplementary Figure 1A**). Parameters were selected such that even in the first scan session, images were not always new, and even in the last scan session, images were not always old (**Supplementary Figure 1B**).

### Neuroimaging data collection on carefully selected subjects

All fMRI data in NSD were collected at 7T using a whole-brain 1.8-mm 1.6-s gradient-echo EPI pulse sequence. After verbally screening a number of potential participants with respect to basic eligibility criteria, we recruited 14 subjects to participate in an initial 7T fMRI screening session which involved population receptive field (pRF) (Benson et al., 2018) and category localizer (fLoc) (Stigliani et al., 2015) experiments. Based on data from this scan session, we ranked the 14 subjects with respect to data quality and invited the top 8 subjects to participate in the full NSD experiment (all subjects accepted). This selection process was conducted to ensure the best possible data quality for NSD. Analyses conducted after completion of the NSD experiment confirm that the ranking procedure successfully identified subjects that yield high-quality data and that data quality would have suffered substantially had we omitted the selection process (**Figure 2C**).

**Figure 2.**
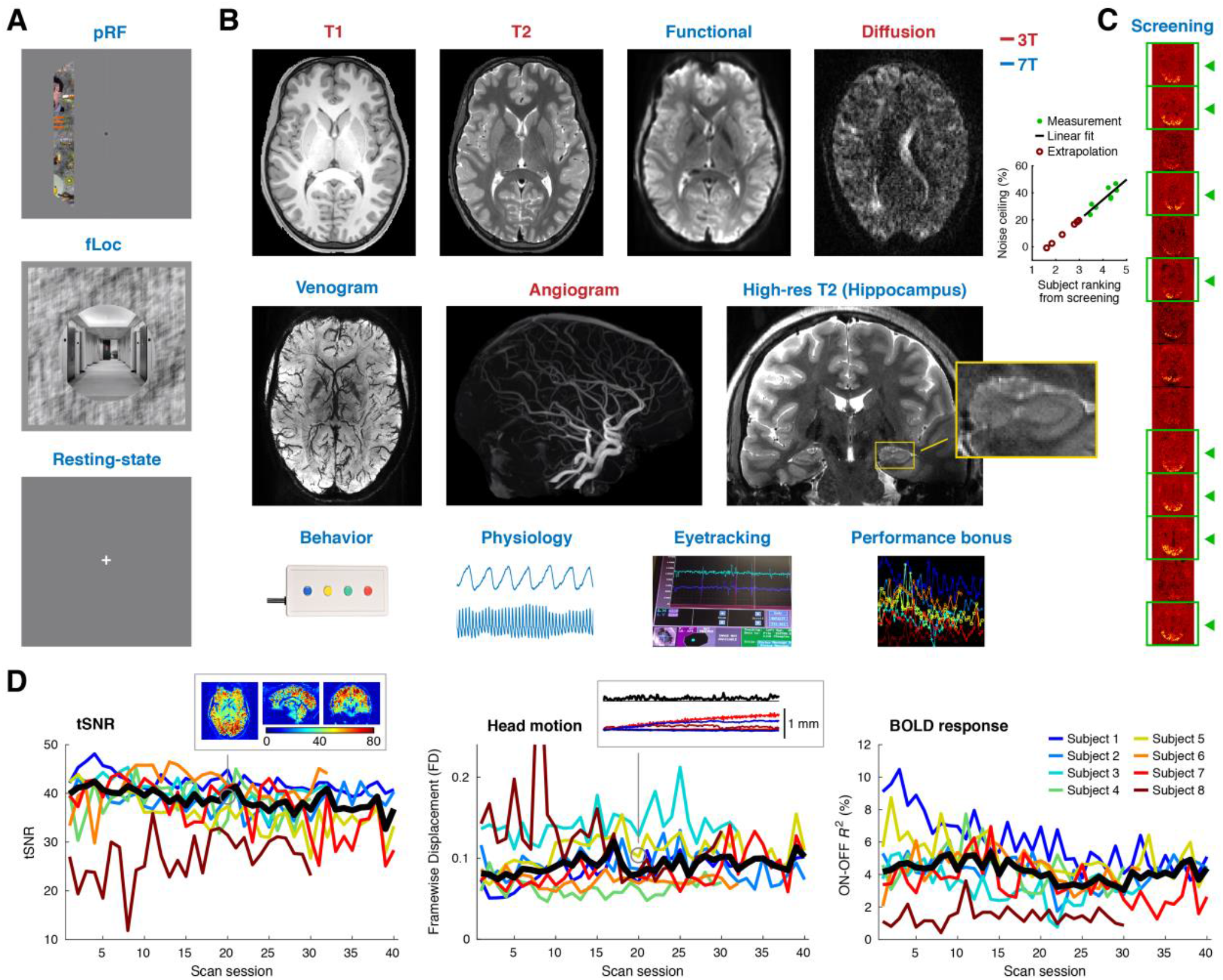
Overview of acquired data. *A*, Auxiliary fMRI experiments. Data from the pRF and fLoc experiments were used to define retinotopic visual areas and category-selective regions, respectively. Resting-state data were collected before and after the NSD runs in a subset of the NSD core sessions (totaling 100 or 180 minutes per subject). *B*, Available measures. Examples of the actual data are depicted. *C*, Participant selection. Data quality from the initial screening session was used to rank a set of 14 participants. On the right is an illustration of one measure contributing to the ranking, specifically, variance explained in the fLoc experiment (one slice per participant; identical color range). The inset compares the participant ranking against the b3 noise ceiling calculated on the full NSD dataset (see **Figure 5**). A line fit to the 8 NSD subjects (green dots) is extrapolated to predict noise ceilings for the subjects who were not selected for participation in NSD (red circles). *D*, Metrics of data quality. Results for individual subjects (thin colored lines) and the median across subjects (thick black line) are shown. The insets show detail on tSNR and head motion for one sample run (see **Supplementary Figures 3–4** for more information).

Data were collected from the 8 NSD subjects over the course of a year (**Figure 1C**). Subjects consistently engaged with the task as indicated by the near perfect response rates (**Figure 1D, top**). Moreover, the subjects correctly recognized items that were repeated within a given scan session (“easy trials”; **Figure 1D, bottom**) at rates higher than a 50/50 guess rate (for details on recognition memory, see **Figure 6B**). The full NSD dataset includes a variety of anatomical neuroimaging measures (including *T*_1_, *T*_2_, diffusion, SWI, and TOF), functional neuroimaging measures (including the pRF and fLoc experiments, the NSD experiment, resting-state data, and two additional experiments involving synthetic stimuli and visual imagery), and behavioral measures (**Figure 2A–B**). In some fMRI sessions, physiological and eyetracking data were also collected. With regards to the core NSD experiment, we completed the full set of 40 NSD scan sessions for four of the subjects, but due to unforeseen summer absences and scheduled decommissioning of the 7T scanner, we completed 30–32 NSD scan sessions for each of the other subjects. A full breakdown of data collection is provided in **Supplementary Figure 2**.

### Stable high-resolution functional imaging across scan sessions

We believe visual inspection is the most effective way to assess many common aspects of fMRI pre-processing (Kay et al., 2019). Accordingly, we generated a comprehensive set of movies that detail the excellent quality of the pre-processed NSD dataset. These include movies that assess the co-registration of the different imaging modalities (e.g. *T*_1_, *T*_2_, EPI; **Supplementary Video 1**); movies that assess the manually-edited cortical surface reconstructions generated using FreeSurfer (**Supplementary Video 2**); movies that assess the registration of the NSD subjects to the fsaverage (**Supplementary Video 3**) and MNI (**Supplementary Video 4**) group spaces; movies that inspect raw and pre-processed EPI volumes (**Supplementary Video 5**); and movies that provide volume and surface visualizations of the stability of mean EPI intensity across sessions (**Supplementary Videos 6 and 7; Supplementary Figure 5**) and the stability of BOLD responses across sessions (**Supplementary Videos 8 and 9**). The movies demonstrate that the quality of the NSD data enables precision functional mapping (Gordon et al., 2017): activity patterns are fine-scale and highly reliable within individual subjects and these patterns are distinct across subjects.

In addition to visual inspection, quantitative data quality metrics were computed for each NSD scan session. This was in fact done on a rolling basis as the data were acquired, allowing us to monitor data quality and provide performance bonuses to the subjects. Inspecting the metrics, we see that tSNR is stable across scan sessions for each subject (**Figure 2D, left**). One subject, subject 8, exhibits low tSNR compared to the other subjects; this can be attributed to higher levels of head motion for this subject (Satterthwaite et al., 2014) (**Figure 2D, middle**). We also observe that BOLD responses are stable across scan sessions for each subject, though there is substantial variation in the strength of BOLD responses across subjects (**Figure 2D, right**).

One feature we implemented in the pre-processing of the fMRI data was to interpolate the data on both a fine temporal grid and a fine spatial grid. This upsampling strategy preserves fine-scale detail that is present in the raw fMRI data due to the temporal jitter of the acquired fMRI volumes relative to the experimental paradigm (Watanabe et al., 2013) and the spatial jitter of the acquired fMRI volumes relative to the brain’s anatomy (Kang et al., 2007; Kay et al., 2019). An illustration of the benefits of upsampling is provided in **Figure 3**. This example highlights the existence of replicable, fine-scale detail in fMRI image intensities as well as in BOLD responses extracted from the fMRI data.

**Figure 3.**
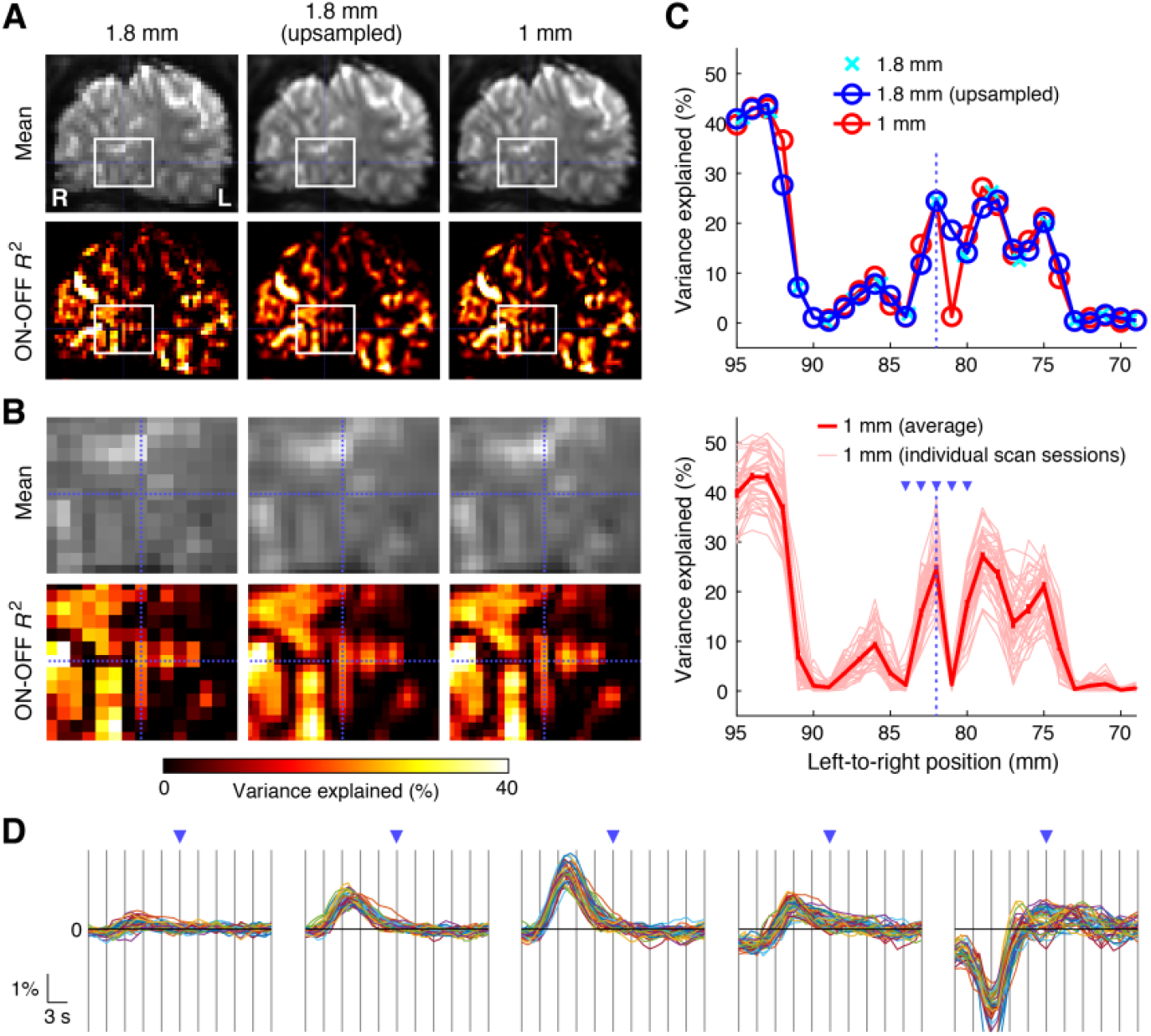
Improvements in spatial detail through upsampling. *A*, Comparison of approaches. For an example coronal slice in Subject 1, we compare the non-upsampled 1.8-mm preparation of the data (left), the upsampled 1-mm preparation (right), and a post-hoc upsampled version of the 1.8-mm results (middle). Two quantities are shown: mean signal intensity and variance explained by an ON-OFF GLM model. *B*, Zoomed view of white rectangle marked in panel A. *C*, Profile view of blue dotted horizontal line marked in panel B. Error bars in the bottom plot indicate ± 1 SEM across sessions. *D*, Timecourse estimates for voxels marked by blue arrows in panel C. Each trace corresponds to an estimate of the hemodynamic timecourse for a single voxel in one NSD scan session.

### Highly reliable diffusion data and derivatives

The diffusion data included with the NSD dataset complements the extensive fMRI measurements. We pre-processed the raw diffusion data using the state-of-the-art DESiGNER pipeline methodology (Ades-Aron et al., 2018) as implemented on brainlife.io (Avesani et al., 2019). We find that the quality of the pre-processed diffusion data for each subject is high, as evidenced by the signal-to-noise ratio (**Supplementary Figure 6B**). We then proceeded to perform diffusion signal modeling (Daducci et al., 2015; Jensen and Helpern, 2010; Pierpaoli et al., 1996; Tournier et al., 2007; Zhang et al., 2012), anatomically-informed tractography (Smith et al., 2012), and profilometry (Yeatman et al., 2012). White-matter microstructural properties are found to be highly reliable for each subject (**Figure 4A**). Structural connectivity matrices (Hagmann et al., 2008) derived from tractography results are also highly reliable, both at the group level (**Figure 4B–C**) as well as at the single-subject level (**Figure 4C, inset**). The ready-to-use diffusion derivatives provided with NSD include a variety of macrostructural and microstructural measures, white-matter tracts, and structural connectivity matrices. These derivatives can easily integrated into machine learning workflows, and serve as launching points for scientific investigations seeking to apply network neuroscience perspectives (Bassett and Sporns, 2017) to understanding brain function in the NSD dataset.

**Figure 4.**
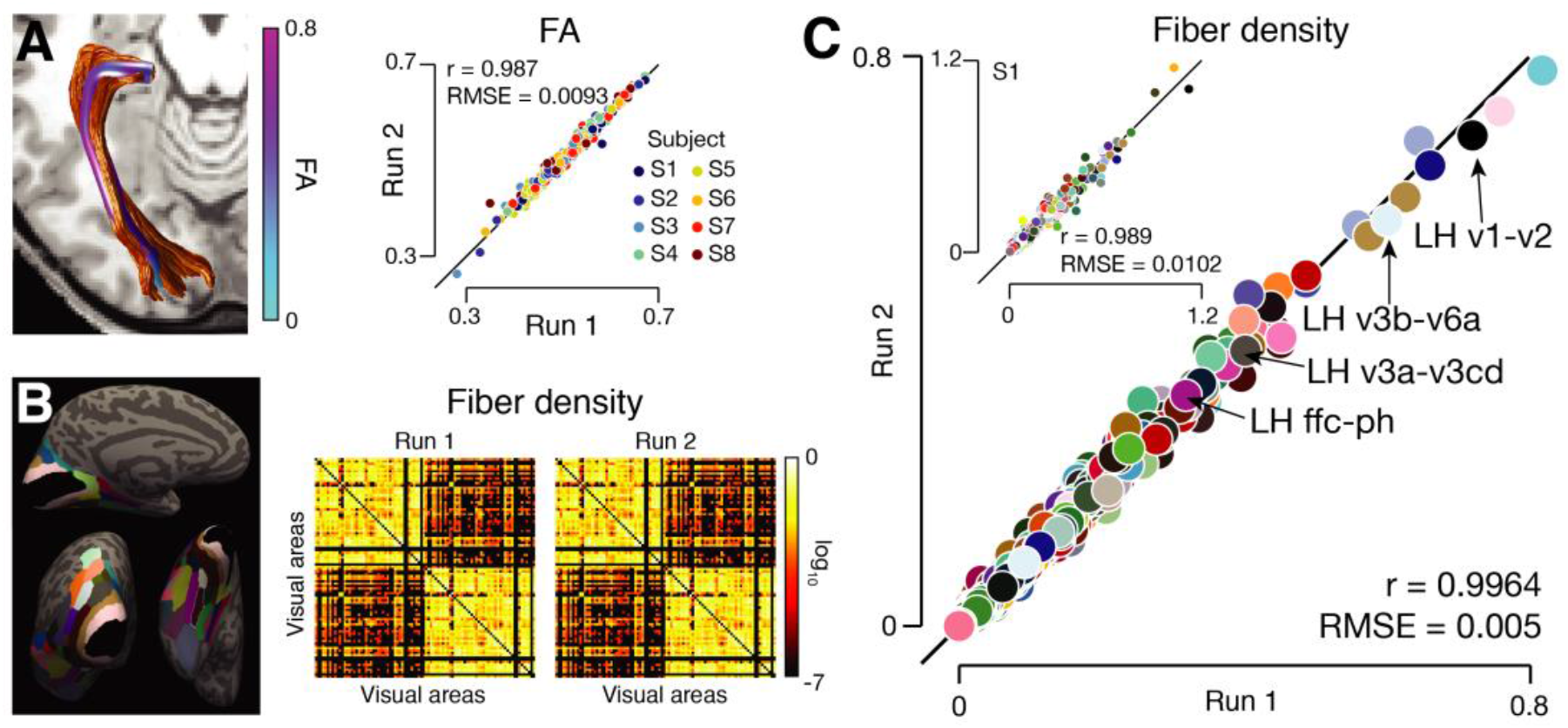
Reliable diffusion derivatives facilitate investigation of white-matter connectivity. *A*, Fractional anisotropy (FA). We show tractography and FA results for the optic radiation identified in subject 7 (left), as well as reliability of FA results for all 61 identified white-matter tracts (right). For other measures, see **Supplementary Figure 6C–E**. *B*, Structural connectivity. Using 43 visual areas × 2 hemispheres = 86 regions from the HCP-MMP1 atlas (Glasser et al., 2016) (left), we construct group-average connectivity matrices indicating the density of fibers connecting pairs of regions (right). *C*, Quantitative summary. Each dot represents fiber density between a pair of regions (as in panel B). Group-average results (main figure) and results for an individual subject (inset) are shown.

### Extensive set of manually defined ROIs

To increase the value of the NSD dataset to the broader community, we performed analysis of the data from the pRF and fLoc experiments and manually defined regions of interest (ROIs) based on the results. The defined ROIs include retinotopic visual areas based on the pRF results (V1, V2, V3, hV4), eccentricity-based regions based on the pRF results (bands between 0°, 0.5°, 1°, 2°, 4°, and beyond), and category-selective regions based on the fLoc results (face-, word-, place-, and body-selective regions). Representative examples illustrating the high quality of the localizer results and associated ROIs are shown in **Supplementary Figure 7**. NSD also includes manual segmentations of the thalamus and the medial temporal lobe. These resources reduce overhead and facilitate scientific analyses of the NSD dataset.

### Novel methods for accurate estimation of single-trial GLM betas

We performed a general linear model (GLM) analysis of the data from the NSD experiment in order to help streamline subsequent analyses of the data. The goal of the GLM was to obtain single-trial betas representing the estimated response of each voxel to each trial conducted. Given the low signal-to-noise ratio of fMRI and the overlap of the hemodynamic response from trial to trial, estimating accurate betas is a challenging endeavor. We thus developed a novel GLM approach consisting of three components. First, we used a library of hemodynamic response functions (HRFs) derived from an initial analysis of the dataset as an efficient and well-regularized method for estimating voxel-specific HRFs (**Figure 5A–C**). Second, we adapted the GLMdenoise technique (Charest et al., 2018; Kay et al., 2013a) to the single-trial GLM framework, thereby enabling the use of data-driven nuisance regressors (**Figure 5D**). Third, to address the challenge posed by highly correlated single-trial regressors, we developed an efficient implementation of ridge regression (Hoerl and Kennard, 1970; Rokem and Kay, 2020) and used this to regularize and improve the accuracy of the betas (**Figure 5E**). To assess the efficacy of these various GLM techniques, we generated three versions of the betas, reflecting increasing sophistication (**Supplementary Figure 8**). Beta version 1 (b1) is the result of simply using a canonical HRF for all voxels. Beta version (b2) is the result of fitting an HRF to each voxel using the library-of-HRFs approach. Beta version (b3) uses the library-of-HRFs approach like b2 but also adds the use of GLMdenoise and ridge regression in an attempt to improve the accuracy of the betas.

**Figure 5.**
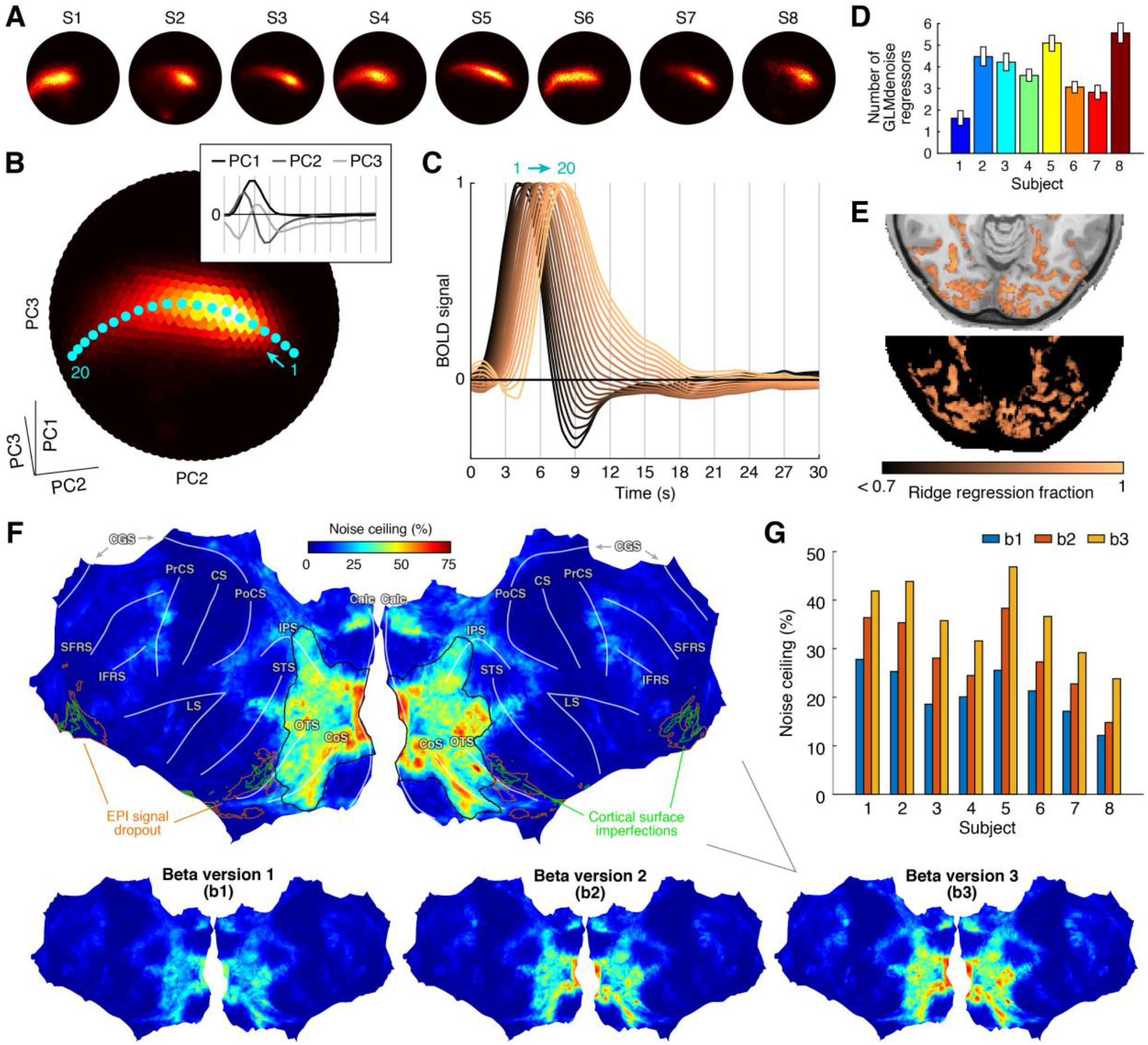
GLM estimation and denoising methods. *A–C*, Library of HRFs. Hemodynamic response functions (HRFs) were estimated within a subspace spanned by 3 principal components (PCs). Distributions of voxel-specific HRFs are shown for individual subjects (panel A) and the group average (panel B). These distributions reside on the unit sphere with coordinate axes corresponding to 3 PC timecourses (see panel B, inset). We defined a series of points on the unit sphere (cyan dots), and the timecourses associated with these points are used as the HRF library (panel C). *D*, GLMdenoise. The average number of GLMdenoise regressors identified for each subject is shown (1.8-mm preparation; error bars indicate bootstrapped 68% confidence intervals). *E*, Ridge regression. Optimal ridge regression fractions are shown for an example scan session (subject 5, nsd10, 1-mm preparation). *F*, Noise ceilings for the case where responses are averaged across 3 trials. Results from individual subjects (nativesurface preparation) were mapped to fsaverage and then averaged. *G*, Performance summary. Each bar indicates the median noise ceiling across vertices in the nsdgeneral ROI.

### Massive scale and quality of NSD responses

To assess the quality of the different beta versions (b1, b2, b3), we calculated noise ceilings for individual voxels. Surface maps of noise ceiling results reveal locations of reliable responses to the NSD stimuli: high noise ceilings are present in occipital cortex and extend into temporal and parietal cortex (**Figure 5F** and **Supplementary Video 10**). Importantly, the maps reveal very large increases in noise ceilings from b1 to b2 to b3, indicating that the additional GLM techniques incorporated into b2 and b3 improve reliability of responses. Detailed quantifications show that these improvements are highly consistent across voxels and subjects (**Figure 5G** and **Supplementary Figure 9**).

The absolute magnitudes of the noise ceilings are noteworthy. For beta version b3, noise ceiling levels in visual cortex are, on average, 36% (calculated by computing the median across the nsdgeneral ROI and then averaging across subjects). This means that a typical visual cortex voxel in the NSD dataset has associated with it a set of 10,000 responses (30,000 trials divided by 3 trials per image = 10,000 images) and a large percentage, 36%, of the variance in these 10,000 values is a signal that is, in theory, predictable. Expressed in terms of Pearson’s correlation (*r*), this is equivalent to a prediction accuracy of *r* = 0.60.

To put these numbers into further perspective, we propose the concept of ‘equivalent trials’ which allows comparison of different datasets that vary in signal-to-noise ratio and trial distribution. The next largest data collection effort that is similar in nature to NSD is BOLD5000 (Chang et al., 2019). Using the same GLM analysis methods on both NSD and BOLD5000, we find that the signal-to-noise ratio per trial is approximately 0.260 for NSD and 0.187 for BOLD5000. Combining these values with the number of trials conducted in each dataset, we estimate that the total size of the NSD dataset is 213,000 trials × (0.260)^2^= 14,399 equivalent trials, whereas the total size of BOLD5000 is 18,870 trials × (0.187)^2^ = 660 equivalent trials. Thus, using the metric of equivalent trials, NSD can be viewed as 14,399/660 = ∼22 times as large as the BOLD5000 dataset. This is a massive increase in power. Note that even if we do not take into account the higher SNR per trial in the NSD dataset, NSD still has substantially more subjects (8 vs. 4), trials per subject (26,625 vs. 4,718, on average), and hours of fMRI per subject (35.5 vs. 13.7, on average) than BOLD5000.

### Successful recovery of retinotopy

Having demonstrated the quality of the NSD data, we now turn to example analyses that illustrate the rich scientific insights that can be derived from the data. As a simple starting example, we fit a voxelwise pRF model that uses local contrast in the NSD images to account for the NSD betas. This simple model is expected to recover spatial tuning in early visual cortex where responses co-vary with stimulus energy (Albrecht and Hamilton, 1982; Boynton et al., 1999). Indeed, in all eight subjects, high-quality maps of angle and eccentricity estimates are obtained in early visual cortex, and these estimates extend all the way to the fovea (**Supplementary Figure 10**). These results provide a check of the validity of the NSD betas, and provide evidence that subjects were able to successfully maintain central fixation during the NSD experiment.

### Reliable and long-term recognition memory effects

The use of a continuous recognition task establishes NSD as one of the largest datasets relevant to human memory. Despite the challenging nature of the task, we find that subjects were able to successfully discriminate old images from new images (average *d*’ across subjects: 1.28, maximum: 1.47, minimum: 0.94). Further, recognition memory remained above chance even at long timescales between repetitions (**Figure 6A**). Specifically, for each session, we calculated a measure of recognition accuracy accounting for guessing (adjusted hit rate: hit rate minus false alarm rate) and binned this measure by the time since last exposure (considering only those trials involving a previously shown image). At the group level, subjects exhibit performance levels greater than chance (adjusted hit rate > 0) in all time bins. At the level of individuals, all subjects show a positive adjusted hit rate in the longest time bin for which data are available for every subject (when binning on a log scale; 7 out of 8 subjects when binning on a linear scale). These results indicate that from its behavioral component alone, NSD is powered to address questions concerning human memory spanning short (seconds) to relatively long (months) timescales.

**Figure 6.**
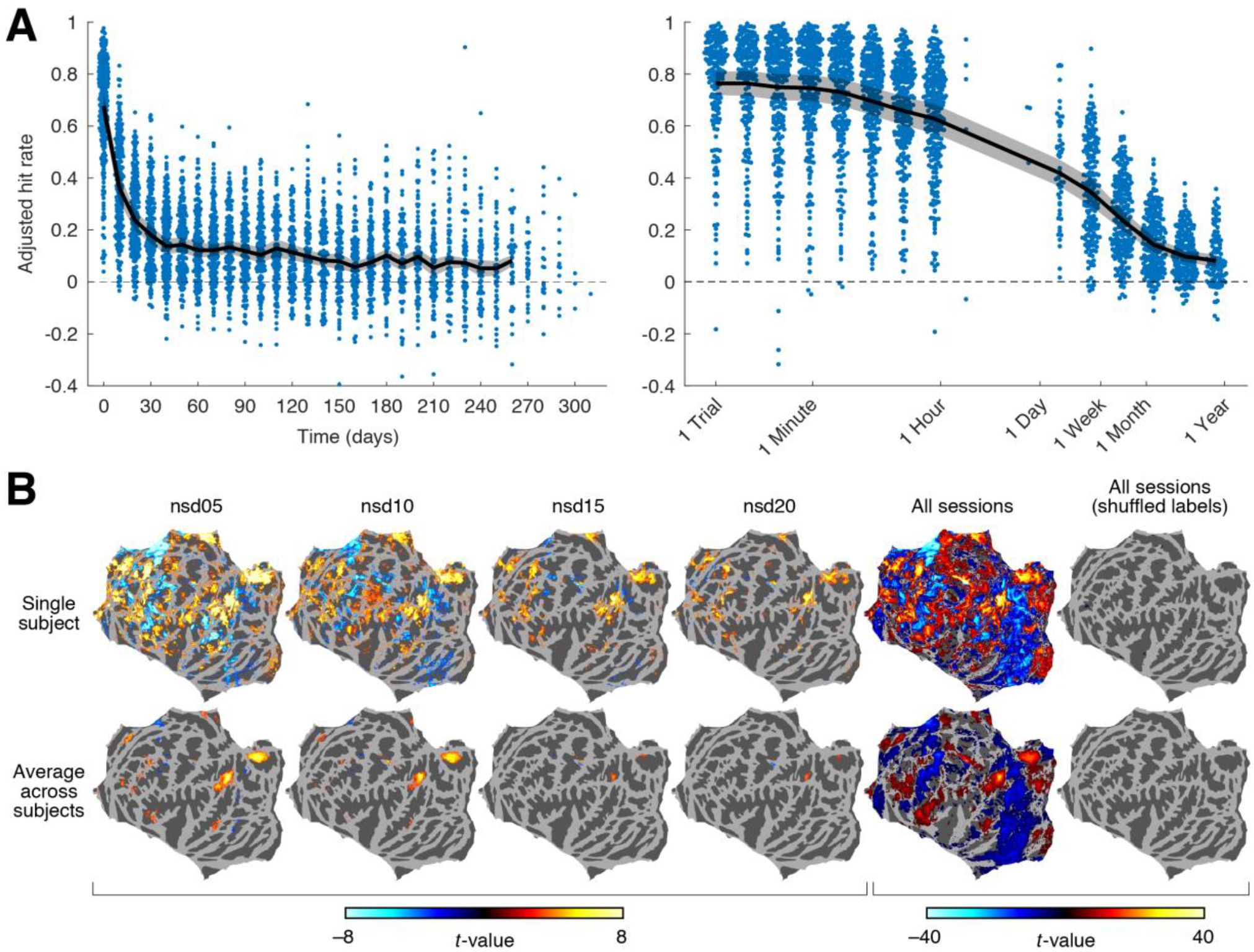
Reliable and long-term recognition memory effects. *A*, Behavioral recognition effects. Adjusted hit rate indicates recognition accuracy accounting for guessing (hit rate minus false alarm rate), and is binned by time between repetitions on a linear (left) or log scale (right). Dashed line indicates chance performance. Each dot in each bin summarizes relevant trials from one scan session. Black line indicates the mean across subjects, with ribbon indicating ± 1 SEM. *B*, Neural recognition effects. We performed two-sample t-tests on NSD betas contrasting ‘hits’ > ‘correct rejections’. All results are shown on a flattened left hemisphere fsaverage surface and thresholded at |*t*| > 3. Tests were performed for trials taken from individual NSD scan sessions (columns 1 through 4) as well as for trials pooled across all NSD scan sessions (column 5). In addition, we perform a control in which trial labels in the pooled analysis are shuffled (column 6). Results for subject 1 (top row) and a simple average of results across subjects (bottom row) are shown.

But what about neural effects? To assess whether recognition effects are present in the fMRI measurements, we performed two-sample *t*-tests contrasting NSD betas observed for hits with NSD betas observed for correct rejections (the so-called ‘old/new effect’; (Wagner et al., 2005)). We find highly consistent old/new effects at the level of individual scan sessions (**Figure 6B, top;** see also **Supplementary Figure 11**). Moreover, these effects occur in expected frontal and parietal regions (Spaniol et al., 2009), and persist at the group level (**Figure 6B, bottom**). The scale and statistical power afforded by the NSD dataset also provides novel insight. Whereas old/new effects are typically studied using group-level analyses, the quality of the NSD dataset reveals highly statistically significant results at the level of individual subjects. Indeed, when pooling trials across all NSD scan sessions, several subjects exhibit statistically significant activity differentiating hits and correct rejections in nearly the entire cerebral cortex (see results for a representative subject in **Figure 6B, top**). Reminiscent of past datasets employing extensive sampling of individuals (Gonzalez-Castillo et al., 2012), the current results suggest that the extent of cortex engaged by basic memory processes is much more widespread than previously appreciated.

### Richness of stimulus sampling

NSD samples a huge variety of natural scenes. To gain insight into the breadth of stimulus sampling available, we constructed representational dissimilarity matrices from the NSD betas and performed *t-* distributed stochastic neighbor embedding (Maaten and Hinton, 2008) (t-SNE) to visualize the underlying representations. We computed t-SNE embeddings in different regions along the ventral visual pathway (**Figure 7A**). By color-coding the resulting embeddings according to animacy attributes (**Figure 7B**), we find that in posterior ventral temporal cortex (pVTC), there is a clear large-scale pattern progressing from images containing people (gray dots; lower left), images containing animals (red dots; middle), and images containing inanimate objects (blue dots; upper right). This progression is not present in early visual areas V1, V2, and V3. In anterior ventral temporal cortex (aVTC), the animacy progression is present to some extent, but a different, more clustered representation emerges that presumably reflects more complex categorical and semantic clusters. Indeed, zooming in on small sections of the t-SNE embedding for aVTC reveals that these clusters contain images with relatively homogeneous semantic content (**Figure 7C**): the blue cluster is dominated by images of round edible objects, while the gray cluster is dominated by images of people interacting with objects. Overall, these results demonstrate the rich inferential power of the NSD dataset and indicate that the dataset can be readily exploited to characterize how visual representations arise in the human brain.

**Figure 7.**
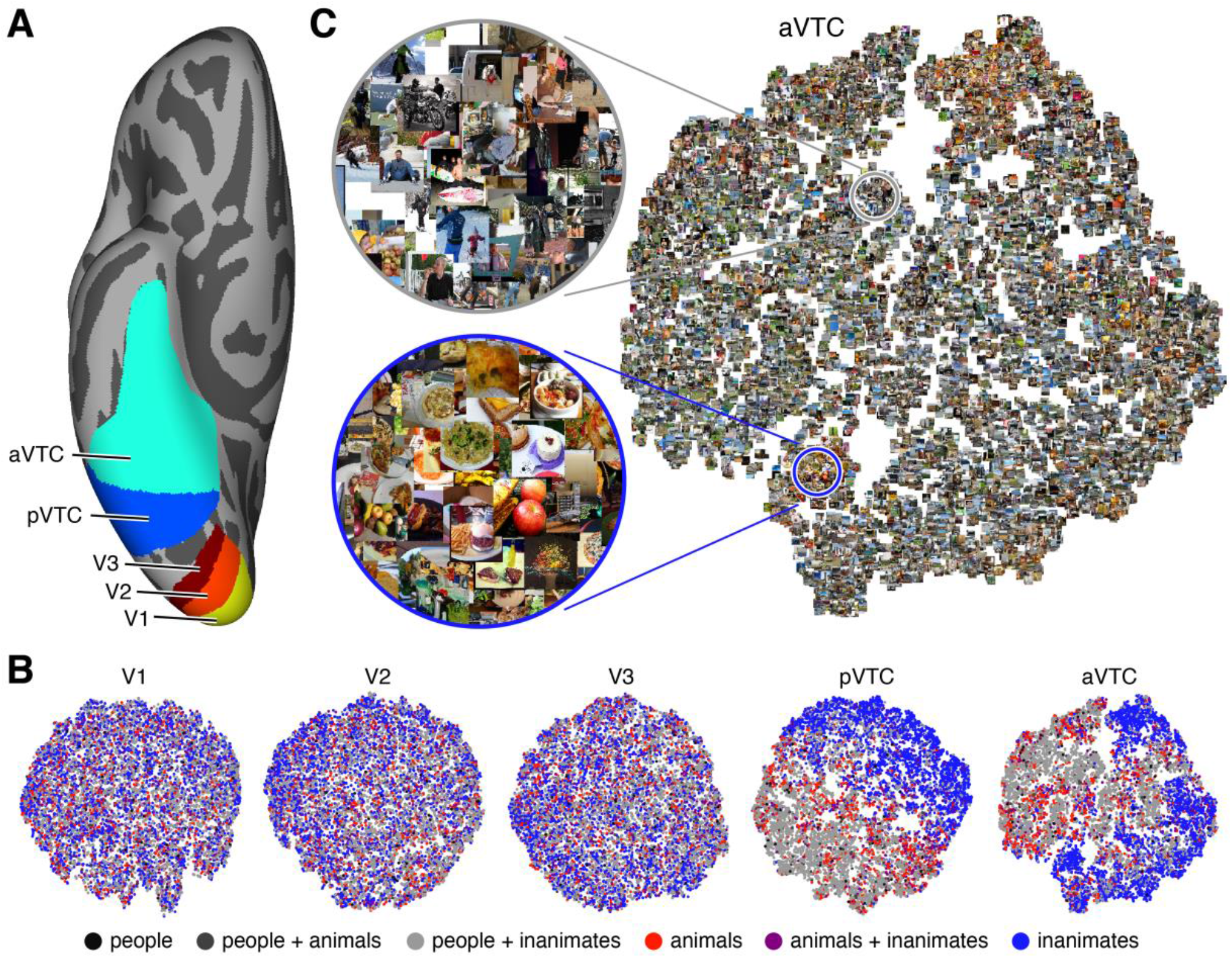
Representational similarity analysis (RSA) reveals transformations of representations along the ventral visual stream. *A*, Illustration of fsaverage ROIs used for the RSA analysis. *B*, t-SNE embedding for each ROI in an example subject (subject 1). Each dot represents a distinct image (total 10,000). Using category labels from the COCO image dataset, we color each dot according to whether the associated image contains particular combinations of people, animals, and inanimates. *C*, t-SNE embedding for anterior ventral temporal cortex with actual images depicted. Insets highlight an inanimate cluster (blue inset) and a cluster of people with inanimate objects (gray inset).

### A brain-optimized neural network model of the visual system

One of the main motivations for NSD was to amass sufficient sampling of brain activity to be able to drive data-hungry machine learning techniques. As an intriguing test case, we specifically investigated whether we could successfully use the scale of NSD to train, from scratch, a deep convolutional neural network (CNN) to accurately predict brain activity (Seeliger et al., 2021). Adopting the framework of encoding models (Naselaris et al., 2011), we took NSD betas from visual areas V1–hV4, divided these data into a training set and validation set, and evaluated how accurately different computational models predict brain responses in the validation set based on the presented image. The primary encoding model of interest is based on a new network we refer to as ‘GNet’, a *brain-optimized* CNN whose parameters are trained using image-response pairings observed in the training set. For comparison, we also evaluated an encoding model based on AlexNet (Krizhevsky et al., 2012), a *task-optimized* CNN whose parameters are pre-trained using explicit labels of objects taken from an image database. AlexNet has been previously shown to provide state-of-the-art performance in modeling visual responses (Güçlü and van Gerven, 2015; St-Yves and Naselaris, 2017). Finally, we included a simple V1-like control model based on oriented Gabor filters (Kay et al., 2008).

Varying the amount of training data provided to the models, we find that the performance of the GNet-based encoding model is relatively poor when only small amounts of training data are available (**Figure 8A, orange arrows**). This is expected since the feature extractors in GNet (see Methods) are not pre-trained and thus require data for tuning. However, when large amounts of training data are available, the GNet model exhibits an impressive increase in performance, achieving approximate parity with the AlexNet-based encoding model (**Figure 8A, blue arrows**). Interestingly, by implementing a variant of GNet that is able to exploit data from multiple subjects, we demonstrate that the GNet model is able to outperform the AlexNet model (two-tailed paired *t*-test across subjects, *p* = 0.013), albeit modestly (**Figure 8A, red arrows**). For additional insight into model performance, we compared voxel-wise performance levels to noise ceiling estimates (**Figure 8B**). Across voxels, prediction accuracy is tightly correlated with the noise ceiling, suggesting that voxel-wise differences in prediction accuracy simply reflect differences in signal-to-noise ratio. In addition, performance levels are close to, but do not reach, the noise ceiling.

**Figure 8.**
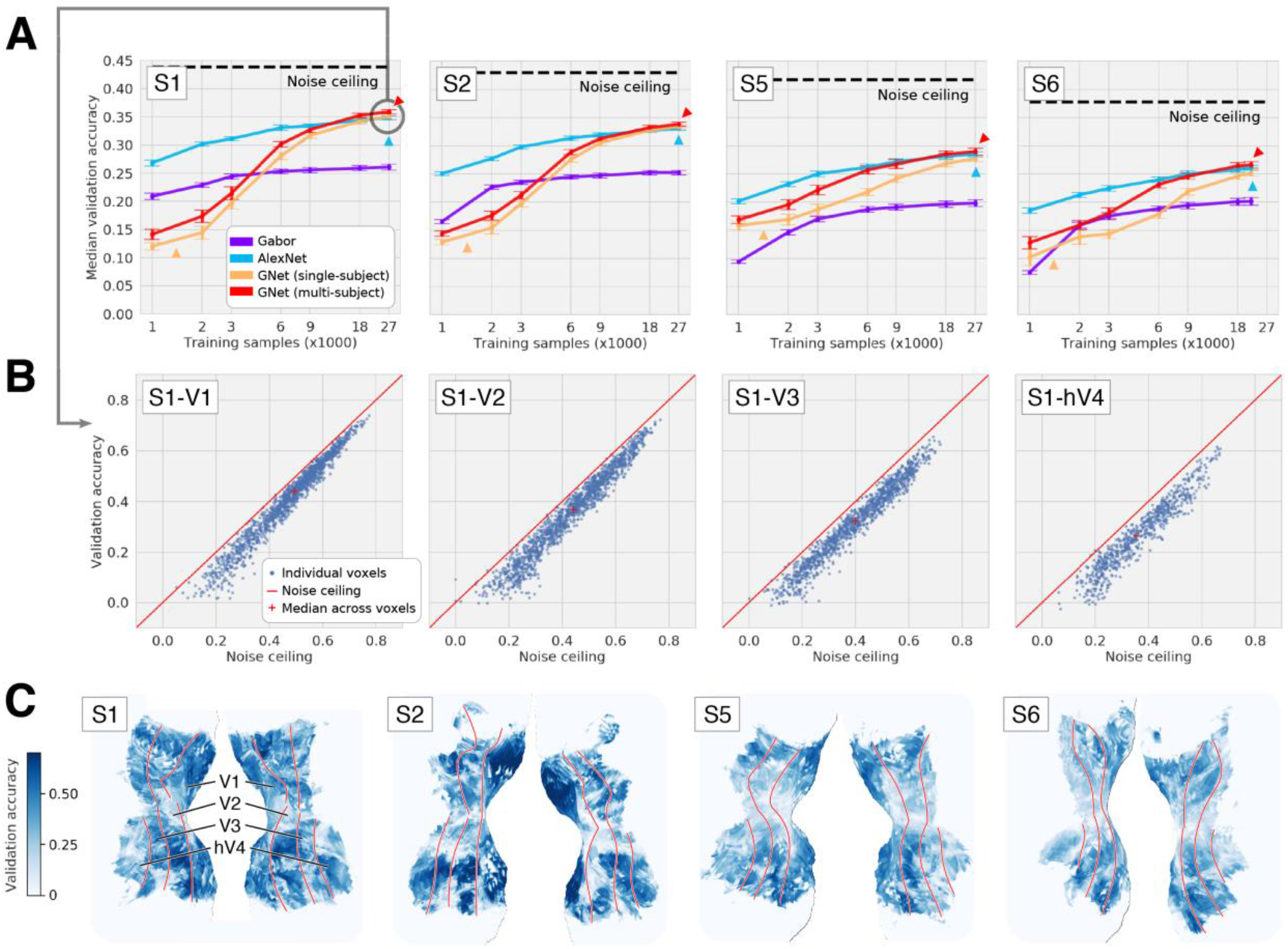
Prediction of brain activity using a brain-optimized neural network. We used encoding models (Naselaris et al., 2011) to predict voxel activity in V1–hV4. NSD betas were divided into a training set and validation set, and the accuracy of different encoding models was quantified as the voxel-wise correlation between model predictions and responses observed in the validation set. *A*, Performance as a function of amount of training data used. Models include an encoding model based on AlexNet which is a task-optimized neural network (blue); encoding models based on GNet which is a brain-optimized neural network trained using data from single subjects (orange) or data from multiple subjects (red); and a V1-like control model based on Gabor filters (purple). Error bars reflect one standard deviation across results obtained from different bootstrap samples of the data. *B*, Detailed view of the performance of the multi-subject GNet model for a representative subject. *C*, Surface maps depicting spatial distribution of validation accuracy for the multi-subject GNet model.

The demonstration that an encoding model based on a brain-optimized CNN (GNet) outperforms an encoding model based on a task-optimized CNN (AlexNet) is significant, as it indicates NSD is large enough to successfully train a complex neural network architecture. Had the NSD dataset been smaller in scale or lower in quality, qualitatively different patterns of model performance would have been obtained (in **Figure 8A**, compare orange arrows reflecting a few thousand trials to red arrows reflecting tens of thousands of trials). The availability of NSD thus opens the door to using brain activity to directly guide the optimization of deep neural networks. Brain-optimized networks (Seeliger et al., 2021) are a useful alternative to task-optimized networks (Khaligh-Razavi and Kriegeskorte, 2014; Yamins et al., 2014), as the narrowly defined tasks that such networks are typically trained to solve do not necessarily respect the diversity of functions supported by the human visual system (Wang et al., 2019) nor necessarily match properties found in biological visual systems (Sinz et al., 2019). In this regard, NSD is a resource that will contribute to a virtuous cycle of competition between models derived from biological and artificial intelligence. In principle, brain-optimized networks might prove to be more effective for solving computer vision tasks (Cox and Dean, 2014; Fong et al., 2018; Toneva and Wehbe, 2019; van de Ven et al., 2020).

## Discussion

In the last several years, there have been a number of large-scale neuroimaging datasets that have been made publicly available for re-use (Aliko et al., 2020; Gordon et al., 2017; Nastase et al., 2020; Taylor et al., 2017; Van Essen et al., 2013). Several distinguishing aspects of the present work set NSD apart from past datasets. One is the *unprecedented scale of the dataset*. NSD shares the motivation of recent ‘deep’ (or ‘precision’) neuroimaging efforts (Bellec and Boyle, 2019; Filevich et al., 2017; Gordon et al., 2017; Pinho et al., 2018; Poldrack et al., 2015; Seeliger et al., 2019) seeking to amass large amounts of data from individual subjects, as opposed to modest amounts of data on a large number of subjects. In this context, NSD is, to our knowledge, the most extensive fMRI data collection effort that has been performed to date. This can be gauged not only in terms of the number of hours of fMRI data acquisition per subject (30–40 hours of data for each of 8 subjects on the core NSD experiment) and the high spatial resolution of the acquired data (1.8 mm), but also the wealth of additional measures beyond the core experiment, including substantial amounts of resting-state and diffusion data, physiological data, and functional localizers. The availability of extensive measures provides the opportunity to build complete models of how individual brains support vision and memory (Naselaris et al., 2021).

A second aspect is the *unusually high quality of the data*. Although quality of neuroimaging data is more complex to assess than quantity, assessment of data quality is essential since MRI data have relatively low sensitivity and are prone to errors and artifacts. In particular, when acquiring massive datasets, there is a risk of accumulating unknown sources of noise and artifact. The work presented in this paper (and in the accompanying files in the data release) guards against this possibility by crafting a customized and highly optimized approach to pre-processing the NSD data and providing comprehensive documentation of the high data quality (see also **Supplementary Discussion**). Several factors likely contributed to the high data quality: these include the use of ultra-high magnetic field strength (7T) which enhances BOLD contrast-to-noise ratio; the screening, training, and incentivization of participants; the detailed inspection and supervision of data processing; and the large network of collaborators who helped guide the design and trajectory of the dataset.

A third aspect of the present work lies in the *novel analysis techniques* developed for improved GLM analysis of fMRI time-series data. These include (i) an efficient and robust method to estimate voxel-specific HRFs, (ii) adaptation of the GLMdenoise technique (Kay et al., 2013a) to a single-trial GLM framework, and (iii) development of ridge regression as an effective method for regularizing single-trial response estimates. These three techniques have been integrated into a toolbox that can be applied to other neuroimaging datasets, and are the subject of a forthcoming paper. An important lesson stemming from our results is that well-executed data collection is important but not the only factor to consider: data preparation methods exert a major influence on the quality of a dataset and hence its scientific value. One can view improvements in data quality as equivalent to increases in data quantity, in the sense that analysis methods that reduce unwanted variability (noise) can be interpreted as increasing the effective amount of data collected (Kay et al., 2013a). Thus, by improving data quality, the methods introduced with NSD are contributing to the massive scale of the dataset.

The NSD dataset has many potential applications in cognitive and computational neuroscience. Given its extensive sampling of natural scenes (70,566 distinct images aggregated across 8 subjects) and high signal-to-noise ratio, the dataset will be useful for investigating a variety of phenomena in low-, mid-, and high-level vision. In addition, the memory component of the NSD experiment provides a unique opportunity to study the neural mechanisms of both short- and long-term memory (ranging from seconds to many months), as well as potential interactions between vision and memory. From a methodological perspective, the repeated scanning of individuals using a consistent experimental manipulation (up to 40 scan sessions of the NSD experiment per subject) provides a unique opportunity for development and evaluation of neuroimaging pipelines. Finally, perhaps the most exciting use of NSD is as a common dataset to bridge the disciplines of cognitive science, neuroscience, and artificial intelligence (Naselaris et al., 2018). As we have shown in the context of deep neural network modeling (see **Figure 8**), there are sufficient data in NSD to successfully drive the training of neural network models with thousands of free parameters. This demonstration exemplifies how NSD—with its large amounts of carefully curated fMRI data collected during a rich cognitive paradigm—enables data-driven approaches towards understanding the complexities of information processing in the brain.

## Author Contributions

E.A. collected the neuroimaging data, coordinated the data collection effort, and performed manual brain segmentations. G.S.-Y. performed neural network analyses. Y.W. performed subject recruitment, assisted with scanning, and prepared eyetracking videos. J.B. assisted in data analysis. L.D. organized and prepared data in BIDS format. B.C. and F.P. analyzed the diffusion data. I.C. performed representational similarity analyses. J.B.H. analyzed the behavioral data. K.K. and T.N. designed the main experiment. J.B.H. and I.C. designed the nsdmeadows and nsdmemory behavioral assessments. K.K. developed analysis methods, analyzed the neuroimaging data, and directed the overall project. K.K., T.N., E.A., B.C., F.P., I.C., and J.B.H. wrote the paper. All authors discussed and edited the manuscript.

## Acknowledgements

We thank E. Aminoff, J. Pyles, M. Tarr, M. Hebart, and C. Baker for advice on experimental design and data collection; J. Power and A. Schapiro for consultation on resting-state and physiological data; V. Carr and R. Olsen for consultation on hippocampal subfield scanning protocols; J. Prince for performing the equivalent-trials analysis on NSD and BOLD5000; A. Grant for assistance with scanner peripherals; F. Gosselin and J. Tardif for contrast sensitivity analysis; B. Klimes-Dougan and K. Cullen for designing the valence/arousal assessment; W. Guo for segmentations of the medial temporal lobe; M. Arcaro, A. Bratch, D. Finzi, A. White, and J. Winawer for assistance with ROI definition; C. Gorgolewski and R. Poldrack for discussion of BIDS and data sharing; R. Cichy, E. Yacoub, K. Grill-Spector, K. Jamison, A. Rokem, A. Huth, N. Kriegeskorte, and J. Winawer for general discussions; and K. Ugurbil for overall project advice. We also thank our NSD collaborators for shaping the trajectory of the project. This work was supported by NSF CRCNS grants IIS-1822683 (to K.K.) and IIS-1822929 (to T.N.).

## Data and code availability statement

Information on the NSD dataset is available at http://naturalscenesdataset.org. The dataset will be made publicly available upon manuscript publication. The data are hosted in the cloud, allowing researchers to exploit high-performance cloud computing to efficiently analyze the dataset. We provide both raw data in BIDS format (Gorgolewski et al., 2016) and prepared data files, along with extensive technical documentation in the NSD Data Manual. To ensure strict validation for an upcoming Algonauts prediction challenge (Cichy et al., 2019), the initial public release will withhold the last three NSD scan sessions from each participant (about 8.4% of the NSD data). We also provide an archive of code used in this paper, as well as utility functions for working with the prepared data. Finally, example scripts demonstrating scientific analyses of the NSD data are available; these scripts may be useful for teaching purposes.

## Methods

### Subject recruitment

The NSD study was advertised to the University of Minnesota community. We sought to recruit right-handed individuals (18–65 years old) with no known cognitive deficits nor color blindness and with normal or corrected-to-normal vision. Those who were interested in participating were contacted for a phone interview to explain the nature of the study and to screen them for eligibility. We discussed the long-term nature of the study, the time commitment it would involve, and the feasibility of traveling to the scanner on a regular basis. We paid attention to the communicativeness of potential participants and their general attitude towards study participation. Selecting participants whom we were confident would provide high-quality data was more important to us than obtaining a random sample of the general population. Based on the phone interviews, we invited 14 people who we felt were strong candidates to participate in an initial 7T fMRI screening session. Of these, 8 were selected to participate in the full NSD experiment.

### Subjects

Eight subjects (two males, six females; age range 19–32) participated in the NSD dataset (subj01– subj08). There were six additional subjects (four males, two females; age range 20–53) who participated in the initial 7T fMRI screening session but not in the remainder of data collection. Subjects were naïve and were not involved in the design nor planning of the NSD dataset. All subjects had normal or corrected-to-normal visual acuity. Informed written consent was obtained from all subjects, and the experimental protocol was approved by the University of Minnesota Institutional Review Board. Additional subject information including height, weight, handedness, and visual acuity was logged and is available online.

Subjects participated in a number of neuroimaging and behavioral data collection sessions (a full breakdown is provided in **Supplementary Figure 2**). Neuroimaging included 3T structural scan sessions and 7T functional scan sessions. The 7T functional scan sessions included an initial screening session termed ‘prffloc’, referring to the population receptive field (pRF) and functional localizer (fLoc) experiments conducted in that session. The 7T sessions also included, for each subject, 30–40 sessions in which the main NSD experiment was conducted (‘nsd01–nsd40’). These sessions are collectively termed the ‘NSD core’. In some of these sessions, resting-state data were acquired before and after the NSD experiment. Finally, the 7T sessions also included two sessions conducted after completion of the NSD core; these sessions, termed ‘nsdsynthetic’ and ‘nsdimagery’, involved measuring responses to synthetic stimuli and cognitive task manipulations (including mental imagery), respectively. The total number of 7T fMRI scan sessions were 43, 43, 35, 33, 43, 35, 43, and 33 for subj01–subj08, respectively. The average number of hours of resting-state fMRI conducted for each subject was 2.0 hours, and the average number of hours of task-based fMRI conducted for each subject was 38.5 hours. Each subject also participated in several behavioral assessments after scanning was complete. These included a variety of behavioral measures (‘nsdpostbehavior’), a final memory test (‘nsdmemory’), and an image-similarity assessment (‘nsdmeadows’).

### MRI data acquisition

MRI data were collected at the Center for Magnetic Resonance Research at the University of Minnesota. Some data were collected using a combination of a 3T Siemens Prisma scanner and a standard Siemens 32-channel RF head coil. Most data were collected using a combination of a 7T Siemens Magnetom passively-shielded scanner and a single-channel-transmit, 32-channel-receive RF head coil (Nova Medical, Wilmington, MA). Illustrations of the different types of MRI data acquired are provided in **Figure 2B**. Below we summarize the scanning protocols (full protocol printouts are available online).

At 3T, we collected a number of anatomical measures (*T*_1_, *T*_2_, diffusion, angiogram). The motivation for collecting data at 3T was to ensure acquisition of *T*_1_ volumes with good gray/white-matter contrast and homogeneity, which is difficult to achieve at ultra-high field (Polimeni et al., 2018). To increase contrast-to-noise ratio and enable the ability to assess reliability, we acquired several repetitions of *T*_1_- and *T*_2_-weighted volumes. For each subject, we collected between 6–10 scans of a whole-brain *T*_1_-weighted MPRAGE sequence (0.8-mm isotropic resolution, *TR* 2400 ms, *TE* 2.22 ms, *TI* 1000 ms, flip angle 8°, bandwidth 220 Hz/pixel, no partial Fourier, in-plane acceleration factor (iPAT) 2, TA 6.6 min/scan) and 2– 3 scans of a whole-brain *T*_2_-weighted SPACE sequence (0.8-mm isotropic resolution, *TR* 3200 ms, *TE* 563 ms, bandwidth 744 Hz/pixel, no partial Fourier, in-plane acceleration factor (iPAT) 2, TA 6.0 min/scan). In addition to *T*_1_ and *T*_2_ data, we also acquired 4 high angular resolution diffusion-weighted spin-echo EPI scans, varying the number of diffusion directions and the phase-encode direction (1.5-mm isotropic resolution, *TR* 3230 ms, *TE* 89.20 ms, flip angle 78°, refocusing flip angle 160°, bandwidth 1700 Hz/pixel, echo spacing 0.69 ms, partial Fourier 6/8, no in-plane acceleration, multiband slice acceleration factor 4, TA 5.6 min/scan for 99 directions, TA 5.7 min/scan for 100 directions). The 4 scans included 99 directions AP (anterior-to-posterior phase-encode direction), 99 directions PA (posterior-to-anterior phase-encode direction), 100 directions AP, and 100 directions PA. Diffusion volumes were acquired at *b*-values of 0, 1,500, or 3,000 s/mm^2^. We also acquired an angiogram using a time-of-flight (TOF) multi-slab 3D sequence (0.39 mm × 0.39 mm × 0.5 mm resolution, *TR* 19.0 ms, *TE* 2.91 ms, flip angle 18°, bandwidth 186 Hz/pixel, phase partial Fourier 6/8, slice partial Fourier 6/8, in-plane acceleration factor (iPAT) 2, *TA* 5.5 min).

At 7T, we collected functional data and associated fieldmaps and a few additional anatomical measures (venogram, high-resolution *T*_2_). Functional data were collected using gradient-echo EPI at 1.8-mm isotropic resolution with whole-brain (including cerebellum) coverage (84 axial slices, slice thickness 1.8 mm, slice gap 0 mm, field-of-view 216 mm (FE) × 216 mm (PE), phase-encode direction anterior-to-posterior, matrix size 120 × 120, *TR* 1600 ms, *TE* 22.0 ms, flip angle 62°, echo spacing 0.66 ms, bandwidth 1736 Hz/pixel, partial Fourier 7/8, in-plane acceleration factor (iPAT) 2, multiband slice acceleration factor 3). The use of moderate spatial resolution capitalizes on the signal-to-noise ratio benefits provided by ultra-high magnetic field strength. At the beginning of each 7T session, we acquired a short test EPI scan and adjusted the gain factor (FFT scale factor) accordingly to ensure large dynamic range while avoiding clipping. Empirical measurements indicate that the acoustic noise caused by the EPI sequence is 112 dBA; assuming a conservative noise reduction estimate of 26 dB for the earplugs that we used, the resulting noise level is 86 dBA, which can be safely endured for approximately 8–16 continuous hours according to guidelines from the National Institute for Occupational Safety and Health (NIOSH) 1998 and Occupational Safety and Health Administration (OSHA) 2009.

In addition to the EPI scans, the 7T sessions also included dual-echo fieldmaps for post-hoc correction of EPI spatial distortion (same overall slice slab as the EPI data, 2.2 mm × 2.2 mm × 3.6 mm resolution, *TR* 510 ms, *TE*_1_ 8.16 ms, *TE*_2_ 9.18 ms, flip angle 40°, bandwidth 301 Hz/pixel, partial Fourier 6/8, *TA* 1.3 min/scan). Fieldmaps were periodically acquired over the course of each scan session to track changes in the magnetic field (details provided below). In one of the 7T sessions held for each subject, we acquired a venogram using a susceptibility-weighted imaging (SWI) 3D sequence (0.5625 mm × 0.5625 mm × 0.6 mm resolution, *TR* 28 ms, *TE* 21 ms, flip angle 17°, bandwidth 120 Hz/pixel, phase partial Fourier 6/8, slice partial Fourier 6/8, in-plane acceleration factor (iPAT) 3, *TA* 10.1 min). In addition, for the purposes of hippocampal segmentation, we acquired in one of the 7T sessions a high-resolution *T*_2_-weighted TSE scan (0.357 mm × 0.357 mm × 1.5 mm resolution, 56 oblique slices oriented perpendicular to the long axis of the hippocampus, field-of-view 160 mm (FE) × 156.4 mm (PE), *TR* 16000 ms, *TE* 53 ms, bandwidth 100 Hz/pixel, no partial Fourier, in-plane acceleration factor (iPAT) 2, turbo factor 15, *TA 4*.*5* min).

In the prffloc 7T fMRI session, the acquisition structure was [F BWLL F BWLL F BWLL F], where F indicates a fieldmap, B indicates a multibar run of the pRF experiment (188 TRs), W indicates a wedgering run of the pRF experiment (188 TRs), and L indicates a run of the fLoc experiment (195 TRs). In the NSD 7T fMRI sessions, the acquisition structure was either [F NNNN F NNNN F NNNN F] or [F RNNNN F NNNN F NNNNR F], where F indicates a fieldmap, N indicates a run of the NSD experiment (188 TRs), and R indicates a resting-state run (188 TRs).

### Stimulus display and scanner peripherals

Ear plugs were used to reduce acoustic noise experienced by the subjects. To minimize head motion, we acquired a headcase (Power et al., 2019) for each of the 8 NSD subjects (Caseforge, Berkeley, CA; http://caseforge.co) and deployed the headcases starting from the second NSD core scan session (nsd02). To ensure maximal subject comfort, only the posterior half of the headcases were used (omitting the anterior half). Standard foam padding was used to mitigate head motion prior to that point (prffloc, nsd01).

Stimuli were presented using a Cambridge Research Systems BOLDscreen 32 LCD monitor positioned at the head of the 7T scanner bed, placed flush against the scanner bore. We chose to use an LCD monitor because it delivers a sharp, high-quality image, in contrast to typical scanner setups involving projectors and backprojection screens. The monitor operated at a resolution of 1920 pixels × 1080 pixels at 120 Hz. The size of the full monitor image was 69.84 cm (width) × 39.29 cm (height). Subjects viewed the monitor via a mirror mounted on the RF coil. The viewing distance was 5 cm from the subjects’ eyes to the mirror + 171.5 cm from the mirror to the monitor image = 176.5 cm total. Measurements of the display spectral power density were obtained using a PR-655 spectroradiometer (Photo Research). The BOLDscreen is designed by the manufacturer to behave as a linear display device, and our measurements confirmed this to be the case.

We determined the maximum square extent visible in both eyes given the constraints of the RF coil to be 8.4° × 8.4° (714 pixels × 714 pixels). Thus, stimuli from the various experiments (e.g., pRF, fLoc, NSD) were adjusted to fill 8.4° of visual angle (details provided below). At the beginning of each scan session, we made an effort to position the monitor in the same location relative to the scanner and to position the subject’s head and RF coil in the same location relative to the scanner. We also used a calibration square (8.4° in size) to determine any incidental horizontal or vertical offsets needed in that session in order for the participant to see the entire square in each eye, unobstructed. Given these efforts, we believe that consistent and high-quality visual stimulation was achieved across scan sessions. Nonetheless, we caution that due to limitations in positioning and/or potential drift over the course of a scan session, some slight occlusion of the corners of the 8.4° × 8.4° square extent may have occurred some of the time.

A Mac Pro computer controlled stimulus presentation using code based on Psychophysics Toolbox 3.0.14 (Brainard, 1997; Pelli, 1997). Behavioral responses were recorded using a button box (Current Designs, Philadelphia, PA). In some scan sessions (nsd21–nsd30, the same sessions in which the primary set of resting-state data were acquired), physiological data were collected using a pulse oximeter and a respiratory belt (stock Siemens equipment). Care was taken to secure the oximeter with tape to the left index finger of the subject and to secure the respiratory belt snugly to the subject’s torso. Visual inspections of the physiological data (available online) suggest that the data are of excellent quality and that more than 90% of the pulse data and more than 95% of the respiratory data are usable. These physiological data are valuable for identifying potential contaminants of resting-state fMRI signals (Lynch et al., 2020). Physiological data were carefully synchronized with the fMRI data and cropped, but are not further described in this paper.

In several scan sessions (see **Supplementary Figure 2** for details), eyetracking was performed using an EyeLink 1000 system (SR Research, Mississauga, Ontario, Canada) combined with a custom infrared illuminator mounted on the RF coil. Eyetracking was performed for the left eye, and eyetracking data were obtained at 2000 Hz using the Pupil-CR centroid mode. We caution that the eyetracking data are variable in quality, as achieving sufficient pupil contrast was often difficult. For complementary information, we also captured video recordings of the eyetracker computer display (see **Figure 2B**) using a cell phone secured to a mount. Eyetracking data and video recordings were carefully synchronized with the fMRI data and cropped accordingly (*analysis_eyetracking*.*m*), but are not further described in this paper. Note that it may be possible to infer eye gaze from fMRI signal intensities in and around the eyeballs (Frey et al., 2020; Son et al., 2020).

### Day-to-day acquisition procedures

Participants were scanned roughly once a week, with attempts to keep a regular weekly scan time. At the beginning of each session (starting at approximately nsd07), participants were asked to rate on a five-point scale how well they slept the night before, their mood, how hungry they were, and their stress level. We also asked whether they had had caffeine in the past three hours. At the end of each scan session, participants were asked to rate how comfortable they were during the session and to provide any general feedback they had about the session. These various measures, as well as any technical issues that arose during the session, were logged onto a spreadsheet (available online).

In the first several scan sessions, we emphasized the importance of fixation and performed simple tests prior to scanning in which we watched the subject’s eyes while they attempted to fixate and while they deliberately broke fixation. This was done to help subjects understand what good fixation feels like. In every scan session, we reminded subjects about the importance of fixation and about the correct mapping between buttons and responses.

During data collection, we monitored aspects of data quality including overall image quality, head motion, quality of physiological data, and behavioral performance. Between functional runs, we checked in with the subject to assess their energy level, enthusiasm, and compliance. If we noticed any substantial drops in response rate, we politely notified the subject and offered short breaks before continuing.

To promote subject engagement and retention, participants were given the opportunity to earn monetary bonuses that gradually increased in size over the course of the NSD study. These bonuses were contingent on achieving certain performance levels on data quality metrics such as head motion and response rate. Information regarding performance was supplied to participants in the form of a continually updated “leaderboard” figure. We found that this figure greatly helped to motivate participants.

### The NSD experiment

#### Basic design

In the NSD experiment, participants performed a long-term continuous recognition task while viewing a large number of color natural scenes. We chose this recognition task because it engages and challenges the observer and is unbiased with respect to the specific content of the images (unlike other tasks such as animacy judgment). In addition, it infuses the experiment with a rich memory dimension that is likely of interest to memory researchers. A total of 73,000 distinct images were prepared. We intended that the 8 NSD subjects would each view 10,000 distinct images presented 3 times each over the course of 40 scan sessions. We designated a special set of 1,000 images as shared images that would be seen by all subjects (‘shared1000’); all other images would be mutually exclusive across subjects. The distribution of the 3 presentations of each image was tightly controlled, and subjects were naïve as to both the number and distribution of the presentations.

Images were presented using a 3-s ON / 1-s OFF trial structure (**Figure 1A**). In informal piloting, we found that this pacing made the recognition task feasible and not overly taxing. In addition, we reasoned that the relatively long stimulus duration would increase neural activity and that the rapidity of the design would allow more trials to be collected and thereby increase overall experimental power. Finally, we speculated that the 3/1 trial structure would yield a pleasant experience for participants, at least compared to slow event-related designs where most experimental time is spent viewing a blank screen.

#### Image preparation

The NSD stimuli are prepared as a single brick of RGB images with dimensionality 425 pixels × 425 pixels 3 RGB channels × 73,000 images and unsigned 8-bit integer format.

Images were taken from Microsoft’s Common Objects in Context (COCO) image database (Lin et al., 2014). COCO images are photographs harvested from online repositories; each image is supplemented by a rich set of annotations (e.g., boundary polygons around objects, natural language captions, body-pose estimates). We used NSD images in the 2017 train/val split (Lin et al., 2014), and restricted selection to the subset of images for which pixel-level annotations of “stuff” (Caesar et al., 2018) (e.g., sky, land, wall, road) in addition to “things” (e.g., car, skateboard, hat) were available.

We selected only images whose smaller dimension (height or width) was at least 425 pixels. Where necessary, we squared image dimensions by cropping out pixels along the largest dimension. For example, if the original image was 425 × 585, we cropped away 160 pixels from the larger dimension, resulting in an image that is 425 × 425. The median number of pixels cropped per image was 160. After cropping, images were downsampled, if needed, to 425 × 425.

Cropping an image can change the way the viewer interprets it. We refer to this effect of cropping as “semantic loss”. In order to be able to take full advantage of the rich annotations available for the COCO images, we attempted to minimized semantic loss when cropping images. For landscape-oriented images, we selected between a center, left, or right crop. For portrait-oriented images, we selected between a center, top, or bottom crop (finer grids of cropping options had little effect on results). Selection of crops were carefully performed based on quantitative analysis and visual inspection (details provided in online materials).

In addition to screening to minimize semantic loss, we implemented a screening procedure to remove duplicate images. Some of the COCO images are extremely similar to each other, differing only by a post-processing operation (i.e., grayscaling or sharpening) or by a few frames in a motion-capture sequence. To remove these near-duplicates, we downsampled all images to 40 × 40 and then computed the correlation of grayscale pixel intensities between all image pairs. We manually inspected the image pairs with the 500 highest correlation values. Of these, 38 image pairs were observed to be near duplicates. We randomly selected another image from the COCO dataset to replace one image in each near-duplicate pair. Finally, we screened captions for all images for indications of violent or salacious content. No images were deemed too offensive to include in the experiment.

The distribution of “thing” categories across the final images selected for NSD was nearly identical to distribution in the full COCO dataset. As a result, the “person” category was over-represented; however, with a few exceptions, all 80 COCO object categories are displayed in at least 100 images to each subject. Note that images tend to depict more than one category, so that a given object category frequently appeared in the same image with other categories. For each subject’s images, at least 90% of the images contain 2 or more of the 80 COCO categories.

#### Distribution of image presentations

We determined the ordering of the 10,000 images × 3 trials = 30,000 trials in advance and kept the ordering fixed across subjects. The idea is that these 10,000 images are actually treated as slots into which different NSD images are inserted. We designated the first 1,000 slots as corresponding to the special shared1000 images; the remaining 9,000 slots were filled with unique images for each subject. Note that because the trial ordering and repetition structure are identical across subjects, the difficulty of the recognition task is comparable across subjects (up to the fact that some images might be more difficult to remember than others).

We controlled the distribution of image presentations in order to prevent the recognition task from becoming too difficult (and risking loss of subject morale). In the procedure, we conceptualized the task of determining the trial ordering as equivalent to placing image presentations on a circle that would eventually be cut and unraveled. To determine presentation times, we created a circular probability distribution by mixing a von Mises distribution and a uniform distribution (**Supplementary Figure 1A**). Using random draws from the resulting distribution (positioning the distribution at a random location on the circle for each image), we determined 3 presentation times for each of the 10,000 images. After completing the placement of all 30,000 trials, we then cut the circle, unraveled it into a linear sequence of image presentations, and divided this sequence into 40 consecutive segments corresponding to the 40 NSD scan sessions (750 trials per session).

We chose specific parameters for the probability distribution: we used a von Mises distribution with concentration parameter of 3^6^ and a mixing ratio of 60% and 40% for the von Mises and uniform distributions, respectively. This choice of parameters yields appealing properties. First, the distribution is relatively narrow (see **Supplementary Figure 1A**) and therefore ensures that there will be many trials involving an image that has been presented in the recent past (thus, making the trials easy), while still allowing the probing of more distant memory events. Second, there is minimal “burn-in” time at the beginning of the experiment: even in the first scan session, there is still a substantial number of trials involving old images (see **Supplementary Figure 1B, blue line**). Third, there is minimal “dead” time at the end of the experiment: even in the last scan session, there is still a substantial number of trials involving new images (see **Supplementary Figure 1B, blue line**).

To provide a sense of the overall experimental design, we computed basic statistics on each NSD scan session. For a typical session, the total number of distinct images shown once, twice, and all three times within that session is 437, 106, and 34, respectively (these numbers reflect the mean across scan sessions, rounding to the nearest integer).

#### Trial and run design

Each trial lasted 4 s, and consisted of the presentation of an image for 3 s, followed by a 1-s gap. A total of 75 trials were conducted in a run; thus, each run lasted 300 s. The first 3 trials (12 s) and last 4 trials (16 s) were blank trials. The remaining 68 trials were divided into 63 stimulus trials and 5 blank trials. The blank trials were randomly positioned in each run such that the minimum and maximum continuous number of stimulus trials was 9 trials (36 s) and 14 trials (56 s), respectively (see **Figure 1B**). For even-numbered runs, the 63^rd^ stimulus trial was designated to be a blank trial. A total of 12 NSD runs were collected in one NSD session, yielding a total of (63 + 62) × 6 = 750 stimulus trials. Moreover, this design was repeated for all 40 NSD sessions: 750 stimulus trials × 40 sessions = 30,000 stimulus trials. The temporal ordering of stimulus and blank trials was generated once and kept fixed across subjects.

#### Stimulus presentation and task

Since the BOLDscreen is calibrated to behave as a linear display device, we used a squaring luminance response when presenting the NSD experiment in order to simulate the typical viewing of digital images. At time of presentation, the prepared NSD images (425 pixels × 425 pixels) were resized to fill 8.4° × 8.4° (714 pixels × 714 pixels) using linear interpolation. Throughout each run (including blank trials), a small semi-transparent red fixation dot (0.2° × 0.2°; 50% opacity) was present at the center of the stimuli (**Figure 1A**). Stimuli were shown against a gray background with RGB value (127,127,127).

Subjects were instructed to fixate the central dot and to press button 1 using the index finger of their right hand if the presented image was new, i.e. the image had never been presented before, or button 2 using the middle finger of their right hand if the presented image was old, i.e. the image is identical to one that had been presented before, either in the current scan session or any previous scan session. Subjects were additionally instructed to continue to fixate and wait for the next image in the event of blank trials.

Before the start of the NSD experiment, we showed the subjects a version of the experiment involving cartoon images in order for them to become familiarized with the feel and timing of the task. During the NSD experiment, minimal feedback was provided to the subjects regarding their performance on the recognition task. Participants were blind to the precise details of the NSD experiment (e.g., total number of images, total number of presentations per image). Participants were informed only about their response rate (fraction of trials on which they successfully made a response) and a vague “performance metric” which, unbeknownst to them, quantified their percent correct for easy trials (trials that involved the presentation of an image that had occurred earlier in the same scan session). A postmortem held after the completion of the NSD experiment revealed the nature of the design (details below).

#### Details on experiment timing

Stimulus presentation was locked to the refresh rate of the BOLDscreen monitor. Empirical measurements confirmed that the monitor refresh rate was nearly exactly 120 Hz: duration of runs were highly reliable, ranging from 299.95–299.98 s. To compensate for the slight offset from 300 s, the fMRI data were pre-processed to achieve a sampling rate of 0.999878 s (high-resolution preparation) or 0.999878 s × (4/3) = 1.333171 s (standard-resolution preparation). For brevity, we refer to these numbers as 1.000 s and 1.333 s. Experimental runs were started by a trigger issued by the MR scanner. Due to input polling and monitor refresh, there was slight variability in the delay between trigger detection and the presentation of the first stimulus frame, ranging from 3–22 ms. We did not attempt to compensate for this delay.

#### Acquisition

Due to constraints on subject availability (including unplanned out-of-town absences in the summer of 2019) and due to constraints on scanner availability (the 7T scanner was decommissioned in November 2019), we did not complete the full NSD experiment for every participant. Fortunately, we were able to collect a sizable amount of data: 40, 40, 32, 30, 40, 32, 40, and 30 NSD sessions for subj01–subj08, respectively. In these collected data, each subject viewed 9,209–10,000 distinct images and participated in 22,500–30,000 trials. Aggregated across subjects, the total number of distinct images shown was 70,566, and the total number of trials was 213,000.

#### Postmortem

After completion of the final memory test (details below), participants filled out a post-NSD questionnaire. This questionnaire probed topics such as strategies used for performing the NSD task and estimates for the number of images viewed and the number of image repetitions. After filling out this questionnaire, the design of the NSD experiment was then revealed to the participants.

### Other experiments

#### Population receptive field (pRF) experiment

We adapted the experiment used in the Human Connectome Project (HCP) 7T Retinotopy Dataset (Benson et al., 2018). Stimuli consisted of slowly moving apertures filled with a dynamic colorful texture (see **Figure 2A**). Apertures and textures were updated at a rate of 15 Hz. Two run types were used. The first, termed ‘multibar’, involves bars sweeping in multiple directions (same as RETBAR in the HCP 7T Retinotopy Dataset). The second, termed ‘wedgering’, involves a combination of rotating wedges and expanding and contracting rings. Both run types included blank periods.

For consistency with the NSD experiment, stimuli were resized to fill a circular region with diameter 8.4°. Each run lasted 300 s (exact empirical timings were highly accurate and ranged between 299.95–300.00 s). Throughout stimulus presentation, a small semi-transparent dot (0.2° × 0.2°) was present at the center of the stimuli. The color of the central dot switched randomly to one of three colors (black, white, or red) every 1–5 s. Subjects were instructed to maintain fixation on the dot and to press a button whenever the color of the dot changed. To further aid fixation, a semi-transparent fixation grid was superimposed on the stimuli and was present throughout the experiment (Schira et al., 2009). A total of 6 runs (3 multibar, 3 wedgering) were collected in the first 7T fMRI session (prffloc).

#### Functional localizer (fLoc) experiment

This experiment was developed by the Grill-Spector lab (Stigliani et al., 2015) (stimuli and presentation code available at http://vpnl.stanford.edu/fLoc/). The experiment consisted of the presentation of grayscale images depicting different stimulus categories (see **Figure 2A**). There were 10 categories, grouped into 5 stimulus domains: characters (word, number), bodies (body, limb), faces (adult, child), places (corridor, house), and objects (car, instrument). Stimuli were presented on a scrambled background (different backgrounds for different stimuli). Stimuli were presented in 4-s trials. In a trial, 8 images from a given category were sequentially presented (image duration 0.5 s). Each run included 6 presentations of each of the 10 categories as well as blank trials (also of 4-s duration).

For consistency with the NSD experiment, stimuli were resized to fill a square region filling 8.4° × 8.4° of visual extent. Each run lasted 300 s (exact empirical timings were highly accurate and ranged between 300.000–300.002 s). Throughout stimulus presentation, a small red fixation dot was present at the center of the stimuli. Subjects were instructed to maintain fixation on the dot and to press a button whenever they noticed an image in which only the background was present (“oddball” task). A total of 6 runs were collected in the first 7T fMRI session (prffloc).

#### Resting-state experiment

Stimuli consisted of a white fixation cross (0.5° × 0.5°) on a gray background (see **Figure 2A**). Each resting-state run lasted 300 s. In the second resting-state run held within a given scan session, the fixation cross turned red after 12 s had elapsed and remained red for 4 s before returning to white.

Resting-state data were acquired in several NSD core scan sessions: nsd21–nsd38 for subj01 and subj05, and nsd21–nsd30 for all other subjects. Thus, a total of 100 or 180 minutes of resting-state data were acquired for each subject; this large amount of data is appealing as it enables stable estimates of resting-state correlations (Laumann et al., 2015). In each session, one resting-state run was acquired at the beginning of the session (prior to the NSD runs) and another resting-state run was acquired at the end of the session (after the NSD runs).

In the first resting-state run, subjects were instructed to stay awake and fixate the cross but otherwise rest. In the second resting-state run, subjects were additionally instructed to inhale deeply when the fixation cross turned red. This instructed breath was designed to aid analysis of the physiological data collected concomitantly with the resting-state data. Prior to each resting-state run, subjects were asked to report their current sleepiness level using the Stanford Sleepiness Scale (Shahid et al., 2012) (1–7 where 1 is most active and 7 is most sleepy). After each resting-state run, subjects were asked to report their sleepiness level during the run that had just completed.

After the last scan session involving resting-state data, participants filled out a post-resting-state questionnaire. This questionnaire queried what the participants were doing during the resting-state runs and whether they thought about the images from the NSD experiment.

#### Synthetic stimuli experiment (nsdsynthetic)

After completion of the NSD experiment, we conducted an additional 7T fMRI scan session in which responses were measured to a variety of carefully controlled synthetic (non-naturalistic) stimuli while the subject performed either a fixation task or a one-back task. These data will be described and released in a forthcoming manuscript.

#### Visual imagery experiment (nsdimagery)

After completion of the nsdsynthetic experiment, we conducted an additional 7T fMRI scan session in which responses were measured while participants engaged in visual imagery and other cognitive tasks. These data will be described and released in a forthcoming manuscript.

#### Additional behavioral measures (nsdpostbehavior, nsdmemory, nsdmeadows)

A number of behavioral assessments were conducted after completion of the NSD experiment. Several of these were relatively brief, and included the following (nsdpostbehavior): open-ended questions regarding language ability; the Vividness of Visual Imagery Questionnaire (Marks, 1973); the Test of Word Reading Efficiency (Torgesen et al., 2012), including both Sight Word Efficiency and Phonemic Decoding Efficiency; the Cambridge Memory Test for Faces (Duchaine and Nakayama, 2006); ultra-fast measurement of contrast sensitivity (Tardif et al., 2016); and an assessment of chromatic sensitivity (participants adjusted intensities of red, green, and blue channels on the BOLDscreen display until minimal luminance flicker was perceived).

We also conducted a final memory test in which we collected various memory-related measures regarding the images shown to the subjects during the NSD experiment (nsdmemory). These data will be described and released in a forthcoming manuscript.

Finally, using the web-based Meadows platform (http://meadows-research.com), we conducted an assessment of how the NSD subjects perceive and interpret the NSD images (nsdmeadows). First, we selected a small set of images that maximally span semantic space. This was done by isolating the 515 images (out of the shared 1,000 NSD images) that were viewed all 3 times by all 8 NSD subjects during the NSD experiment; computing shifted inverse frequency sentence embeddings for the sentence captions provided by the COCO dataset (Arora et al., 2017); and using a greedy approach to determine the subset of 100 images that maximize the average distance between each image’s embedding and its closest neighbor. We then asked participants to perform a Multiple Arrangements Task (Kriegeskorte and Mur, 2012) in which they arrange using a drag-and-drop interface the 100 images within a white circular arena according to the similarity of their content. Using an adaptive procedure, subsequent arrangements were conducted using subsets of the images in order to maximize information gain. This was done until 45 minutes had elapsed. Using a similar interface on Meadows, participants then provided valence and arousal ratings for the 100 images as well as 3 additional images pulled from the 515 images. Ratings were performed separately for valence and arousal, and were accomplished by freely arranging using a drag-and-drop interface the images (delivered in small batches) along a one-dimensional axis ranging from low to high. This assessment took about 15 minutes.

### Overview of data analysis

We designed custom analysis strategies to maximize the quality of derived measures from the NSD data. A number of methods are based on recent work in the lab where further details can be found (Kay et al., 2020, 2019). Data analysis and visualization were performed using custom code in MATLAB and Python as well as tools from various packages such as FreeSurfer, SPM, FSL, ANTs (Avants et al., 2011), and ITK-SNAP (Yushkevich et al., 2006). An archive of code used is provided online, and specific code files are referenced in the text below.

The analysis of the NSD data can be divided into three components: (i) pre-processing of the anatomical, diffusion, and functional data, (ii) time-series analysis of the fMRI data to estimate trial-wise betas, and (iii) further analyses of the trial-wise betas to answer specific scientific questions. The first two components produce the so-called ‘prepared data’ that are generally useful to the community, whereas the third component refers to analyses performed for the purposes of this paper (pRF estimation, univariate memory analysis, representational similarity analysis, end-to-end neural-network training).

The pre-processing approach that we designed for the NSD dataset prioritizes accuracy and preservation of information (e.g. avoiding spatial smoothing). We avoid “baking in” unnecessary assumptions (e.g. aggressively removing signal fluctuations without careful assessment of validity), and care is taken to manually inspect each pre-processing step to ensure satisfactory results. While we believe our pre-processing is general and likely suitable for most downstream uses of the data, the raw data are also available for those who wish to explore other pre-processing approaches such as fmriprep (Esteban et al., 2019). We note several aspects of the NSD dataset that may render the dataset challenging from a pre-processing standpoint: the relatively high spatial resolution of the fMRI data (1.8 mm) places higher demands on spatial accuracy, the ultra-high field strength (7T) used for the fMRI data yields higher levels of EPI spatial distortion compared to lower field strengths, and the emphasis on many repeated scans of individuals heightens the importance of achieving consistent imaging results across scan sessions.

### Pre-processing of anatomical data

#### T_1_ and T_2_ volumes

*T*_*1*_- and *T*_*2*_-weighted volumes were corrected for gradient nonlinearities using a custom Python script (https://github.com/Washington-University/gradunwarp) and the proprietary Siemens gradient coefficient file retrieved from the scanner. The multiple *T*_*1*_ and *T*_*2*_ volumes acquired for a given subject were then co-registered (*preprocess_nsd_structuralalignment*.*m*). This was accomplished by first co-registering the *T*_*1*_ volumes to each other (rigid-body transformation; correlation metric; the first *T*_*1*_ volume serving as the target) and then by co-registering the *T*_*2*_ volumes to the *T*_*1*_ volumes (rigid-body transformation; mutual information metric; the prepared *T*_*1*_ data serving as the target). In the estimation of registration parameters, a manually defined 3D ellipse was used to focus the cost metric on brain tissue. Individual volumes were manually inspected and rejected if substantial image artifacts were visible (only the 2nd and 4th *T*_*1*_ volumes for subject 8 were rejected). The final *T*_*1*_ and *T*_*2*_ data were created by performing cubic interpolation at a resolution of 0.5 mm. Results for the multiple acquired volumes were averaged (within modality) to increase contrast-to-noise ratio. Finally, the 0.5-mm volumes were resampled to create alternative 0.8-mm and 1.0-mm versions.

#### SWI volume

We co-registered the SWI volume (magnitude component only; corrected for gradient nonlinearities) to the prepared 1.0-mm EPI volume (*preprocess_nsd_SWI*.*m*). To compensate for the acquisition being performed on different scanners, we used a slightly flexible nonlinear warp as implemented in ANTs 2.1.0 (BSplineSyN with parameters [0.1, 400, 0, 3]). The final SWI volume was prepared in the subject-native anatomical space by performing B-spline interpolation at a resolution of 0.5 mm. The resulting volume was then resampled to create alternative 0.8-mm and 1.0-mm versions.

#### TOF volume

We co-registered the TOF volume to the prepared 1.0-mm *T*_*1*_ volume (*preprocess_nsd_TOF*.*m*). This was accomplished using a slightly flexible nonlinear warp as implemented in ANTs 2.1.0 (BSplineSynN with parameters [0.1, 200, 0, 3]). To aid estimation of registration parameters, a temporary version of the TOF volume was used in which extremely bright pixels were dampened. The final TOF volume was prepared in the subject-native anatomical space by performing B-spline interpolation at a resolution of 0.5 mm. The resulting volume was then resampled to create alternative 0.8-mm and 1.0-mm versions.

#### High-resolution T_2_ volume

We co-registered the high-resolution *T*_*2*_ volume to the prepared 0.5-mm *T*_*2*_ volume (*external_mtl*.*m*). Given that these volumes were acquired on different scanners, we evaluated several strategies for achieving accurate co-registration. We obtained the best results by performing a simple linear co-registration (affine transformation; correlation metric) in combination with a rectangular box that focused the cost metric on regions of interest in the medial temporal lobe. The estimated registration was subsequently used to map labels defined on the high-resolution *T*_*2*_ volume to the subject-native anatomical space.

#### De-identification

We mapped a liberal brain mask defined in MNI space to the subject-native anatomical space, and then used the result to mask and thus de-identify the anatomical volumes (*preprocess_nsd_applybrainmask*.*m*).

#### FreeSurfer processing

The prepared 0.8-mm *T*_*1*_ volume was processed using FreeSurfer version 6.0.0 with the *-hires* option (*analysis_freesurfer*.*m*). Manual edits of tissue segmentation (labeling voxels as gray matter, white matter, or cerebrospinal fluid) were performed for each of the eight subjects to optimize the accuracy of the cortical surface representations generated by FreeSurfer. The prepared 0.8-mm *T*_*2*_ volume was used to inform manual segmentation decisions, but was not explicitly used in the FreeSurfer processing. We also manually marked surface imperfections that remained even after manual edits; these are labeled in the surface inspections (**Supplementary Video 2**) and are largely confined to a few difficult regions located in the inferior aspects of the temporal and frontal lobes (see **Figure 5F**).

Several additional FreeSurfer processing steps were performed. Using *mris_expand*, we generated cortical surfaces positioned at 25%, 50%, and 75% of the distance between the pial surface and the boundary between gray and white matter. These surfaces are useful for creating surface representations of the fMRI data. Multiple surfaces at different gray-matter depths were created given the relatively high spatial resolution of the fMRI data (1.8-mm acquisition). We also generated several flattened surface representations: for each hemisphere in each subject, we created a flattened version of the entire cortical sheet (using manually defined cuts) as well as flattened versions of cortical patches covering ventral temporal cortex and early visual cortex (patches were determined automatically based on a set of cortical patches defined on fsaverage). Finally, in line with the ‘surface voxels’ visualization technique (Kay et al., 2019), we sampled 1-, 2-, and 3-mm volumetric test patterns onto surface vertices using nearest-neighbor interpolation (*analysis_surfacevoxels*.*m*). The test patterns are useful for understanding the impact of cortical curvature on surface visualizations.

### Pre-processing of the diffusion data

Code scripts used to analyze the diffusion data are accessible via the hyperlinks indicated below, which refer to brainlife.io apps (http://brainlife.io) (Avesani et al., 2019).

The prepared 0.8-mm *T*_1_-weighted volume for each subject was segmented into different tissue types using MRTrix3 (Tournier et al., 2019) ([brainlife.app.239](https://brainlife.io/app/5db234008aeeee3329f25d9c)). The gray- and white-matter interface mask was subsequently used as a seed mask for white-matter tractography. For network generation, the HCP-MMP cortical parcellation (Glasser et al., 2016) was mapped to subject-native surfaces and then to the volumetric Freesurfer segmentation (ribbon.mgz) for each subject ([brainlife.app.23](https://brainlife.io/app/5aeb34f2f446980028b15ef0)).

The raw data from the four diffusion scans (99 AP, 99 PA, 100 AP, 100 PA) were corrected for gradient nonlinearities, concatenated, and then pre-processed following a published protocol (Ades-Aron et al., 2018). Specifically, diffusion volumes were denoised and cleaned with respect to Gibbs ringing using MRTrix3 before being corrected for susceptibility, motion, and eddy distortions using FSL’s *topup* and *eddy* functions (brainlife.app.287). Following these corrections, the diffusion volumes were bias-corrected and had background noise removed using MRTrix3. Finally, the diffusion volumes were co-registered to the 0.8-mm *T*_1_-weighted anatomical volume using FSL’s *epi_reg* (rigid-body transformation, boundary-based registration), and then resliced to 0.8-mm isotropic voxels. The diffusion data were organized into two runs: data from the 99 AP and 99 PA scans constitute ‘Run 1’ and data from the 100 AP and 100 PA scans constitute ‘Run 2’ (brainlife.app.371). To assess data quality, we calculated signal-to-noise ratio in the corpus callosum using workflow provided by Dipy (Garyfallidis et al., 2014) ([brainlife.app.120](https://doi.org/10.25663/bl.app.120)).

Following pre-processing, brain masks were generated using Dipy’s *median_otsu* ([brainlife.app.70](https://brainlife.io/app/5a5100c306b9e8003f1ee684)). This mask was used in subsequent model fitting and tractography. Multiple models of myelinated microstructural organization were fit to the diffusion data from each run. This included the diffusion tensor (DTI) model (Pierpaoli et al., 1996), diffusion kurtosis (DKI) model (Jensen and Helpern, 2010), and the neurite orientation dispersion and density imaging (Daducci et al., 2015; Zhang et al., 2012) (NODDI) models ([brainlife.app.9](https://brainlife.io/app/5ae9d732f446980028b15ec2),brainlife.app.365). The NODDI model was fit twice for each run: once for white-matter tract microstructure using an intrinsic free diffusivity parameter (*d*_*‖*_) of 1.7 × 10^−3^ mm^2^/s, and once for cortical microstructure using *d*_*‖*_ = 1.1 × 10^−3^ mm^2^/s, following previously described procedures (Fukutomi et al., 2018). The constrained spherical deconvolution (Tournier et al., 2007) (CSD) model was fit for 4 spherical harmonic orders (*L*_max_ = 2, 4, 6, 8) using MRTrix3 (brainlife.app.238). The fiber orientation distribution functions for *L*_max_ = 6 and *L*_max_ = 8 were subsequently used to guide anatomically-constrained probabilistic tractography (Smith et al., 2012) using MRTrix3 (brainlife.app.297). A total of 3 million streamlines across *L*_max_ = 6 and *L*_max_ = 8 for each run were generated, using a step size of 0.2 mm, minimum length of 25 mm, maximum length of 250 mm, and a maximum angle of curvature of 35°. Finally, structural connectivity matrices representing fiber density were generated using a combination of SIFT2 (Smith et al., 2015) and MRTrix3 (brainlife.app.394). Fiber density was quantified as the number of streamlines connecting two regions divided by the average volume of the two regions.

Following model fitting and tractography, 61 major white-matter tracts were segmented for each run using a customized version of the white matter query language (Bullock et al., 2019) ([brainlife.app.188](https://brainlife.io/app/5cc73ef44ed9df00317f6288)). Then, using Vistasoft (https://github.com/vistalab/vistasoft), outlier streamlines were removed ([brainlife.app.195](https://brainlife.io/app/5cc9eca04b5e4502275edba6)) and tract profiles (each tract sampled with 200 nodes) were generated for DTI, DKI, and NODDI measures ([brainlife.app.361](https://brainlife.io/app/5ed02b780a8ed88a57482c92)). Finally, these measures were mapped to the cortical surface following previously published procedures (Fukutomi et al., 2018) using FreeSurfer and Connectome Workbench (https://github.com/Washington-University/workbench) ([brainlife.app.379](https://brainlife.io/app/5eea8548d39ceb4a30a090eb)).

### Pre-processing of functional data

#### Overall strategy

We implemented a pre-processing approach that aimed to preserve as much spatial and temporal detail as possible. In short, the fMRI data were pre-processed by performing one temporal resampling to correct for slice time differences and one spatial resampling to correct for head motion within and across scan sessions, EPI distortion, and gradient nonlinearities. This produced volumetric fMRI time-series data in subject-native space for each NSD subject. The functional data were pre-processed independently of the anatomical data; this was done intentionally in order to avoid dependence of the pre-processed functional data on choices such as how to co-register the functional and anatomical data and how to map the functional data to cortical surface representations. Also, to minimize the risk of inaccurate or unwanted assumptions, we did not include any temporal filtering (e.g. detrending, confound regression, censoring). Pre-processing results were carefully visually inspected to ensure quality control. There were a few anomalous cases, such as acquisition being split across two different scan sessions; special modifications were made to the pre-processing to accurately compensate for these occurrences (see online notes for details).

#### First-stage pre-processing

Given the fMRI data acquired in a scan session, a series of steps were performed (*preprocess_nsd*.*m, preprocessfmri*.*m*):

1. *Temporal resampling*. A cubic interpolation of each voxel’s time-series data in each run was performed. This interpolation corrected differences in slice acquisition times (as determined from the DICOM header) and also upsampled the data (in the same step) to either 1.333 s (standard-resolution preparation) or 1.000 s (high-resolution preparation). Data were prepared such that the first time-series data point coincides with the acquisition time of the first slice acquired in the first volume of each run. The upsampling exploits the benefits of temporal jitter between the acquisition and the experiment and synchronizes the time-series data to convenient multiples of the experiment trial structure (Kay et al., 2020). For example, in the standard-resolution preparation, there are 3 time points for each 4-s trial in the NSD experiment.
2. *Fieldmap preparation*. The multiple fieldmaps acquired in the scan session (3.6-mm slices) were upsampled using nearest-neighbor interpolation to match the slice resolution of the fMRI data (1.8-mm slices). The fieldmaps were then phase-unwrapped using the FSL utility *prelude* and regularized by performing 3D local linear regression using an Epanechnikov kernel with radius 5 mm. Values in the magnitude component of the fieldmaps were used to regularize the fieldmaps and the regression in order to improve robustness of the field estimates. Finally, the fieldmaps were linearly interpolated over time, producing an estimate of the field for each fMRI volume acquired.
3. *Spatial undistortion*. The temporally resampled volumes from Step 1 were undistorted based on the field estimates from Step 2 using the standard unwarping method (Jezzard, 2012). Undistorted volumes were generated using cubic interpolation.
4. *Motion estimation*. The undistorted volumes from Step 3 were used to estimate rigid-body motion parameters using the SPM5 utility *spm_realign* (the first fMRI volume in the scan session served as the reference). A manually defined 3D ellipse was used to focus the cost metric on brain regions unaffected by gross susceptibility effects. Note that the estimated motion parameters reflect temporally upsampled data and should be interpreted accordingly (e.g. when assessing framewise displacement). Also, note that the motion parameters may reflect apparent image motion due to respiration-induced *B*_0_ fluctuations (Gratton et al., 2020); this was particularly apparent in subject 3.
5. *Spatial resampling*. A single cubic interpolation was performed on the temporally resampled volumes from Step 1 in order to correct for the combined effects of head motion and spatial distortion.

#### Gradient nonlinearity correction and session registration

Given the results of the first-stage pre-processing, we computed the mean fMRI volume and corrected this volume for gradient nonlinearities (*preprocess_nsd_epigradunwarp*.*m*). We then co-registered this gradient-corrected volume to the gradient-corrected volume from the first NSD scan session (affine transformation, correlation metric). Thus, the first NSD scan session determined the target space for preparing fMRI data from different scan sessions (*preprocess_nsd_epialignment*.*m*).

#### Second-stage pre-processing

We repeated the pre-processing steps (Steps 1–5 above) but with the final spatial resampling step incorporating the effects of the gradient nonlinearity correction and the session registration (*preprocess_nsd_secondstage*.*m*). In this way, a single cubic interpolation is used to compensate for the effects of head motion, spatial distortion, gradient nonlinearities, and session registration. For this final interpolation step, we used either a 1.8-mm grid (standard-resolution preparation) or a 1.0-mm grid (high-resolution preparation). The latter approach intentionally upsamples the data in order to exploit the benefits of small head displacements and preserve as much spatial detail as possible (Kang et al., 2007; Kay et al., 2019). To minimize storage requirements, the interpolations were performed within a 3D box that was just large enough to cover the brain of each subject.

To facilitate assessment of *T*_2_* effects, we created a bias-corrected version of the mean EPI volume (*analysis_biascorrection*.*m*). For each preparation, we took the mean EPI volume and fit a 5th-degree 3D polynomial, considering only voxels labeled as cortical or cerebellar gray matter in the FreeSurfer aseg file. The fitted volume (‘coilbias’) was then divided from the mean EPI volume, producing the bias-corrected volume (‘bc’).

#### Final outputs

The final result of pre-processing was volumetric fMRI time-series data in subject-native space. Two versions were generated: the standard-resolution version was prepared at a spatial resolution of 1.8 mm and a temporal resolution of 1.333 s, whereas the high-resolution version was prepared at a spatial resolution of 1 mm and a temporal resolution of 1.000 s.

### Mapping between spaces

We performed several analyses related to mapping data between different spaces:

- *Mapping between functional and anatomical spaces*. We co-registered the mean fMRI volume (1-mm preparation; mean of first five NSD sessions) to the prepared 1.0-mm *T*_*2*_ volume (*preprocess_nsd_functionaltostructuralalignment*.*m*). To compensate for acquisition on different scanners, we used a slightly flexible nonlinear warp as implemented in ANTs 2.1.0 (BSplineSyN with parameters [0.1, 400, 0, 3]). A small amount of nonlinearity was necessary to achieve accurate co-registration (see inspections provided online).
- *Changing resolutions in anatomical space*. For resampling data to different anatomical resolutions (0.5-, 0.8-, or 1.0-mm), we used an ideal Fourier filter (10th-order low-pass Butterworth filter) followed by cubic interpolation (*changevolumeres*.*m*).
- *Mapping to and from fsaverage*. We calculated the indexing information that maps subject-native surfaces to and from fsaverage using nearest-neighbor interpolation in the spherical space defined by FreeSurfer. Visual inspections confirm the quality of the folding-based alignment achieved by FreeSurfer (**Supplementary Video 3**).
- *Mapping to and from MNI*. Using FSL’s utility *fnirt*, we co-registered the subject-native prepared 1.0-mm *T*_*1*_ volume to the MNI152 *T*_*1*_ 1-mm template provided by FSL (*preprocess_nsd_MNIandbrainmask*.*m*). Visual inspections confirm the quality of the registration (**Supplementary Video 4**).
- *Converting surface data to volumetric format*. We implemented a method that, for a given target anatomical volume with resolution *R* mm, allows each surface vertex to contribute a triangular (linear) kernel of size +/– *R* mm and then calculates a weighted average of data values at the position of each voxel in the volume (*cvnmapsurfacetovolume_helper*.*m*).

Based on the results of the above analyses, we calculated coordinate transformations that indicate how to map between the functional spaces, the anatomical spaces, the subject-native surfaces, fsaverage, and MNI space (*preprocess_nsd_calculatetransformations*.*m*). Finally, we created a lightweight utility (*nsd_mapdata*.*{m,py}*) that uses the coordinate transformations to map user-supplied data from one space to another space under a given interpolation scheme (nearest-neighbor, linear, cubic, winner-take-all). For example, data in subject-native functional space can be mapped to MNI space using a single interpolation of the functional data. We used this utility to perform a number of mundane but useful transformations, such as generating versions of the anatomical volumes that are matched to the functional volumes (*analysis_transforms*.*m*). Label data (e.g. ROI labels) were transformed by performing a separate interpolation for each label and then a winner-take-all operation.

### Data quality metrics

Several data quality metrics were calculated (*export_runmetrics*.*m*) and summarized in **Figures 1D** and **2D**. Temporal signal-to-noise ratio (tSNR) was computed from raw fMRI volumes (no pre-processing) by first detrending the time-series data from each voxel (quadratic polynomial fit) and then dividing the mean signal intensity by the standard deviation of signal intensity values (*autoqc_fmri*.*m*). We calculated the median tSNR across voxels within a simple brain mask (mean volume thresholded at 1/10th of the 99th percentile of values) and then computed the median across runs. Head motion was quantified by calculating framewise displacement (FD) (Power et al., 2012) based on motion parameter estimates (1.8-mm preparation). We calculated the mean FD across volumes in a run and then computed the median across runs. BOLD response was quantified by calculating the percentage of variance explained by a simple ON-OFF GLM model (1.8-mm preparation). We calculated the median variance explained across voxels within the nsdgeneral ROI and then computed the median across runs. Response rate was quantified by calculating the percentage of trials for which the subject pressed a button and then computing the mean across runs. Percent correct for easy trials was quantified by calculating the percentage of trials for which the subject gave the correct response, considering only those trials for which the presented image had been previously presented in the same scan session.

To identify EPI signal dropout regions (*export_signaldropout*.*m*), we divided the *T*_*2*_ volume (resampled to match the EPI data) by the mean EPI volume (1-mm preparation). The resulting volume is useful as it indicates which voxels have high signal intensity in the *T*_*2*_ but are corrupted by signal dropout in the EPI. We mapped the volume to the cortical surface (cubic interpolation; mean across depth), transformed the result to fsaverage, and then used a data-driven threshold to mark atypically high values. Vertices marked in at least four of the eight subjects are indicated in **Figure 5F**. To visualize surface imperfections, we took the voxels that were marked in the 0.8-mm anatomical space (during the inspection of surface quality), smoothed this binary volume with a 3D Gaussian with full-width-half-max of 2 mm, mapped the result to the cortical surface (cubic interpolation; max across depth), and then transformed the result to fsaverage. Vertices exceeding 0.01 in at least one of the eight subjects are indicated in **Figure 5F**.

### Regions of interest (ROIs)

#### Volume-based subject-native ROIs

The *thalamus* ROI collection consists of ROIs related to the lateral geniculate nucleus, pulvinar, and superior collicus (*external_subcortical*.*m*). Manual labeling of these ROIs was performed by an expert (M. Arcaro) based on the *T*_*1*_ and *T*_*2*_ anatomical volumes obtained for each subject as well as functional results obtained in prior studies (Arcaro et al., 2015), projected from MNI space to the native space of each subject. Labels were defined in 0.5-mm anatomical space and were resampled to create alternative 0.8-mm and 1.0-mm versions. To provide labels in functional space, the 0.8-mm anatomical volume was mapped to the 1.0-mm functional space, and the 1.0-mm anatomical volume was mapped to the 1.8-mm functional space.

The *MTL* ROI collection consists of ROIs related to the hippocampus and surrounding brain regions in the medial temporal lobe (*external_mtl*.*m*). Manual labeling of these ROIs was performed by an expert (W. Guo) based on the high-resolution *T*_*2*_ volume obtained for each subject, following a published protocol (Berron et al., 2017). Labels were defined on the raw high-resolution volume, and were mapped to 0.5-mm anatomical space using the previously determined affine transformation. Note that the resulting labels have some amount of jaggedness due to the anisotropy of the voxels in the high-resolution *T*_*2*_ volume. Alternative versions of the labels were created in the same way as described for the *thalamus* labels.

#### Surface-based subject-native ROIs

Results of the pRF experiment were used to define *prf-visualrois*, a collection of ROIs consisting of the dorsal and ventral subdivisions of V1, V2, and V3, and area hV4 (*analysis_drawrois_prf\**.*m*). These ROIs were manually drawn on cortical surfaces by experts (K. Kay, J. Winawer) based on pRF angle and eccentricity estimates, following common practices (Winawer and Witthoft, 2017). The ROIs extended to the fovea (0° eccentricity) but were restricted to the extent of cortex stimulated by the pRF experiment. The pRF results were also used to define *prf-eccrois*, a collection of ROIs consisting of concentric regions with increasing eccentricity coverage (0.5°, 1°, 2°, 4°, >4°). Labeled regions were confined to the same cortical extent labeled in *prf-visualrois*.

Results of the fLoc experiment were used to define several collections of category-selective ROIs (*analysis_drawrois\**.*m*). These ROIs were manually drawn on cortical surfaces by experts (K. Kay, A. White, A. Bratch) based on a combination of anatomical location and stimulus selectivity *t*-values obtained from the fLoc experiment, following general procedures used in prior studies (Gomez et al., 2018; Kay and Yeatman, 2017; Stigliani et al., 2015). For each ROI collection (*floc-bodies, floc-faces, floc-places, floc-words*), several ROIs exhibiting preference for the associated category were defined (e.g., *floc-faces* was based on *t*-values for the contrast of faces > non-faces). ROIs were defined by drawing a polygon around a given patch of cortex and then restricting the ROI to vertices within the polygon that satisfy *t* > 0. This liberal criterion was used to provide maximum flexibility (the user can simply restrict the ROI further to a specific desired threshold).

#### Surface-based atlas ROIs

To help summarize results in this paper, we defined *nsdgeneral*, an ROI in occipital cortex reflecting regions generally responsive in the NSD experiment (*analysis_drawnsdgeneral*.*m*). This ROI was drawn on fsaverage based on group-average results for variance explained by the b3 version of the GLM. The ROI is shown in **Figure 5F**.

To provide anatomical reference, we defined *corticalsulc*, a collection of ROIs consisting of major sulci and gyri, and *streams*, a collection of ROIs reflecting large-scale divisions of visual cortex (early, midventral, midlateral, midparietal, ventral, lateral, parietal). These ROI collections were manually drawn on fsaverage. Annotations reflecting several of the corticalsulc ROIs are shown in **Figure 5F**. Abbreviations: CGS = cingulate sulcus, PrCS = precentral sulcus, CS = central sulcus, PoCS = postcentral sulcus, SFRS = superior frontal sulcus, IFRS = inferior frontal sulcus, LS = lateral sulcus, Calc = calcarine sulcus, OTS = occipitotemporal sulcus, CoS = collateral sulcus, STS = superior temporal sulcus, IPS = intraparietal sulcus.

For convenience, the NSD dataset also includes a few publicly available atlases. These include *Kastner2015* (Wang et al., 2015), an atlas of visual topography, and *HCP_MMP1* (Glasser et al., 2016), a whole-brain cortical parcellation based on multimodal measures. Both atlases were prepared in fsaverage and converted as described below.

#### Conversion of surface-based ROIs

A number of conversions were performed to prepare volumetric versions of surface-based ROIs (*analysis_surfaceroistovolume*.*m*). ROIs defined on fsaverage were mapped to subject-native surfaces using nearest-neighbor interpolation. ROIs defined on subject-native surfaces were mapped to 0.8-mm anatomical space by assigning labels to the 3 depth-dependent surfaces and then performing weighted linear conversion (as described earlier). The 0.8-mm volume was then mapped to the 1.0-mm and 1.8-mm functional spaces.

### Analysis of the pRF experiment

The pre-processed fMRI data from the pRF experiment were analyzed using the Compressive Spatial Summation model (Kay et al., 2013b) as implemented in analyzePRF (*analysis_prf*.*m*). First, the time-series data from the three repetitions of each run type (multibar, wedgering) were averaged. Stimulus apertures indicating the position of the texture were prepared at a resolution of 200 pixels × 200 pixels. We then used analyzePRF (http://cvnlab.net/analyzePRF/) to estimate pRF parameters for each voxel (canonical hemodynamic response function; seedmode 2). Results were mapped to the cortical surface by performing linear interpolation on the volumes (1-mm preparation) and then averaging across cortical depth. To quantify behavioral performance, we calculated, for each run, (*A* – *B*)/*C* × 100, where *A* indicates the number of successful detections of color changes (button pressed within 1 s of a color change), *B* indicates the number of extraneous button presses, and *C* indicates the total number of color changes. Performance averaged across the six runs ranged between 93.5–98.9% for the eight NSD subjects.

### Analysis of the fLoc experiment

The pre-processed fMRI data from the fLoc experiment were analyzed using GLMdenoise (Charest et al., 2018; Kay et al., 2013a), a data-driven denoising method that derives estimates of correlated noise from the data and incorporates these estimates as nuisance regressors in a general linear model (GLM) analysis of the data (*analysis_floc*.*m*). We coded the 10 stimulus categories using a “condition-split” strategy (Kay et al., 2019) in which the trials associated with each category were split into separate conditions in each run. We used six condition-splits, thereby producing six response estimates (betas) for each category. After fitting the GLM, *t*-values were computed from the GLM betas in order to quantify selectivity for different categories and domains (e.g., selectivity for faces was quantified by perfoming a *t*-test that contrasts adult and child faces vs. all other categories). Results were mapped to the cortical surface by performing linear interpolation on the volumes (1-mm preparation) and then averaging across cortical depth. To quantify behavioral performance, we calculated the hit rate for each run (button pressed within 1 s of an oddball image). Performance averaged across the six runs ranged between 90.8–97.5% for the eight NSD subjects.

### Rankings from the 7T fMRI screening session

Six quality measures (pRF BOLD, fLoc BOLD, pRF behavior, fLoc behavior, raw motion, detrended motion) were computed for each of the 14 subjects who participated in the screening session. BOLD quality was quantified as the percentage of voxels for which variance explained by modeling the fMRI time-series data (either pRF model fitting or GLM model fitting) exceeded 20%. Behavior quality was quantified as described above. Motion was quantified by calculating the median voxel displacement relative to the reference volume used for motion correction, computing the median of this quantity across volumes, and then computing the mean across runs. This motion quantification was performed using raw motion parameter estimates (thereby providing a measure of global head displacement over the course of the session) as well as using motion parameter estimates that are linearly detrended within each run (thereby providing a measure of within-run head instability). Each of the six measures was linearly scaled to span the range 1–5 where 1 corresponds to the worst performance and 5 corresponds to the best performance observed across subjects. Finally, the normalized measures were averaged to produce an overall ranking for each subject, as depicted in **Figure 2C**.

### Analysis of behavioral data from the NSD experiment

The behavioral data from the NSD experiment were lightly reformatted for the convenience of subsequent analyses (*analyzebehavior_nsd*.*m*). We first checked whether the subject had accidentally positioned their fingers on incorrect buttons on the button box, and compensated for this if necessary. (In a few instances, we deliberately instructed subjects to use alternative buttons due to hardware malfunction of the button box.) We then recorded, for each stimulus trial, several quantities including time of image presentation, whether the image presented was new or old, whether the response was correct or incorrect, and the reaction time. Button responses were extracted from a time window extending 250– 4250 ms after image onset. In the case of multiple buttons pressed during a trial, we scored the final button pressed, excluding any redundant presses of that button (subjects sometimes repeated button presses for good measure).

### GLM analysis of the NSD experiment

#### Overview of approach

We performed a GLM analysis of the pre-processed time-series data from the NSD experiment. To maximize flexibility for subsequent analyses, the GLM approach was designed to provide estimates of BOLD response amplitudes (‘betas’) for single trials. Due to low signal-to-noise ratio, single-trial estimation in fMRI is challenging. We therefore developed several analysis components in order to optimize the quality of single-trial betas. These components are packaged into a tool called *GLMsingle*, and is the subject of a forthcoming manuscript where additional details and discussion can be found.

The first analysis component of GLMsingle is the use of a library of hemodynamic response functions (HRFs) whereby the best-fitting HRF from the library is chosen for each voxel. This simple approach for compensating for differences in hemodynamic timecourses across voxels (Handwerker et al., 2012) has several appealing features: it is efficient and can be executed with little computational cost (and hence can accommodate the massive scale of NSD); and it invariably provides well-regularized HRF estimates. The second analysis component is an adaptation of GLMdenoise to a single-trial GLM framework. GLMdenoise (Kay et al., 2013a) is a technique in which data-derived nuisance regressors are identified and used to remove noise from—and therefore improve the accuracy of—beta estimates. The third component is an application of ridge regression (Hoerl and Kennard, 1970) as a method for dampening the noise inflation caused by correlated single-trial GLM predictors. To determine the optimal level of regularization for each voxel, we make use of a recently developed efficient re-parameterization of ridge regression called ‘fractional ridge regression’ (Rokem and Kay, 2020).

#### Derivation of the library of HRFs

To generate a library of HRFs that accurately capture empirically occurring timecourse variation, we performed an initial analysis of data from the first NSD core session (nsd01). The first step was to create a comprehensive summary of observed timecourses (*hrf_derivecanonicalpcs*.*m*). The time-series data from each subject’s nsd01 session was fit using a finite impulse response model (0–30 s) where all of the stimulus trials are treated as instances of a single experimental condition (this simplification is necessary to make estimation feasible). We identified voxels for which model variance explained (*R*^2^) was greater than 10%, and from these voxels randomly drew 20,000 voxels (with replacement). Pooling across subjects, timecourse estimates from the resulting 160,000 voxels were subjected to singular value decomposition to determine the top 3 principal components (shown in **Figure 5B, inset**). To fine-tune timecourse estimates, we re-fit the time-series data from the nsd01 session using these 3 principal components as the basis (as opposed to the finite impulse response basis). Finally, adopting the visualization approach of the Temporal Decomposition Method (Kay et al., 2020), we projected voxel timecourse estimates onto the unit sphere (using the same voxel selection criterion of *R*^2^ > 10%), and constructed a 2D histogram for each subject (shown in **Figure 5A**).

The second step was to define a set of timecourses that span the observed timecourse variation (*hrf_constructmanifold*.*m*). To do this, we converted the 2D histograms to units of relative frequency and then averaged the histograms across subjects. Inspecting the group-average histogram (shown in **Figure 5B**), we manually clicked a sequence of points on the unit sphere that follow the data density as closely as possible. We then parameterized the path traced by these points (a simple 1D manifold) by positioning regularly spaced points where successive points are separated by six angular degrees (**Figure 5B, cyan dots**). The timecourses corresponding to the resulting set of 20 points were cubic interpolated to a sampling rate of 0.1 s and normalized to peak at 1 (**Figure 5C**). Finally, we fit each timecourse using a double-gamma function as implemented in SPM’s *spm_hrf*.*m* (*hrf_fitspmhrftomanifold*.*m*). This yielded a library of 20 canonical HRFs that may be useful for application to other experimental datasets (*getcanonicalhrflibrary*.*m*).

#### Cross-validation framework for single-trial GLM

The GLMdenoise and ridge regression analysis components of GLMsingle both require tuning of hyperparameters. To determine the optimal setting of hyperparameters, we use a cross-validation approach in which out-of-sample predictions are made for single-trial beta estimates, as opposed to time-series data. This simplifies and reduces the computational requirements of the cross-validation procedure. Note that because of cross-validation, although GLMsingle produces estimates of responses to single trials, it does require the existence of and information regarding repeated trials, i.e., trials for which the stimulus is the same.

The first step of the cross-validation procedure is to analyze all of the available data using no regularization. In the case of GLMdenoise, this amounts to the inclusion of zero nuisance regressors; in the case of ridge regression, this amounts to the use of a shrinkage fraction of one, indicating ordinary least-squares regression. In both cases, the analysis produces a full set of unregularized single-trial betas (e.g., in one NSD session, there are 750 single-trial betas distributed across 12 runs). The second step of the procedure is to systematically evaluate different settings of the hyperparameter (e.g., number of nuisance regressors; shrinkage fraction), each time assessing how well the resulting beta estimates generalize to left-out runs. For example, in leave-one-run-out cross-validation, one run is held out as the validation run, stimuli that occur in both the training runs and the validation run are identified, and squared errors between the regularized beta estimates from the training runs and the unregularized beta estimates from the validation run are calculated. This procedure is iterated with each run serving as the validation run, and errors are summed across iterations.

#### The GLMsingle algorithm

Given the procedures described above, we now turn to the steps in the GLMsingle algorithm. GLMsingle involves fitting several different GLM variants. Each variant includes polynomial regressors to characterize the baseline signal level: for each run, we include polynomials of degrees 0 through round(*L*/2) where *L* is the duration in minutes of the run.

1. *Fit a simple ON-OFF GLM*. In this model, stimulus trials are treated as instances of a single experimental condition, and a canonical HRF is used (*getcanonicalhrf*.*m*). Thus, there is a single “ON-OFF” predictor that attempts to capture signals driven by the experiment. The utility of this simple model is to provide variance explained (*R*^2^) values that help indicate which voxels carry experiment-driven signals.
2. *Fit a baseline single-trial GLM*. In this model, each stimulus trial is modeled separately using the canonical HRF. This model provides a useful baseline for comparison.
3. *Identify HRF for each voxel*. We fit the data multiple times with a single-trial GLM, each time using a different HRF from the library of HRFs. For each voxel, we identify which HRF provides the best fit to the data (highest variance explained), and inherit the single-trial betas associated with that HRF.
4. *Use GLMdenoise to determine nuisance regressors to include in the model*. We define a pool of noise voxels (brain voxels that have low ON-OFF *R*^2^) and then perform principal components analysis. The top principal components are added one at a time to the GLM until cross-validation performance is maximized on-average across voxels.
5. *Use fractional ridge regression to regularize single-trial betas*. With the nuisance regressors determined, we use fractional ridge regression (Rokem and Kay, 2020) to estimate the single-trial betas, systematically evaluating different shrinkage fractions. Cross-validation is used to select the optimal shrinkage fraction for each voxel. To mitigate bias on the overall scale of betas, we apply a post-hoc scaling and offset on betas obtained for a given voxel in order to match, in a least-squares sense, the unregularized betas obtained for that voxel.

#### Application of GLMsingle to the NSD data

We used GLMsingle to analyze the time-series data from each NSD scan session (*glm_nsd*.*m*). Major algorithmic parameters included the following: we evaluated up to 10 nuisance regressors; we evaluated shrinkage fractions from 0.05 to 0.90 in increments of 0.05 and from 0.91 to 1 in increments of 0.01 (representing a finer grain for voxels with the best signal-to-noise ratio); we performed 6-fold cross-validation (consecutive pairs of runs) for Steps 4 and 5; and we used an ON-OFF *R*^2^ threshold of 5% in Step 4.

Three different versions of the single-trial betas were computed and saved. The first beta version (b1, ‘betas_assumehrf’) is the result of Step 2, and reflects the use of a canonical HRF. The second beta version (b2, ‘betas_fithrf’) is the result of Step 3, and reflects the result of voxel-wise HRF estimation. The third beta version (b3, ‘betas_fithrf_GLMdenoise_RR’) is the result of Step 5, and reflects the additional GLMdenoise and ridge regression procedures. Betas were converted to units of percent BOLD signal change by dividing amplitudes by the mean signal intensity observed at each voxel and multiplying by 100.

For user convenience, we created preparations of the single-trial betas in additional spaces other than the native 1.8-mm and 1.0-mm functional spaces. For the ‘nativesurface’ preparation, we performed cubic interpolation of the 1.0-mm betas onto each of the 3 cortical surface depths and averaged across depths (*analysis_transformfsaverage*.*m*). The result was then mapped using nearest-neighbor interpolation to fsaverage space to create the ‘fsaverage’ preparation. For the ‘MNI’ preparation, we mapped the 1.0-mm betas to MNI space using cubic interpolation (*analysis_transformMNI*.*m*).

### GLM analysis of the resting-state experiment

As an optional resource, we fit the time-series data from the resting-state experiment using methods that parallel those used for the NSD experiment (*glm_nsdresting*.*m*). For each scan session involving resting-state, we took the two resting-state runs (first and last run acquired) and analyzed the data using the design matrix of the neighboring NSD runs and the same voxel-wise HRFs determined from analyzing the NSD runs in that scan session (this is analogous to beta version b2). Although there is no reason to think that spontaneous resting-state activity conforms to the 4-s trial structure of the NSD experiment, these resting-state betas may be useful as a direct comparison for the NSD betas.

### Noise ceiling estimation

To obtain a measure of data quality, noise ceilings were estimated for the NSD betas (*export_noiseceiling*.*m*). The noise ceiling for a given voxel is defined as the maximum percentage of variance in the voxel’s responses that can in theory be explained, given the presence of measurement noise. Our method for estimating the noise ceiling follows the general framework laid out in previous studies (Kay et al., 2013b; Lage-Castellanos et al., 2019). Several assumptions are made: (i) the signal contained in the voxel’s response is determined solely by the presented image, (ii) the variability of the signal across different images is Gaussian-distributed, (iii) the noise is Gaussian-distributed with zero mean, and (iv) the response to an image is equal to the signal plus noise:

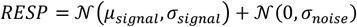

where *RESP* indicates the NSD beta observed on a given trial, μ_signal_ is the mean signal across different images, σ_signal_ is the standard deviation of the signal across different images, and σ_noise_ is the standard deviation of the noise. Note that the first Gaussian distribution characterizes true signal variability, whereas the second Gaussian characterizes variability due to noise. Also, note that this framework treats response variability unrelated to the stimulus as “noise”, but such variability may in fact reflect “signal” from the perspective of functional connectivity (Biswal et al., 1995).

To compute the noise ceiling, we first take the trial-wise NSD betas for each voxel and *z*-score these betas within each scan session. This simple normalization compensates for nonstationarities that may exist across sessions. We then calculate the variance of the betas across the three presentations of each image (using the unbiased estimator that normalizes by *n*–1 where *n* is the sample size), average this variance across images (thus, stabilizing the variance estimates), and then compute the square-root of the result. This produces an estimate of the noise standard deviation:

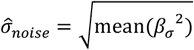

where *α*_γ_^2^indicates the variance across the betas obtained for a given image. Next, given that the variance of the *z*-scored betas is 1, we estimate the signal standard deviation as follows:

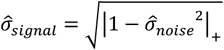

where | |_+_ indicates positive half-wave rectification. Finally, we simplify by calculating a single scalar quantity:

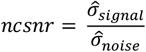

where *ncsnr* indicates the noise ceiling signal-to-noise ratio.

Given the framework described above, the noise ceiling can be calculated as the amount of variance contributed by the signal expressed as a percentage of the total amount of variance in the data:

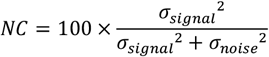

where *NC* indicates the noise ceiling. We would like to be able to calculate the noise ceiling based on the single scalar *ncsnr*. Moreover, since a researcher may wish to average across multiple presentations of each image before attempting to explain the NSD betas, we would like a method for flexibly expressing the noise ceiling for different levels of trial-averaging. With some algebra, it can be shown that the noise ceiling can be expressed as follows:

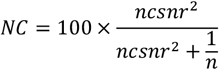

where *n* indicates the number of trials that are averaged together (see online notes for additional details). We note that there is a direct relationship between the metric of split-half reliability and the noise ceiling: if a voxel has two sets of responses that reflect the same image presentations, then the correlation between the two sets of responses multiplied by 100 is equal to the noise ceiling for single-trial responses expressed in percent variance explained.

Using the above methods, we calculated noise ceilings for each of the beta versions and for each of various spatial preparations (1.8-mm, 1-mm, fsaverage, nativesurface). For simplicity, noise ceiling estimates were calculated using betas associated with images with all three presentations available. The noise ceiling results shown in **Figure 5F–G** were computed assuming *n =* 3, reflecting the scenario in which trial-wise betas are averaged across three trials for each image.

Note that our noise ceiling metric refers to activity levels in individual voxels in individual subjects. It is thus quite different from, for example, noise ceiling metrics computed for group-average representational dissimilarity matrices (Nili et al., 2014). The latter are more abstracted away from the data given that they summarize properties observed across a collection of voxels, reflect second-order computations on activity levels and not activity levels themselves, and probe responses at the group level and not at the individual level.

### Calculation of equivalent trials

To provide a common basis for comparing different datasets, we define the number of equivalent trials present in a dataset as *N* × *ncsnr*^2^ where *N* indicates the number of trials conducted and *ncsnr* is the noise ceiling signal-to-noise ratio (as defined earlier). The assumptions here are that (i) every trial has equal value, irrespective of whether it is used to measure brain responses to an image that has already been shown or a new image (e.g., two trials for one image is equivalent to one trial for two distinct images), and (ii) increases in signal-to-noise ratio (SNR) are equivalent to the collection of additional trials (e.g., doubling SNR is equivalent to collecting 4 times as many trials).

We conducted an auxiliary analysis that directly compares NSD against the BOLD5000 dataset (Chang et al., 2019). The goal of this analysis was to calculate a summary *ncsnr* value for each dataset, so that the number of equivalent trials can be calculated. For fair comparison, both NSD and BOLD5000 were analyzed using the exact same GLM methods described in this paper (beta version b3). We then defined a common brain region on which data quality can be compared. This was done by transforming the nsdgeneral ROI to MNI space and then mapping the resulting MNI mask to each subject in the two datasets. Finally, we computed the median *ncsnr* observed across voxels in the mask in each subject.

The median *ncsnr*, averaged across subjects, was 0.260 for NSD (averaged across the first four NSD subjects), and 0.187 for BOLD5000 (averaged across the four subjects in BOLD5000). This indicates that, despite the longer time duration allocated per trial in BOLD5000 (10 s) compared to NSD (4 s), the quality of a single-trial beta in NSD is higher than that in BOLD5000. Specifically, one NSD trial is approximately equivalent to (0.260)^2^/(0.187)^2^ = 1.93 BOLD5000 trials. This increase in quality is likely due, in part, to the screening of subjects and the ultra-high magnetic field strength (7T) used in NSD. Note that the *ncsnr* metric quantifies the SNR per trial and is expected to be unbiased with respect to the number of repeated trials used to calculate it. Thus, although the exact number of trials per image is different in the NSD and BOLD5000 datasets, the *ncsnr* values can still be directly compared.

### pRF estimation

For this analysis (results shown in **Supplementary Figure 10**), we used version 3 of the NSD betas (b3) in the nativesurface preparation. Betas for each surface vertex were *z*-scored within each scan session, concatenated across sessions, averaged across repeated trials for each distinct image, and then re-normalized using a scale and offset such that 0 corresponds to 0% BOLD signal change and the standard deviation of the betas equals 1.

To prepare stimuli for pRF estimation, the NSD images were converted to grayscale, resized to 800 pixels × 800 pixels (cubic interpolation), and squared to mimic the luminance response of the display. The images were then placed against the gray background and divided into a 51 × 51 grid such that the first and last grid elements were centered at the edges of the stimulus (each grid element spanned 0.168° × 0.168°). Finally, to quantify local contrast, we computed the standard deviation of pixel values within each grid element.

Based on the local-contrast preparation of the NSD images, we used analyzePRF (http://cvnlab.net/analyzePRF/) to fit the Compressive Spatial Summation pRF model (Kay et al., 2013b) to the trial-averaged betas obtained for each vertex. The non-shared NSD images were used as training data; the shared NSD images were used as validation data. pRFs were constrained to have non-negative gain. No offset term was included in the model (opt.maxpolydeg = NaN); thus, the model necessarily predicts a response of 0 for an image with zero contrast. For model fitting, an initial gridding of model parameters was performed (opt.seedmode = 2), and parameter optimization started from the best parameter combination (opt.modelmode = 2; opt.algorithm = ‘trust-region-reflective’). Model fitting produced, for each vertex, an estimate of pRF angle, eccentricity, size, exponent, and gain, as well as variance explained in the training data and the validation data.

### Univariate analysis of memory recognition

For this analysis (results shown in **Figure 6B**), we used version 3 of the NSD betas (b3) in the fsaverage preparation. Betas for each surface vertex were kept in percent signal change units. Using the behavioral responses, we identified trials involving hits (subjects responded ‘old’ to a previously presented image) and trials involving correct rejections (subjects responded ‘new’ to a novel image). Then, for each subject, two-sample *t*-tests were performed at each surface vertex. This was done both for trials pooled within individual NSD scan sessions as well as for trials pooled across all sessions.

### Representational similarity analysis

For this analysis (results shown in **Figure 7**), we used version 3 of the NSD betas (b3) in the fsaverage preparation. Betas for each surface vertex were *z*-scored within each scan session, concatenated across sessions, and averaged across repeated trials for each distinct image. To support the representational similarity analysis (Kriegeskorte et al., 2008), we defined a set of ROIs (V1, V2, V3, pVTC, aVTC) on the fsaverage surface. This was done by mapping the manually-defined V1, V2, and V3 from each subject to fsaverage, averaging across subjects, and using the result to guide the definition of group-level ROIs. We also defined a posterior and anterior division of ventral temporal cortex (pVTC and aVTC, respectively) based on anatomical criteria.

For each subject, we extracted betas for vertices within each ROI (concatenating across hemispheres). We then computed Pearson’s correlation between beta patterns across all possible pairs of images. This yielded representational dissimilarity matrices (RDMs) with rows and columns indexing distinct images (e.g., the RDMs for subject 1 have dimensionality 10,000 × 10,000 with correlations corresponding to 49,995,000 possible pairs). To help visualize and interpret these large dissimilarity matrices, we performed *t*-distributed stochastic neighbor embedding (Maaten and Hinton, 2008; Pedregosa et al., 2011) (t-SNE) using a perplexity level of 100. This projects the high-dimensional representations onto a two-dimensional plane such that the distance of a given pair on the plane reflects that pair’s distance in the high-dimensional representation as accurately as possible.

### Encoding models based on deep convolutional neural networks

For this analysis (results shown in **Figure 8**), we used version 3 of the NSD betas (b3) in the 1.8-mm volume-based preparation. Before modeling, betas for each voxel were *z*-scored within each scan session and concatenated across sessions. Models were implemented using PyTorch (code available online).

#### Model architecture

We considered several variants of voxel-wise encoding models (Naselaris et al., 2011) that attempt to predict the NSD betas. All three models consist of (i) a feature extractor implemented as a convolutional neural network (CNN) and (ii) a network-to-brain coupling model that maps extracted features into predictions of activity observed for individual voxels. cIn the first model (AlexNet), the feature extractor is the AlexNet CNN (Krizhevsky et al., 2017), a *task-optimized* network that has been trained to classify object categories in the ImageNet database (Deng et al., 2009). In the second model (GNet), the feature extractor is a different CNN—referred to here as ‘GNet’—a *brain-optimized* network that is trained to directly predict brain activity in the NSD dataset. The third model is a simple control model in which the feature extractor consists of a single fixed layer of Gabor filters (Kay et al., 2008). The specific network architectures for AlexNet and GNet are illustrated in **Supplementary Figure 12**.

To facilitate direct comparison, all models are designed to have comparable coupling models. For GNet, both the feature extractor and coupling model are trained jointly using brain data; for AlexNet and the Gabor models, the feature extractors are fixed and only the coupling model is trained using brain data.

The CNNs in the AlexNet, GNet, and Gabor models consist of hierarchically composed functions of an input image *x*:

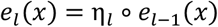

where η_*l*_ is a feature extractor that operates at layer *l* on the output of *e*_*l*−1_(x) (also a composite function). e_*l*_ may denote an arbitrary sequence of transformations. The encoding models leverage the intermediate representations *e*_*l*_(x), which are feature maps with pixels denoted by [*e*_*l*_(x)]_*kji*_, where (*i, j*) is the location of the pixel in the *k*th feature map. Predicted brain activity for voxel *v*, 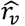, is given by the expression:

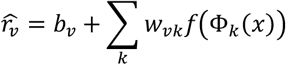

where *w*_*vk*_ are feature weights for voxel *v* and feature *k, b*_*v*_ is a bias term,

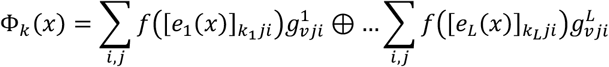

*f*(·) is typically a compressive nonlinearity, 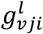 indicates a weight assigned to pixel (*i, j*) in the *l*th feature map, and ⊕ denotes summation along the feature axis *k* = (*k*_1_, … *k*_*L*_). Note that this formulation incorporates feature-space separability, which reduces overfitting and generally improves prediction accuracy for brain activity (St-Yves and Naselaris, 2017).

In the Gabor model, the feature extractor consists of a single fixed set of convolutions involving 12 log-spaced spatial frequency Gabor wavelets between 3 and 72 cycles/stimulus and constructed at 6 evenly spaced orientations between 0 and *π* (St-Yves and Naselaris, 2017).

#### Spatial pooling fields

Constraints were placed on the weights 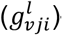—termed ‘spatial pooling fields’—that couple the feature maps to voxel activity. For the AlexNet- and Gabor-based encoding models, the spatial pooling field for each voxel was a 2D isotropic Gaussian that was applied to all feature maps (see **Supplementary Figure 12B, middle**). We find that this constrained model of spatial pooling typically yields better prediction accuracy (relative to other possible variants) in the scenario where feature extraction parameters are fixed (St-Yves and Naselaris, 2017). For the GNet-based encoding model, the weights of the spatial pooling fields were independently adjustable; hence, we refer to these as flexible spatial pooling fields (see **Supplementary Figure 12B, left**). Feature maps with the same spatial resolution were grouped together, and a distinct, independently optimized spatial pooling field was applied to each group. Thus, the GNet model for each voxel was specified by multiple, independently optimized spatial pooling fields.

#### Model training and validation

Given the demanding memory requirements of training large-scale neural networks to jointly predict tens of thousands of voxels, we selected the four NSD subjects with the highest noise ceilings (see **Figure 5G**). For the selected subjects (1, 2, 5, 6), NSD betas were extracted from visual areas V1–hV4. These betas were separated into those evoked by the shared1000 images and those that were not; the former were designated as the validation set, while the latter were designated as the training set. For example, for subject 1, there were 9,000 images × 3 trials = 27,000 samples in the training set, and 1,000 images × 3 trials = 3,000 samples in the validation set. After model training, accuracy was quantified as the voxel-wise correlation between model predictions and observed responses in the validation set.

For the AlexNet-based encoding model, parameters of the feature extractors were pre-trained based on classification of objects in the ImageNet database (Krizhevsky et al., 2012). For both the AlexNet- and Gabor-based encoding models, feature weights for the coupling model were optimized via ridge regression, with the ridge parameter selected to maximize accuracy on a held-out subset (20%) of the training data. Line search was used to optimize the position and size of the Gaussian spatial pooling field for each voxel (see **Supplementary Figure 12B, right**).

For the GNet-based encoding model, parameters of the feature extractors, spatial pooling fields, and feature weights were all optimized via stochastic gradient descent of an *L*_2_-norm weighted loss function:

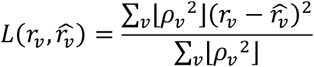

where ⌊*ρ*_*v*_^2^⌋ is the batchwise prediction accuracy for a given voxel *v* with an imposed floor of 0.1 in order to permit contribution of yet-to-be-predicted voxels.

Two versions of the GNet model were developed and evaluated. In the single-subject GNet model, different instantiations of GNet were created for different subjects, and only the data from a given subject were used to train the GNet-based encoding model for that subject. In the multiple-subject GNet model, a single instantiation of GNet was created for all four subjects, and data from all subjects were used to train the GNet-based encoding models. In this scheme, all subjects share a common feature extractor, but each subject has independently adjusted coupling models and feature weights.

To train the GNet-based encoding model, stochastic gradient descent with early stopping was performed using the ADAM optimizer (Kingma and Ba, 2017) (*l*r = 10^*−*3^, β_1_ = 0.99, β_2_ = 0.99). Parameter updates for feature extractors, spatial pooling fields, and feature weights were alternated to promote stability of the training procedure.

Note that the NSD subjects view largely non-overlapping sets of images. Thus, when training GNet on data from multiple subjects, we used a modified procedure for selecting batches of training data. For each iteration of training, we first extracted a batch of training samples from one subject’s data and calculated the gradient with respect to the loss function. Coupling model parameters for that subject and feature extractor parameters were then updated and the process was repeated until all batches from all subjects were used. This corresponded to one training epoch.

## Supplementary Discussion

Our examination of the NSD data did not reveal any severe problems or artifacts that could not be compensated for in data pre-processing. Nonetheless, there are several known limitations of the data that should be considered. EPI pulse sequences invariably suffer from signal dropout and spatial distortion in locations with magnetic field inhomogeneties. The NSD data exhibit signal dropout in typical locations such as near the ear canals and the frontal sinuses (see **Figure 5F**), and our approach for distortion correction may have some imperfections (see **Supplementary Video 6**). While we stressed the importance of central fixation to the NSD subjects, some inadvertent eye movements are likely present in the data. Methods for retrospectively estimating eye position from imaging data (Frey et al., 2020; Son et al., 2020) might prove useful. In a few fMRI scan sessions (6 out of 308; 1.9%), the subject exited the scanner and re-entered either on the same day or a different day to complete the session. We compensated for these occurrences in the pre-processing of the data, but they nonetheless contribute some variability. Due to hardware errors, behavioral responses are missing for a few of the NSD runs (2 out of 3,408; 0.06%). Finally, a few fMRI scan sessions (4 out of 308; 1.3%) had slightly incomplete brain coverage due to subject motion. A comprehensive summary of data anomalies is available in the online materials.

## Supplementary Figures

**Supplementary Figure 1.**
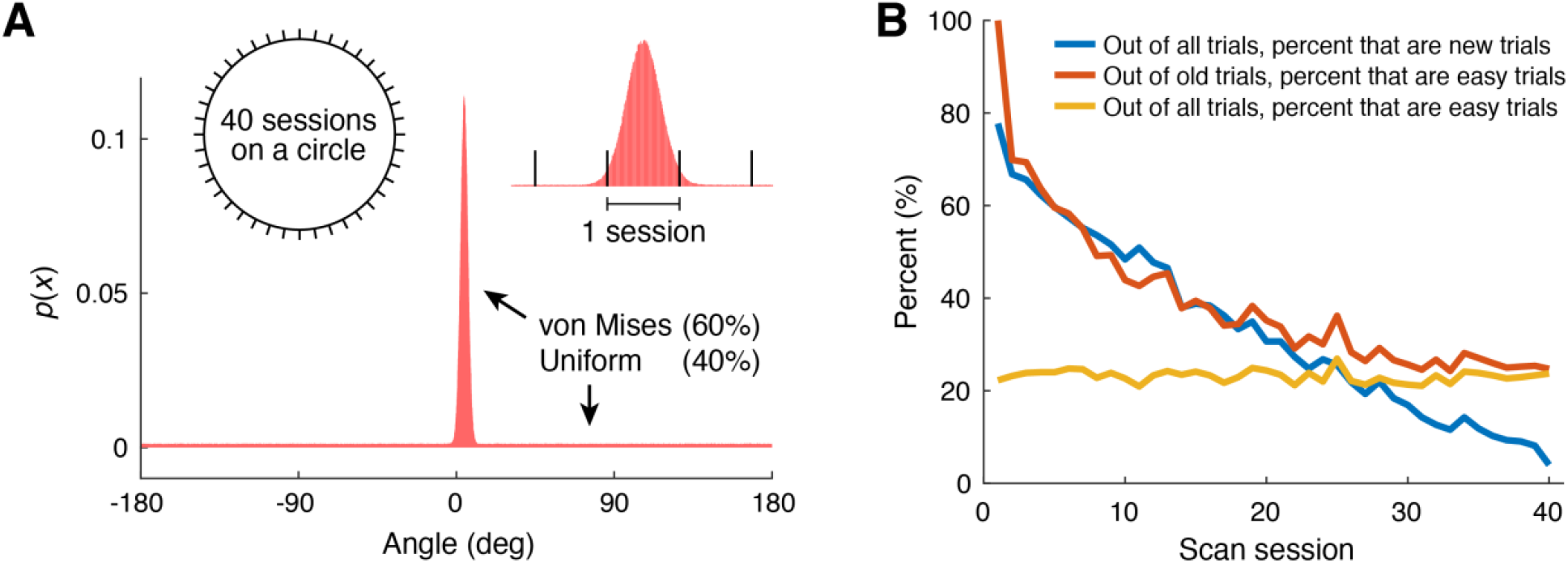
Design of the NSD experiment. *A*, Image presentations. Each of 10,000 distinct images was placed 3 times on a circle according to a probability distribution created by mixing a relatively narrow von Mises distribution and a uniform distribution. The resulting image sequence was divided into 40 equally-sized segments for the 40 NSD scan sessions. *B*, Basic statistics of image repetitions. We define *new trial* as a trial involving an image never shown before, *old trial* as a trial that is not a new trial, and *easy trial* as an old trial for which the presented image had been shown previously in the same scan session.

**Supplementary Figure 2.**
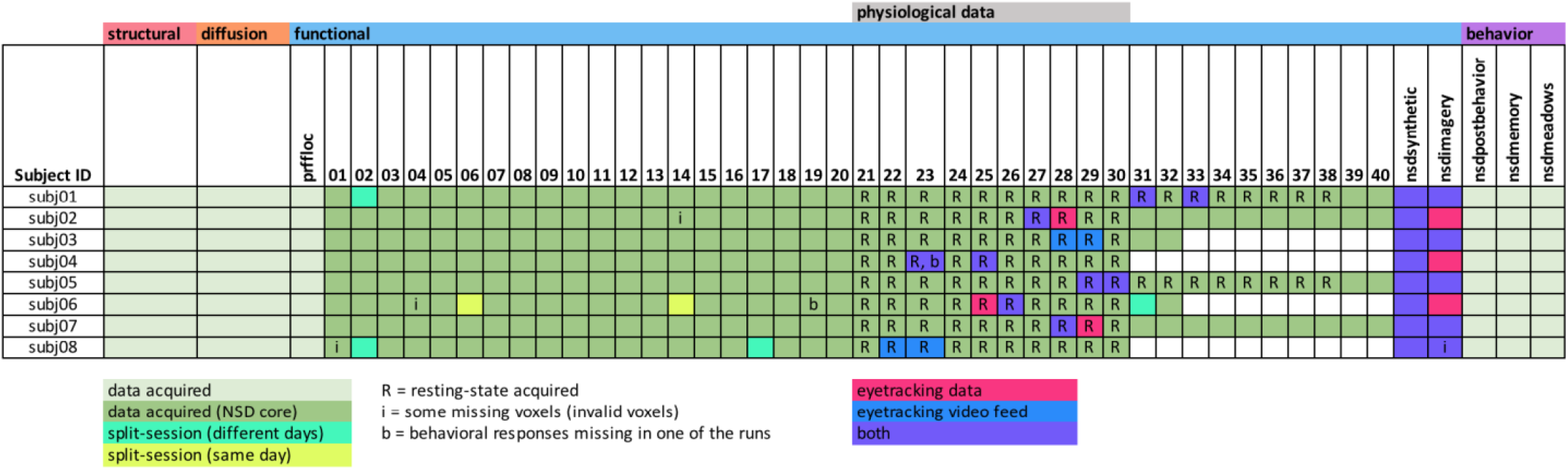
Overview of data collection. This table summarizes the overall NSD data collection effort. Structural and diffusion MRI data were collected at 3T. Functional MRI data were collected at 7T. The breakdown of the 7T fMRI scan sessions is indicated: for example, subject 2 participated in 1 (prffloc) + 40 (nsd01–nsd40) + 1 (nsdsynthetic) + 1 (nsdimagery) = 43 7T fMRI scan sessions. Additional behavioral data were acquired outside of the scanner (nsdpostbehavior, nsdmemory, nsdmeadows). Note that scan sessions were occasionally split across multiple magnet entries (see aquamarine and yellow cells). For simplicity, we treat these cases as if they represent single scan sessions.

**Supplementary Figure 3.**
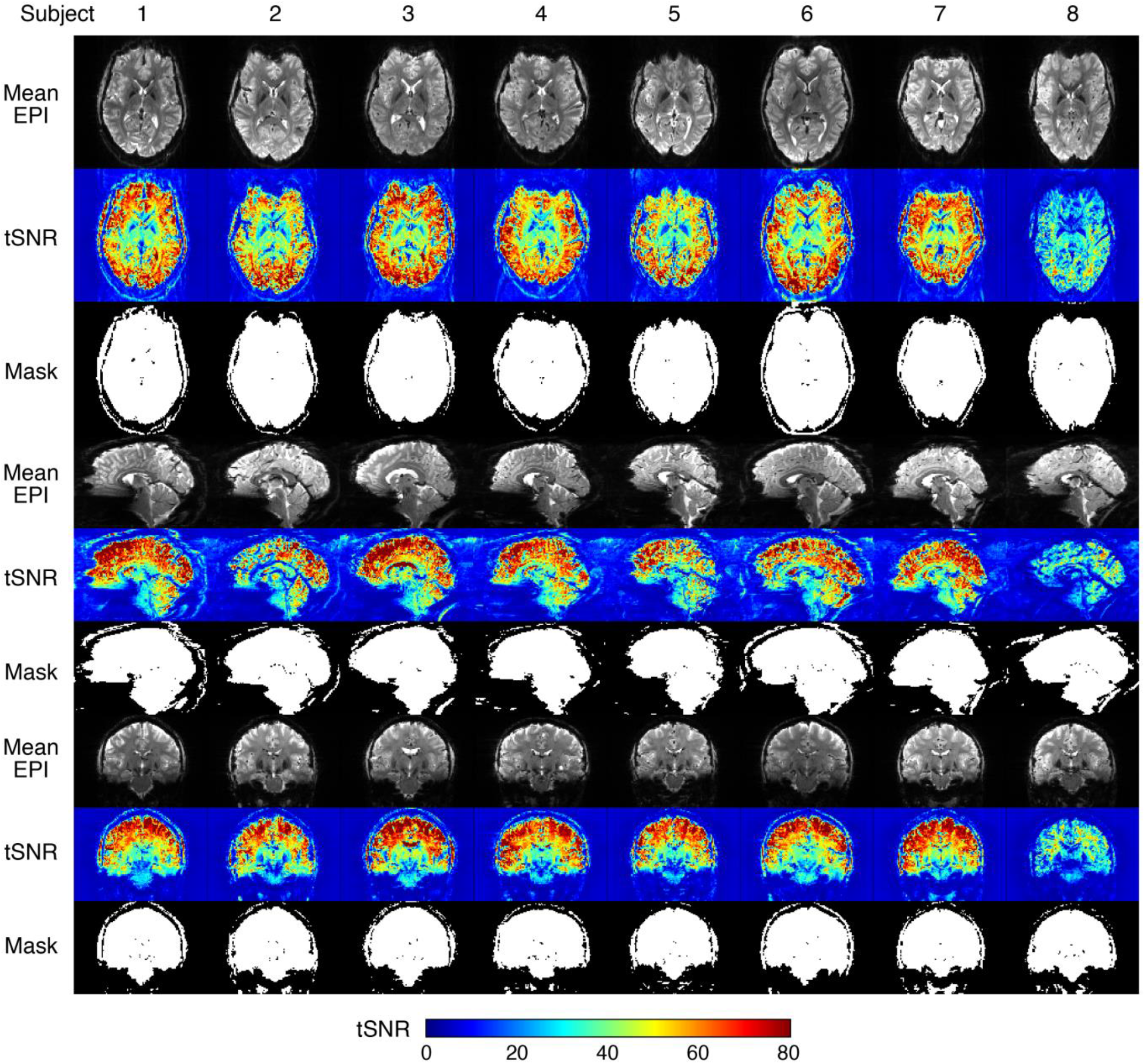
Details on the quantification of tSNR. This figure shows example tSNR results (nsd20 scan session, first NSD run). The middle slice in each of three orthogonal views (axial, sagittal, coronal) is displayed. To compute tSNR, the raw fMRI volumes from a given run are obtained, and the mean across volumes is computed (Mean EPI). A brain mask is computed by identifying voxels whose intensity is at least 10% of the 99th percentile of intensities in the mean volume (Mask). tSNR is calculated by quadratically detrending the time-series of each voxel (preserving the mean) and then computing the mean divided by the standard deviation of the time-series values (tSNR). A summary tSNR value is determined by calculating the median tSNR across voxels within the brain mask. This corresponds to the summary metric shown in **Figure 2D, left** (the inset shows results from subject 2).

**Supplementary Figure 4.**
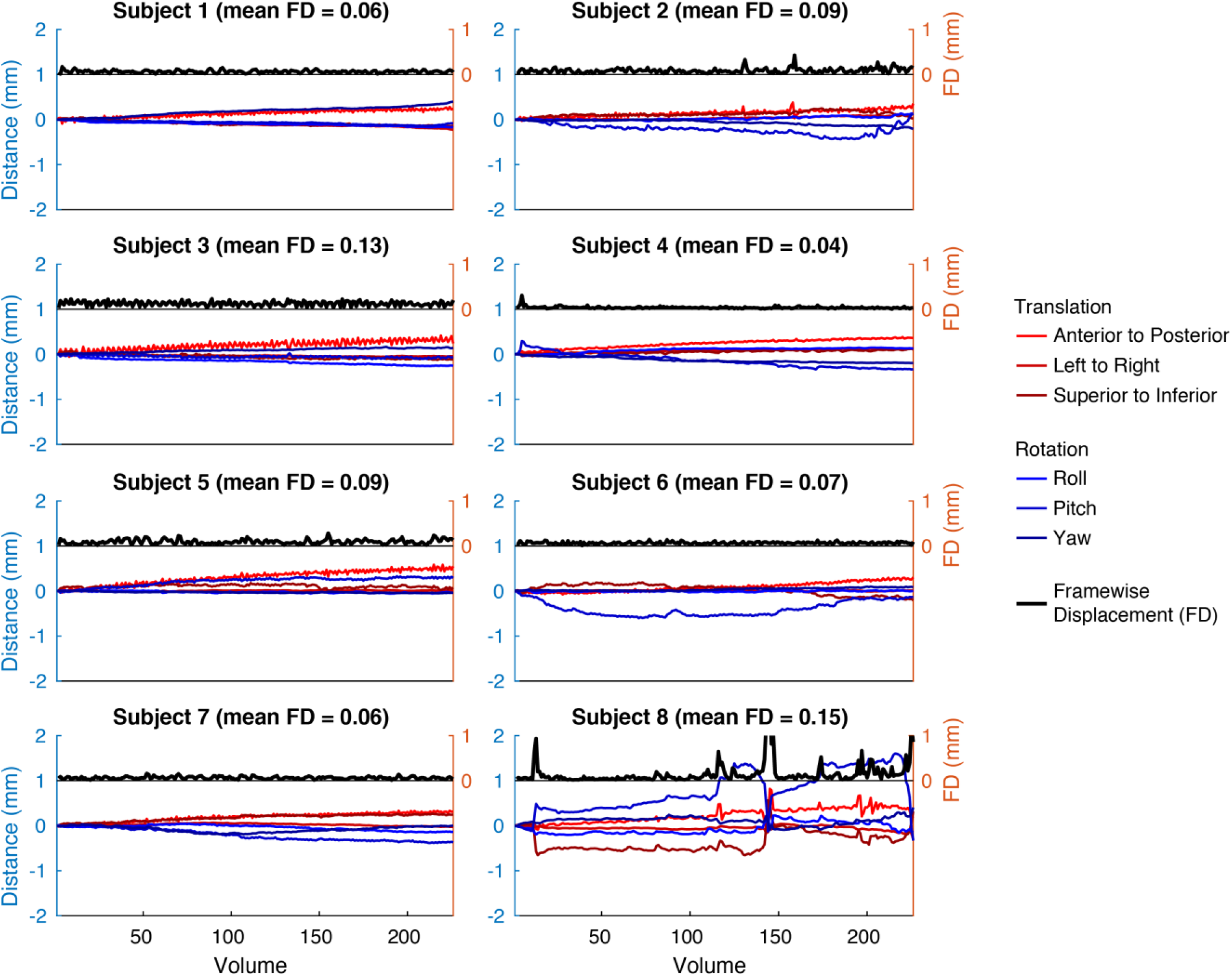
Details on the quantification of head motion. This figure shows example motion parameter estimates obtained during data pre-processing (nsd20 scan session, first NSD run, 1.8-mm data preparation). The rotation parameters, originally in radian units, are multiplied by 50 in order to allow interpretation in terms of millimeters of displacement for a circle of diameter 100 mm (Power et al., 2012). Motion parameters are relative to the reference volume which is the first volume in each scan session. Framewise displacement (FD), calculated as the sum of the absolute differences of the motion parameters for successive pairs of volumes, is also plotted. The mean FD across volumes is indicated in the plot titles, and corresponds to the summary metric shown in **Figure 2D, middle** (the inset shows results from subject 5).

**Supplementary Figure 5.**
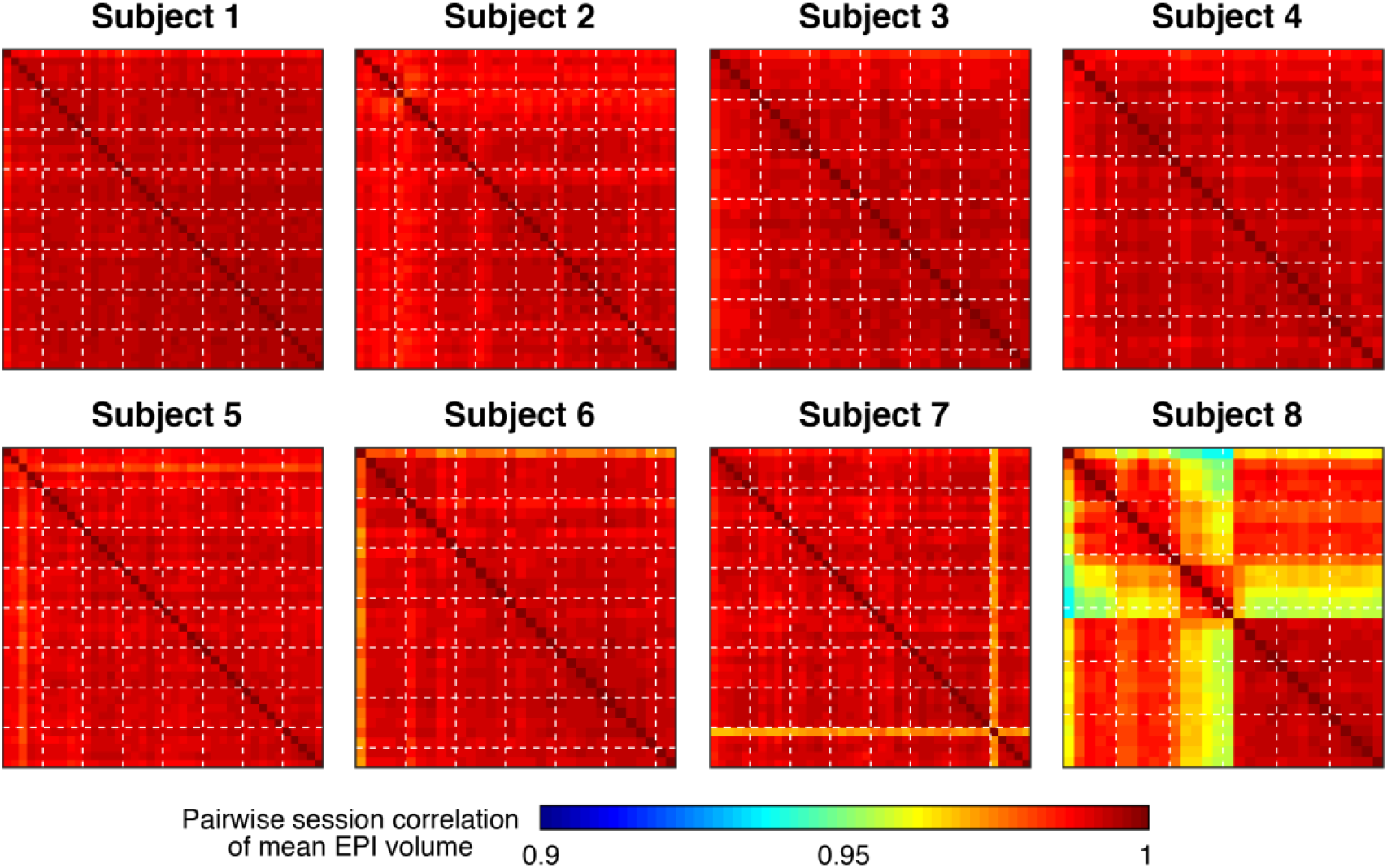
Quantification of functional imaging stability. We took the mean fMRI volume (1-mm preparation) in each scan session, bias-corrected the volume by dividing by a fitted 3D polynomial (*autoqc_nsd_grand*.*m*), and then computed pairwise correlation across sessions. Dotted white lines mark increments of five NSD scan sessions. Inspection of similarity of the mean EPI volume across sessions reveals a few minor anomalies. We investigated these cases further and determined the following: the nsd36 scan session in subject 7 involved a poor scanner shim, which was largely but not fully corrected by the fieldmap-based processing; the nsd01 scan session in subject 8 involved an unusually large amount of head motion, which resulted in some residual spatial distortion; and the nsd12–nsd16 scan sessions in subject 8 involved a temporary sinus infection near frontal cortex that manifested as bright signal intensities in the EPI volumes outside the brain but otherwise did not cause any data problems. For visual inspection of these effects, see **Supplementary Videos 6–7**.

**Supplementary Figure 6.**
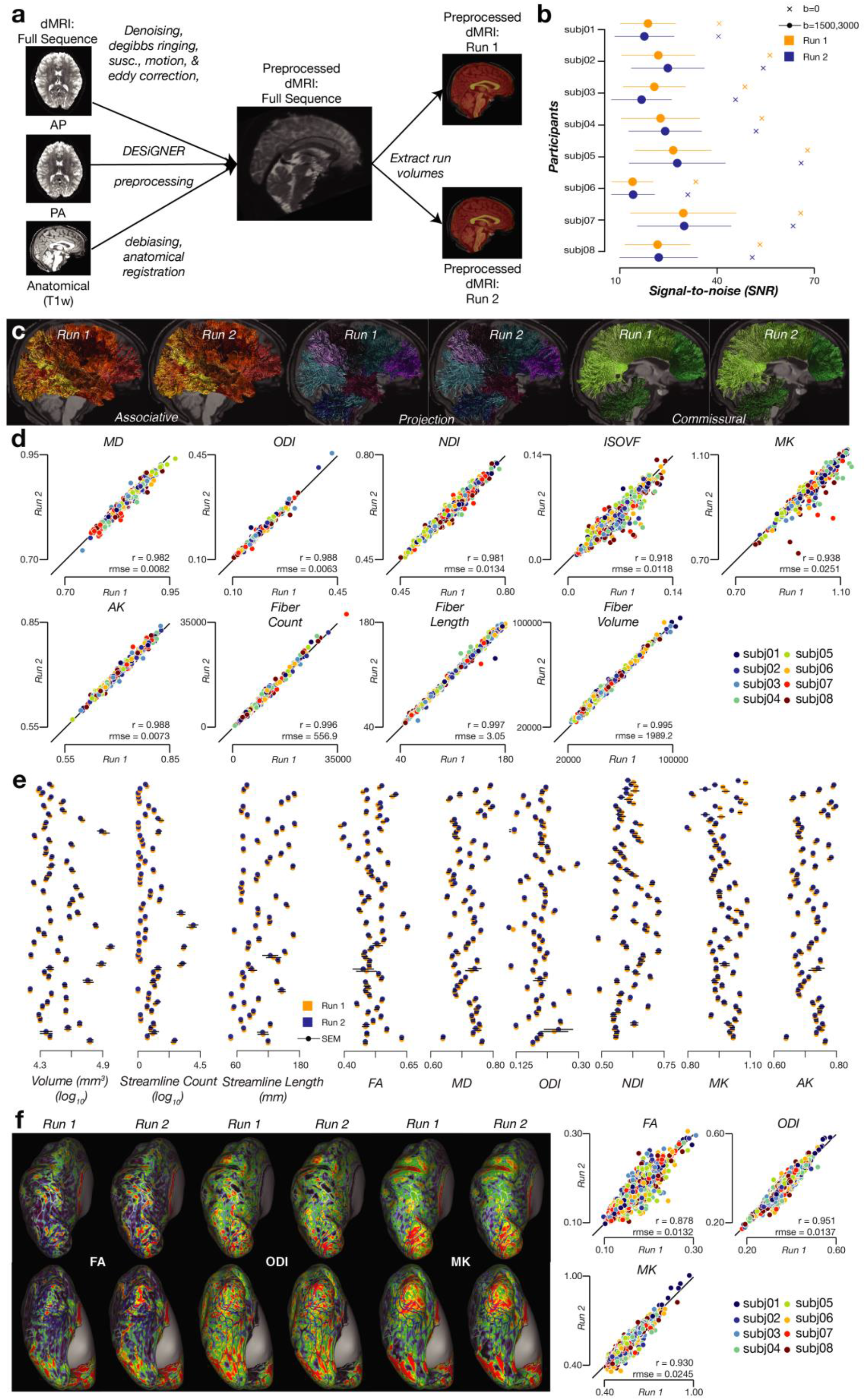
Diffusion processing for investigation of white-matter connectivity. *A*, Schematic of the diffusion pre-processing pipeline. Diffusion volumes were corrected for noise, Gibbs ringing, susceptibility, motion, eddy currents, and bias fields before being co-registered to the *T*_1_ anatomy. Following pre-processing, the data were organized into two runs (corresponding to the 99-direction and 100-direction scans, respectively). *B*, Signal-to-noise ratio, computed in the corpus callosum. *C*, White-matter tract segmentation from an example subject (subject 7). White-matter tracts are organized based on typical anatomical and functional definitions into associative (left), projection (middle), and callosal (right) tracts and overlaid on the *T*_1_ anatomy. *D*, Reliability of MD, ODI, NDI, ISOVF, Mean Kurtosis (MK), Axial Kurtosis (AK), and fiber count, length, and volume. Each dot indicates results averaged along a single tract. Pearson’s correlation (*r*) and root-mean-squared error (RMSE) for each measure are indicated in the inset. *E*, Macrostructural and microstructural properties observed for different tracts. Error bars indicate ± 1 SEM across subjects. *F*, Microstructural properties of cortical regions. Shown are tensor (FA; left), NODDI (ODI; middle), and kurtosis (MK; right) results mapped to the cortical surface of the example subject, with dorsal (top) and ventral (bottom) viewpoints of occipital cortex. Quantitative results are shown on the right, where each dot indicates results obtained for a single region in the HCP-MMP1 parcellation.

**Supplementary Figure 7.**
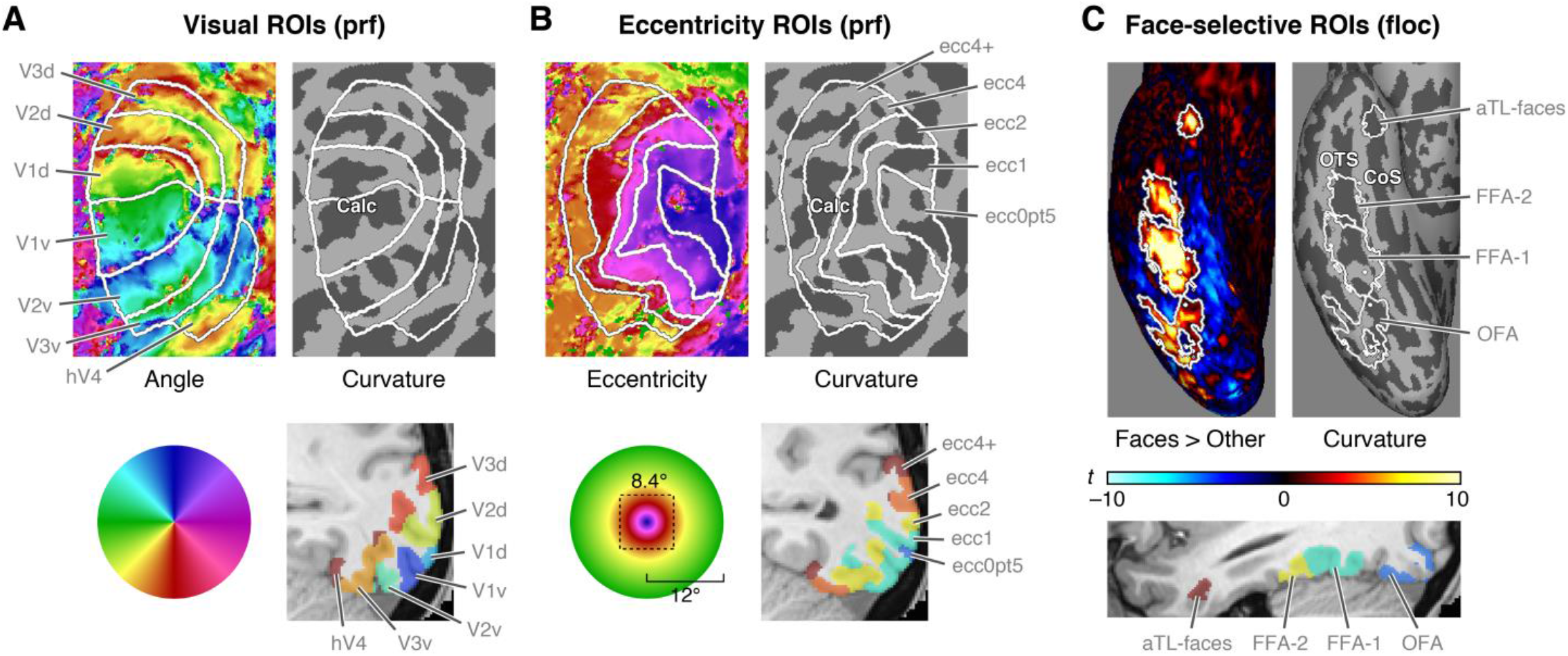
Regions of interest (ROIs) provided with NSD. A variety of ROIs were defined based on auxiliary fMRI experiments (pRF, fLoc). Here we show example results for subject 3, right hemisphere. *A*, Early visual areas. Results are shown on FreeSurfer’s sphere surface as well as in the 0.8-mm anatomical volume space. *B*, Eccentricity-based regions. Similar format to panel A. Note that the total stimulus extent is 8.4° × 8.4° in the pRF, fLoc, and NSD experiments. *C*, Face-selective regions. Regions were defined based on *t*-values computed for the contrast of faces against all other categories. Results are shown on FreeSurfer’s inflated surface as well as in the 0.8-mm anatomical space. Calc = calcarine sulcus, OTS = occipitotemporal sulcus, CoS = collateral sulcus.

**Supplementary Figure 8.**
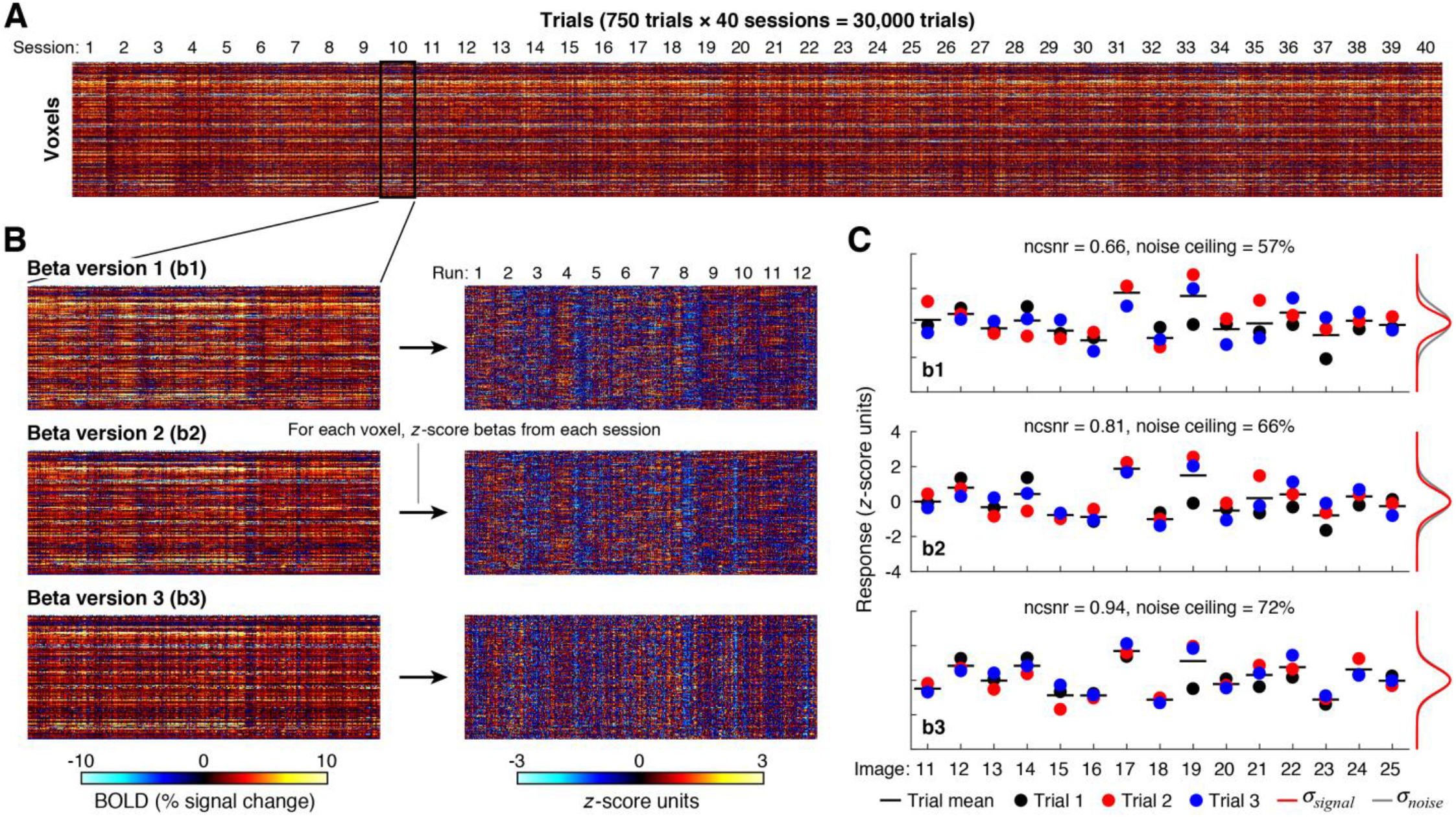
Detailed visualization of NSD betas. We prepared three beta versions (b1, b2, b3) reflecting GLM analyses of increasing sophistication. *A*, Inspection of NSD betas. The full set of estimated single-trial responses (betas) is shown for voxels in subject 1, right hemisphere, FFA-1 (1.8-mm preparation, b1). *B*, Zoomed view of one scan session. Shown are all three beta versions, as well as the result of *z*-scoring betas within each scan session. *C*, Detailed inspection of one voxel. To assess the reliability of evoked responses, we group trials according to the image presented. The estimated signal standard deviation (σ_signal_) and noise standard deviation (σ_noise_) are illustrated at the right of each subplot. In these visualizations, we observe horizontal stripes (panel A), indicative of gross variation in percent BOLD signal change across voxels. We find that the different beta versions generally resemble one another (panel B, left column), implying that the variations in GLM methods do not drastically change the data. Vertical stripes visible in the visualizations (panel B, left column) tend to decrease from b1 to b2, suggesting that fitting voxel-wise HRFs reduces artifacts. Vertical stripes also tend to decrease from b2 to b3, which might reflect the reduction of correlated noise achieved by GLMdenoise. In general, we suggest that users may wish to *z*-score each voxel’s responses within each scan session in order to eliminate potential non-stationarities and to equalize units across voxels. The effect of such normalization is depicted (panel B, right column). Finally, quantitative inspection of the beta versions for a single voxel indicates that b2 and b3 reduce variability of betas across the 3 trials associated with each image (panel C).

**Supplementary Figure 9.**
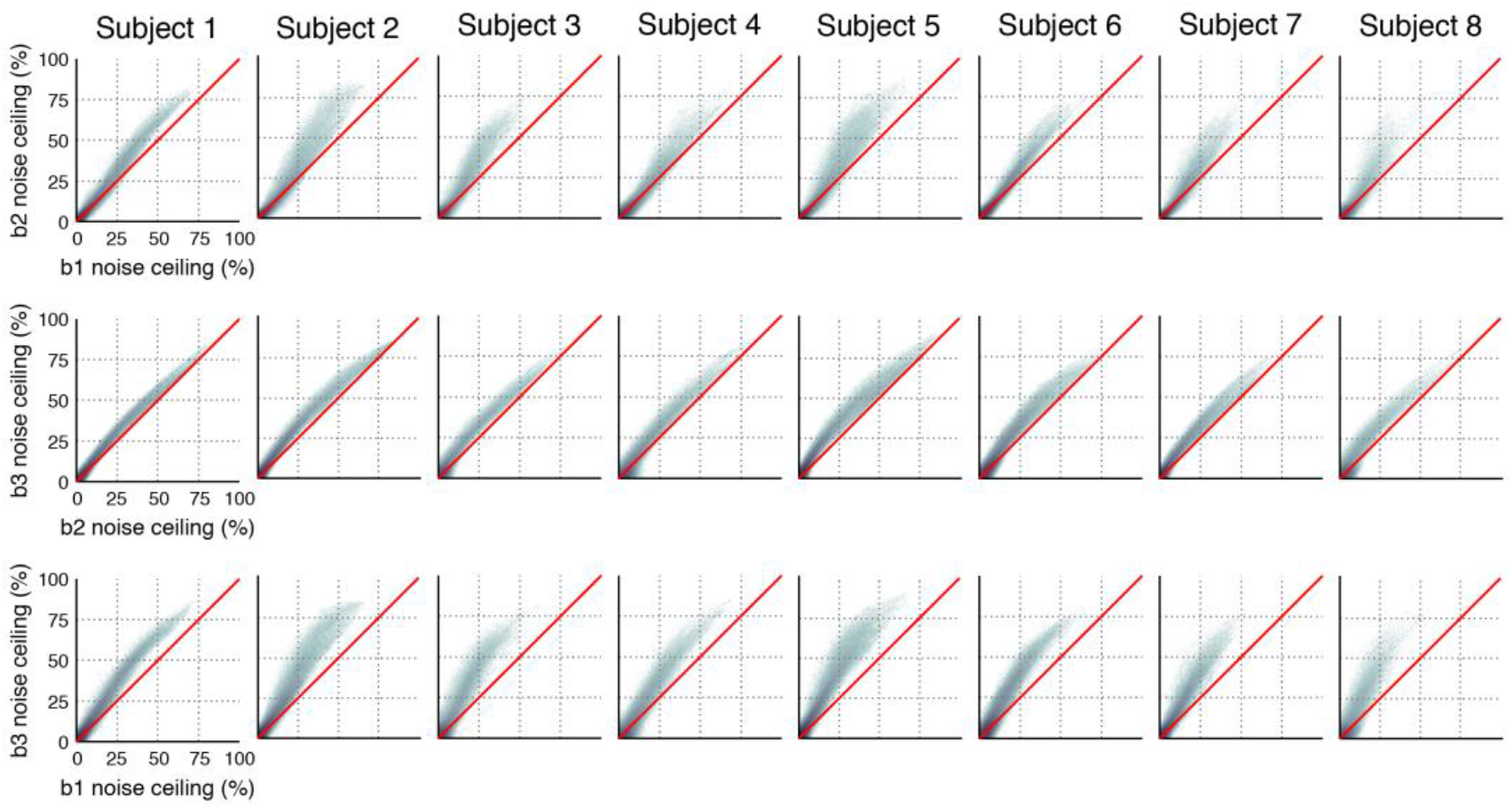
Detailed comparison of noise ceiling results for different beta versions. This figure provides additional detail on results shown in **Figure 5F–G**. Each subplot is a 2D histogram comparing noise ceilings for two different beta versions. Improvements in noise ceilings are consistent across voxels and subjects.

**Supplementary Figure 10.**
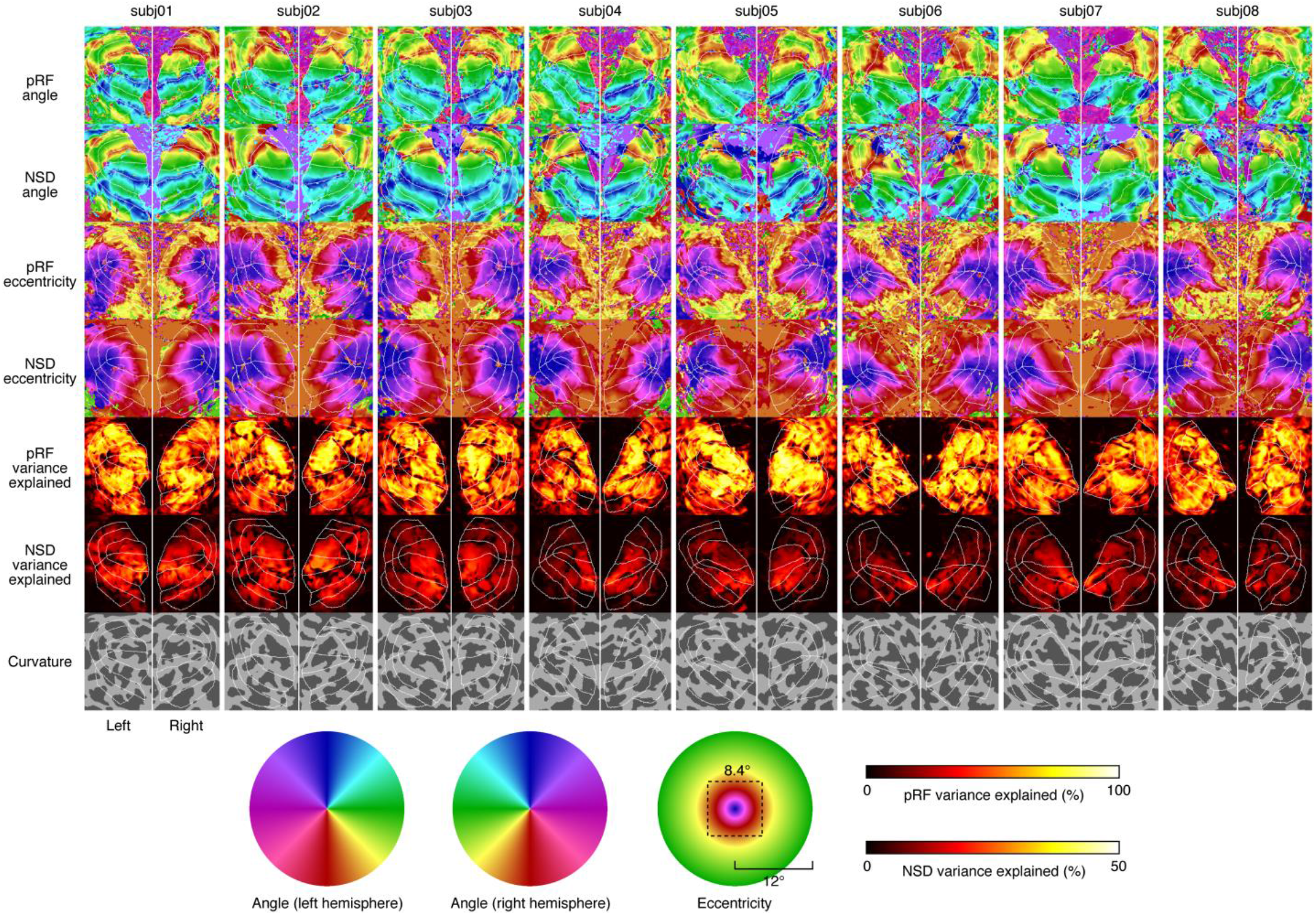
Angle and eccentricity estimates from the NSD data. Here we show results from the analysis of the pRF experiment and results from an analogous analysis performed on trial-averaged NSD betas (see Methods for details). Each panel shows an occipital view of FreeSurfer’s sphere surface, and white lines indicate borders of visual areas V1–hV4 (defined based on results of the pRF experiment). Angle and eccentricity estimates are plotted using the same colormaps as in Benson et al. (Benson et al., 2018). We also plot the amount of time-series variance explained in the pRF data (variance relative to the mean signal level) and the amount of variance explained in the NSD betas (variance relative to 0% BOLD signal change). Clear retinotopic maps in early visual cortex are visible in the NSD results, including robust angle estimates even in foveal regions. In addition, there is high consistency of retinotopic estimates across the pRF and NSD datasets. There is some discrepancy in absolute eccentricity estimates at peripheral locations; this is likely due to technical differences in how modeling procedures behave for voxels near the stimulus edge.

**Supplementary Figure 11.**
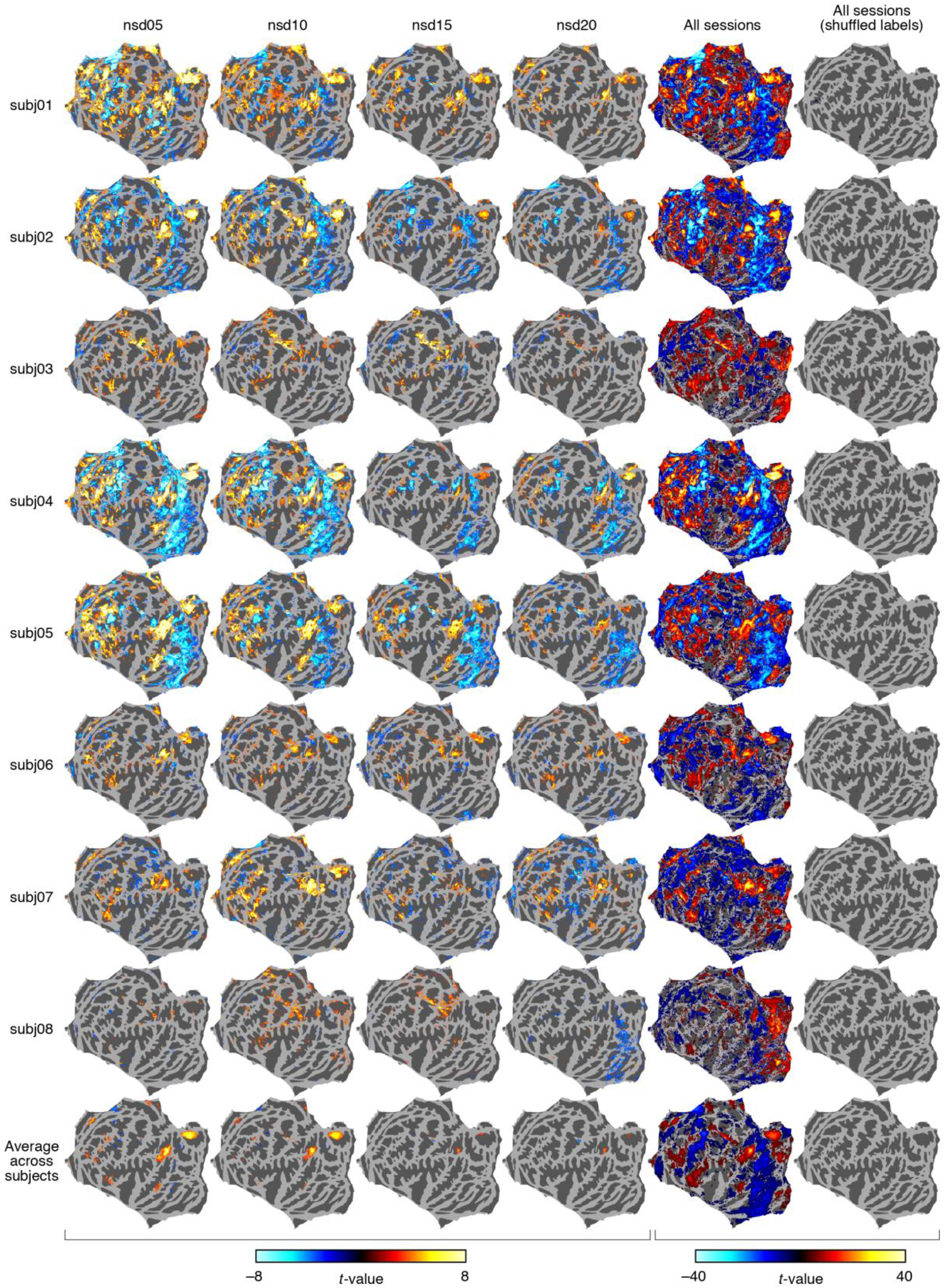
Recognition memory effects in the NSD data. Same format as **Figure 6B**, but showing results for all individual subjects. Note that positive values indicate BOLD responses are greater for hits than for correct rejections, whereas negative values indicate BOLD responses are greater for correct rejections than for hits. The observed decrease in the magnitudes of the t-values (e.g. from nsd05 to nsd20) likely reflects a decrease in the subjects’ recognition accuracy over the course of the NSD experiment.

**Supplementary Figure 12.**
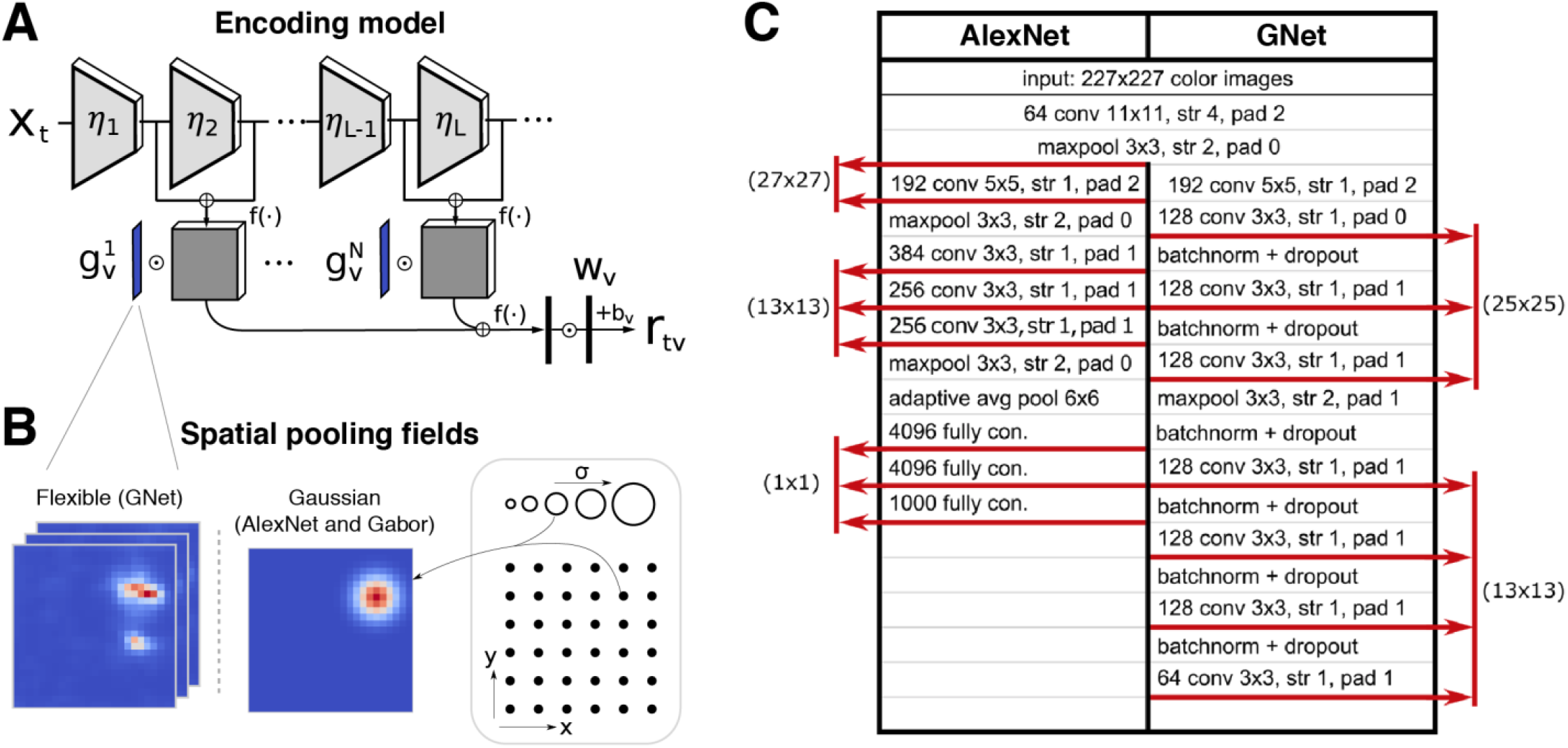
Design of AlexNet- and GNet-based encoding models. *A*, Illustration of an encoding model that predicts brain activity in a given voxel (r_*tv*_) in response to images (x_*t*_). Images are passed to nonlinear feature extractors, *η*_*l*_ (trapezoids), that output feature maps (grey cuboids). Feature maps are grouped, passed through an element-wise nonlinearity, *f*(·), and then multiplied pixel-wise by a spatial pooling field (*g*^1^, …, *g*^*N*^ where superscripts index distinct groups of feature maps) that determines the region of visual space that drives voxel activity. The weighted pixel values in each feature map are then summed, reducing each feature map to a scalar value. These scalar values are concatenated across all feature maps, forming a single feature vector that is passed through another element-wise nonlinearity (left black rectangle) and then weighted by a set of feature weights, *w* (right black rectangle), to yield predicted voxel activity. Note that for each type of encoding model (e.g., AlexNet-based encoding model, GNet-based encoding model), the feature extractors are identical for all voxels, but the spatial pooling fields and feature weights are optimized and may vary across voxels. For the AlexNet-based encoding model, the feature extractors were pre-specified, the spatial pooling fields were optimized via line search, and the feature weights *w* were optimized via ridge regression. For the GNet-based encoding model, stochastic gradient descent with early stopping was used to optimize the parameters of the feature extractors *η*_*l*_, the spatial pooling fields *g*^1^, …, *g*^*N*^, and the feature weights *w. B*, Illustration of spatial pooling fields. For the AlexNet model, a single isotropic 2D Gaussian pooling field (middle) selected from a set of candidates (right) was applied to all feature maps. For the GNet model, an independent, flexible pooling field (left) was applied to each group of feature maps. Applying flexible pooling fields to AlexNet leads to lower prediction accuracy overall (results not shown), so we present the version that uses isotropic 2D Gaussian fields. *C*, Comparative architecture of AlexNet and GNet. AlexNet and GNet are both deep convolutional neural networks, but differ in the types and sequencing of layers (rows of the table). The first three layers are the same for both networks and correspond to the first three layers of an AlexNet trained to classify objects in the ImageNet dataset. For both networks, these shared “pre-filtering” layers are followed by sequences of convolutional layers (rows labeled “conv”; values indicate feature depth and convolutional filter resolution; “str” = filter stride, “pad” = convolutional padding), max-pooling layers (“maxpool”), batch-normalization and weight-dropout layers (“batchnorm + dropout”), adaptive averaging layers (“adaptive avg”), and fully-connected layers (“fully con.”; value indicates number of units). Feature maps in the convolutional or fully connected layers (indicated by red arrows; resolution of the feature maps in parentheses) are used as predictors of brain activity in the context of an encoding model (see panel A).

## Supplementary Videos

**Supplementary Video 1.**
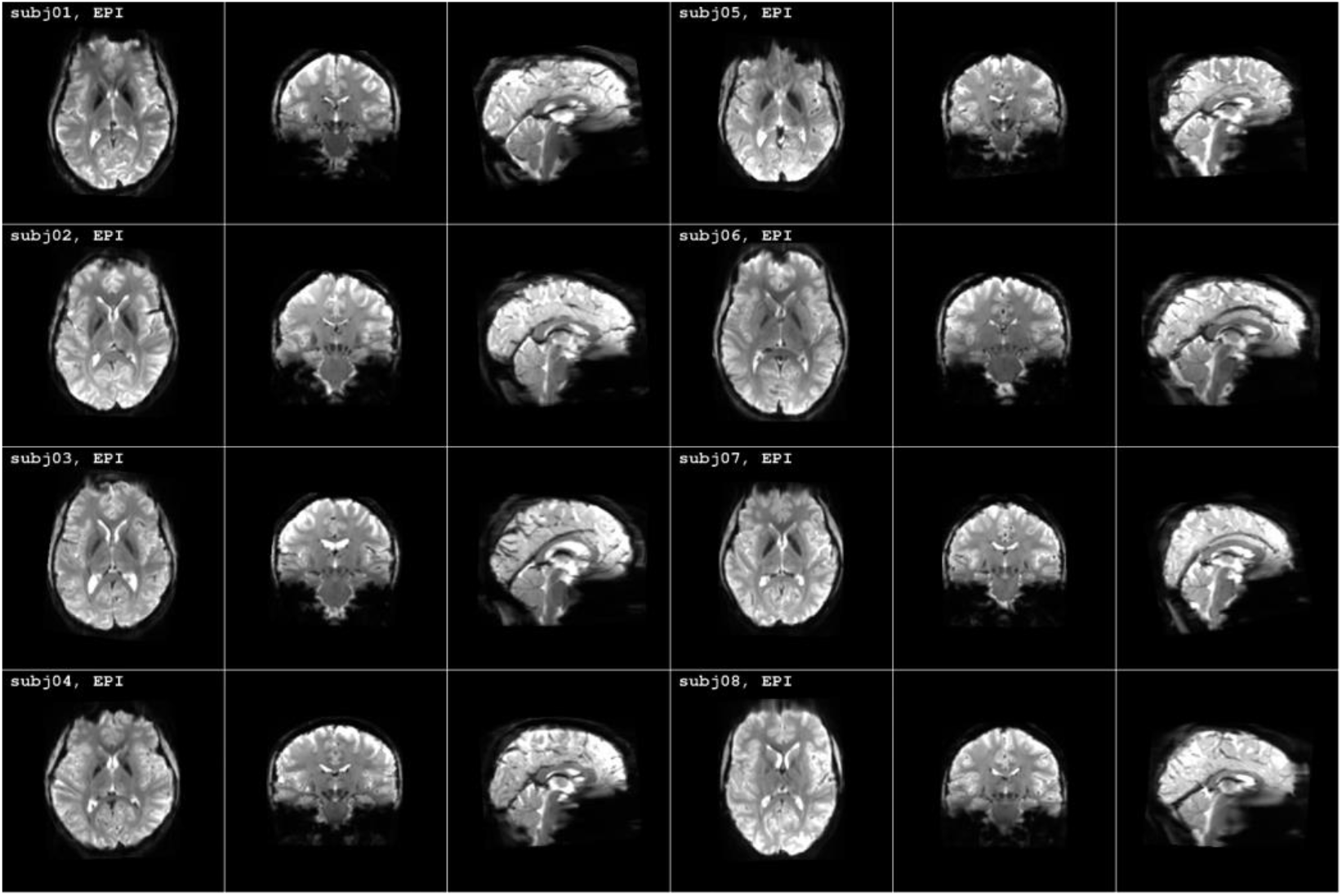
Inspection of image quality and co-registration quality. Videos available online (https://osf.io/tg5dw/,https://osf.io/g86ep/, https://osf.io/s7b2a/). Three videos are provided. One video cycles between the *T*_1_, *T*_2_, and EPI volumes, another cycles between the *T*_2_ and SWI volumes, and the third cycles between the *T*_1_ and TOF volumes. All volumes have been transformed to a common anatomical space (set by the *T*_1_ volume) in the course of data pre-processing.

**Supplementary Video 2.**
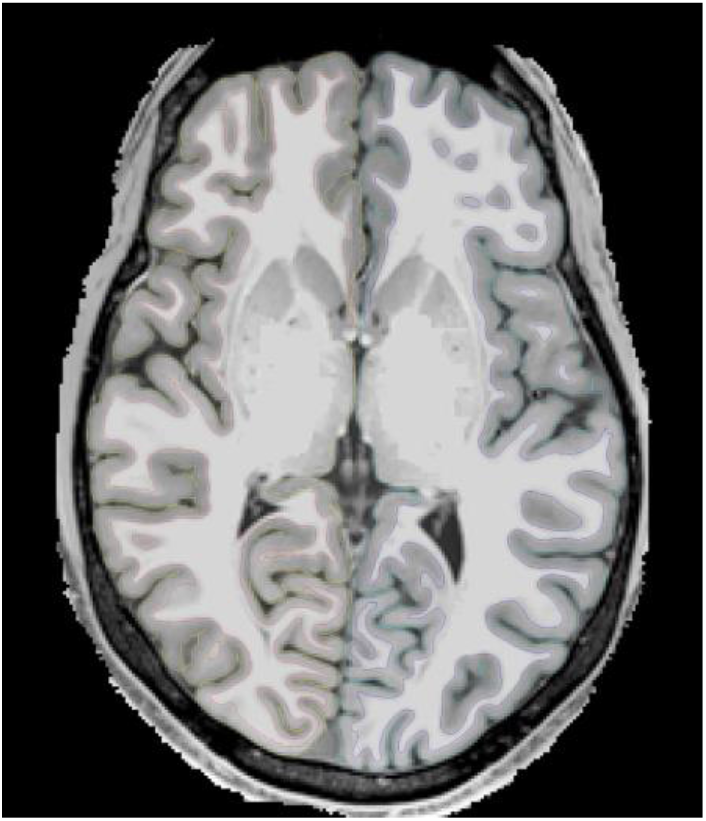
Inspection of cortical surfaces. Videos available online (https://osf.io/zyb3t/; see files named subj{01–08}_{axial,coronal,sagittal}.mp4). These videos show the FreeSurfer cortical surface reconstructions superimposed on the *T*_1_ volume. Left hemisphere white and pial surfaces are colored blue and cyan, respectively; right hemisphere white and pial surfaces are colored red and yellow, respectively. Blue voxels indicate locations that have been judged to have surface imperfections.

**Supplementary Video 3.**
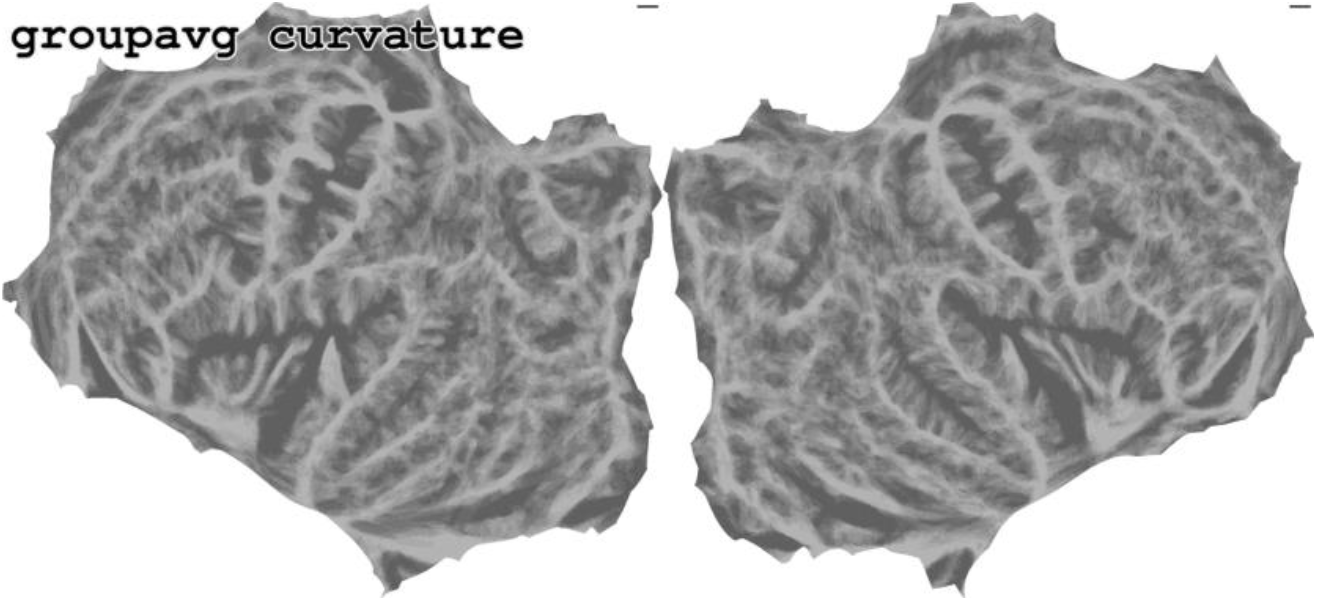
Inspection of fsaverage alignment. Video available online (https://osf.io/gh5bs/). This video cycles through (i) the binarized curvature of each of the NSD subjects mapped via nearest-neighbor interpolation to *fsaverage*, (ii) the average of this binarized curvature across subjects, and (iii) the *fsaverage* binarized curvature. The video is useful for assessing the quality of the folding-based alignment performed by FreeSurfer.

**Supplementary Video 4.**
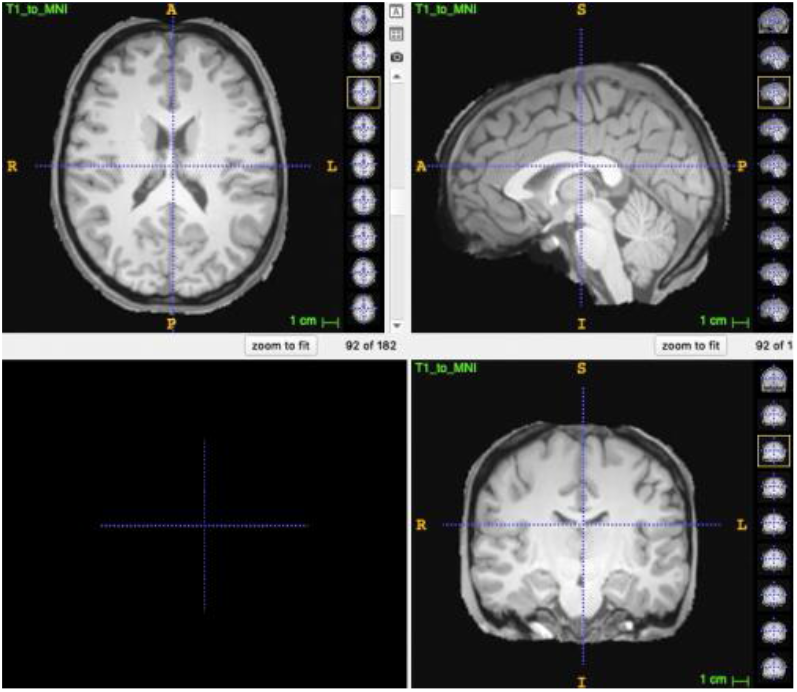
Inspection of MNI alignment. Video available online (https://osf.io/p3zqm/). This video cycles through the *T*_1_ volumes of the NSD subjects after nonlinear warping to MNI space and the MNI template volume. The video is useful for assessing the quality of the nonlinear volume-based alignment.

**Supplementary Video 5.**
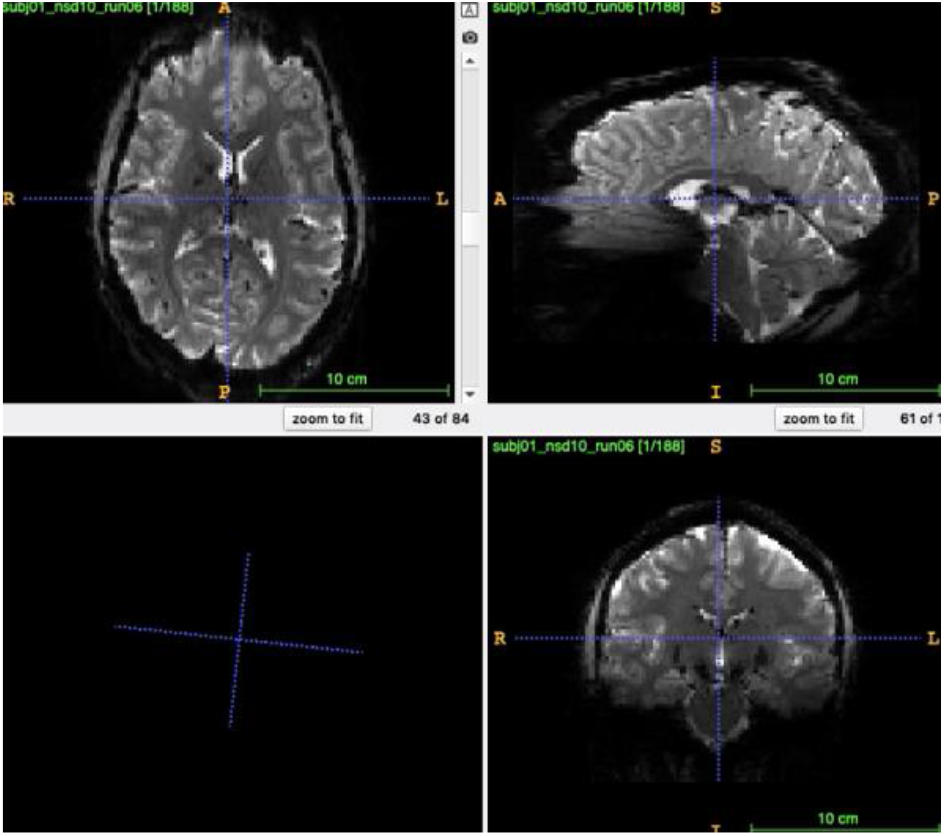
Inspection of raw and pre-processed EPI volumes. Videos available online (https://osf.io/zyb3t/; see files named subj{01–08}_nsd10_run06_{raw,pp}.mp4). These videos quickly scroll through all EPI volumes in a sample run. This is useful for assessing quality and stability of the functional imaging.

**Supplementary Video 6.**
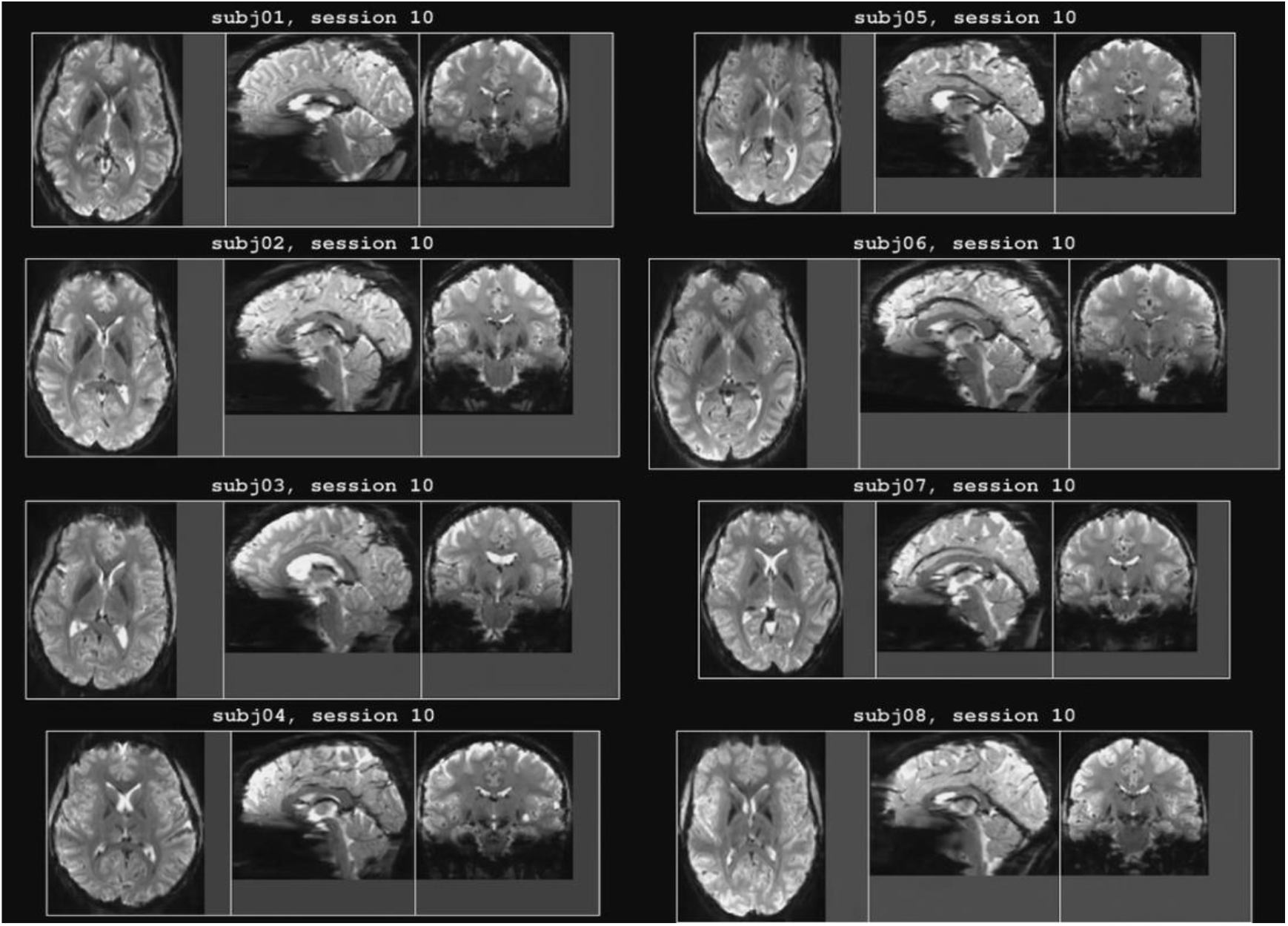
Inspection of mean EPI across scan sessions (volume visualization). Video available online (https://osf.io/ydf9j/). Each frame shows the mean EPI volume from a single scan session (1-mm data preparation). Note that session 0 corresponds to the prffloc scan session and the last two scan sessions from each subject correspond to the nsdsynthetic and nsdimagery scan sessions. This video is useful for assessing overall image quality and the stability of functional imaging across scan sessions. For quantitative analysis, see **Supplementary Figure 5**.

**Supplementary Video 7.**
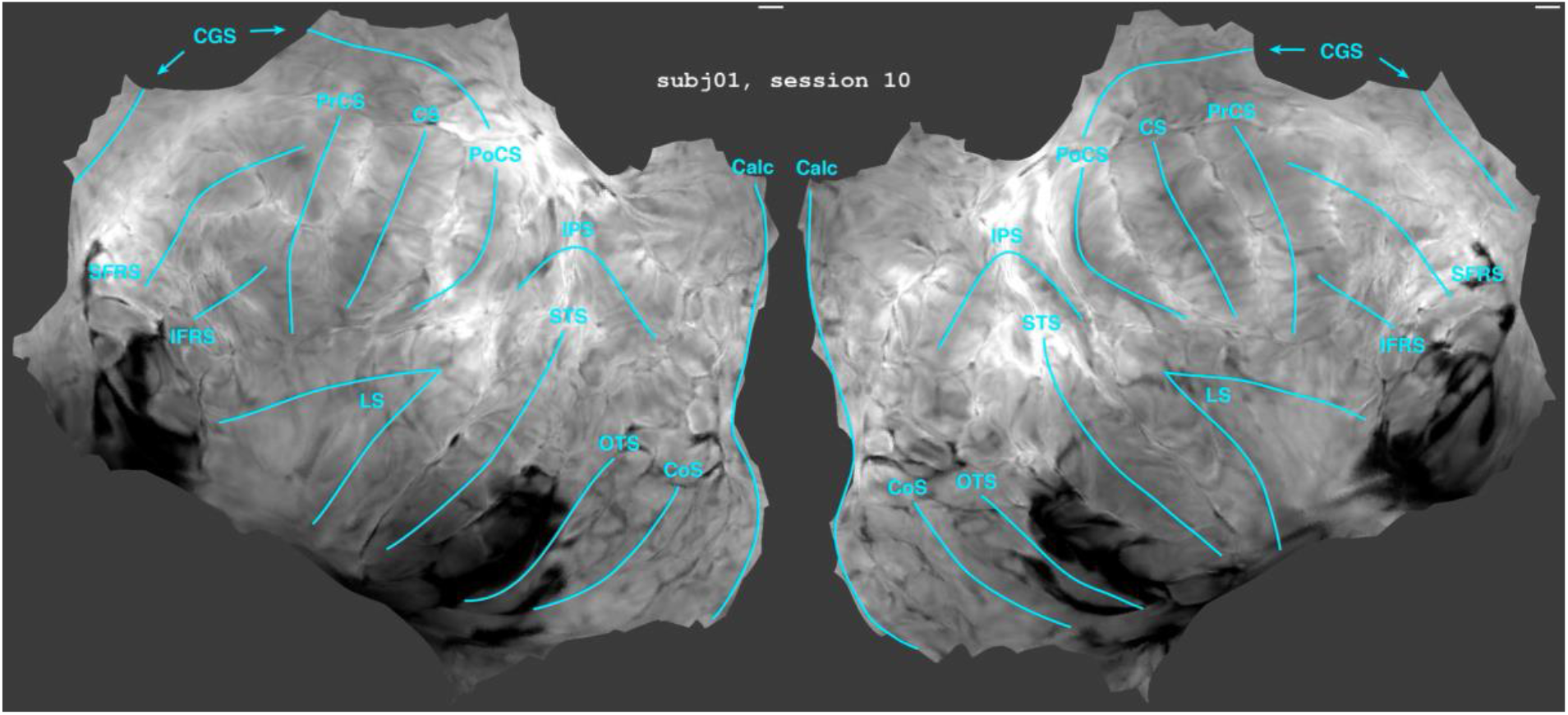
Inspection of mean EPI across scan sessions (surface visualization). Video available online (https://osf.io/ytjk4/). This is similar in spirit to **Supplementary Video 6**, except that the mean EPI volumes have been projected onto each subject’s cortical surface and then transferred to the *fsaverage* surface.

**Supplementary Video 8.**
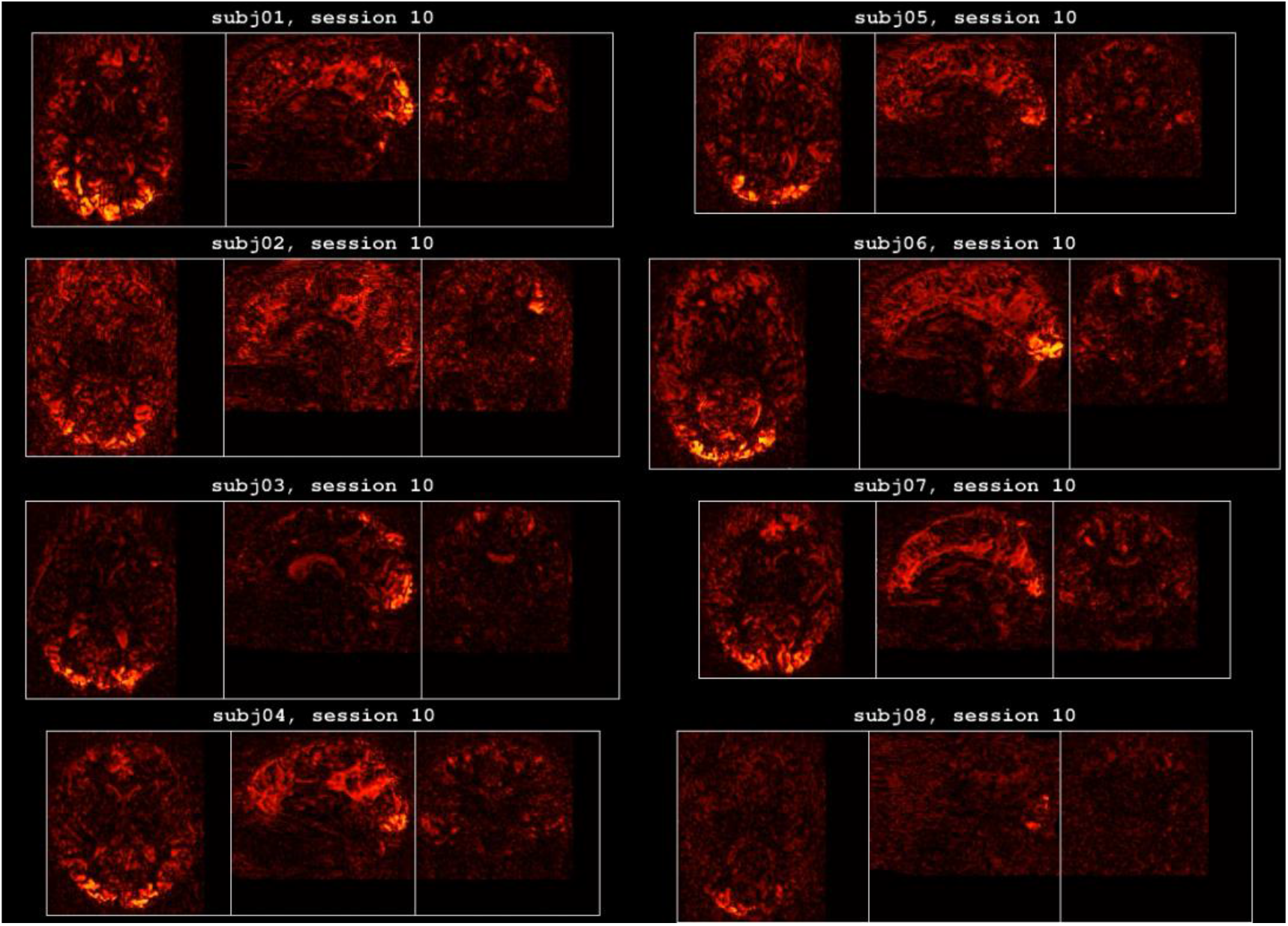
Inspection of BOLD signal strength across scan sessions (volume visualization). Video available online (https://osf.io/kwxta/). Each frame shows the amount of variance explained by the ON-OFF GLM model (1-mm data preparation; fixed color range). This video is useful for assessing the overall strength and stability of BOLD responses in the NSD dataset.

**Supplementary Video 9.**
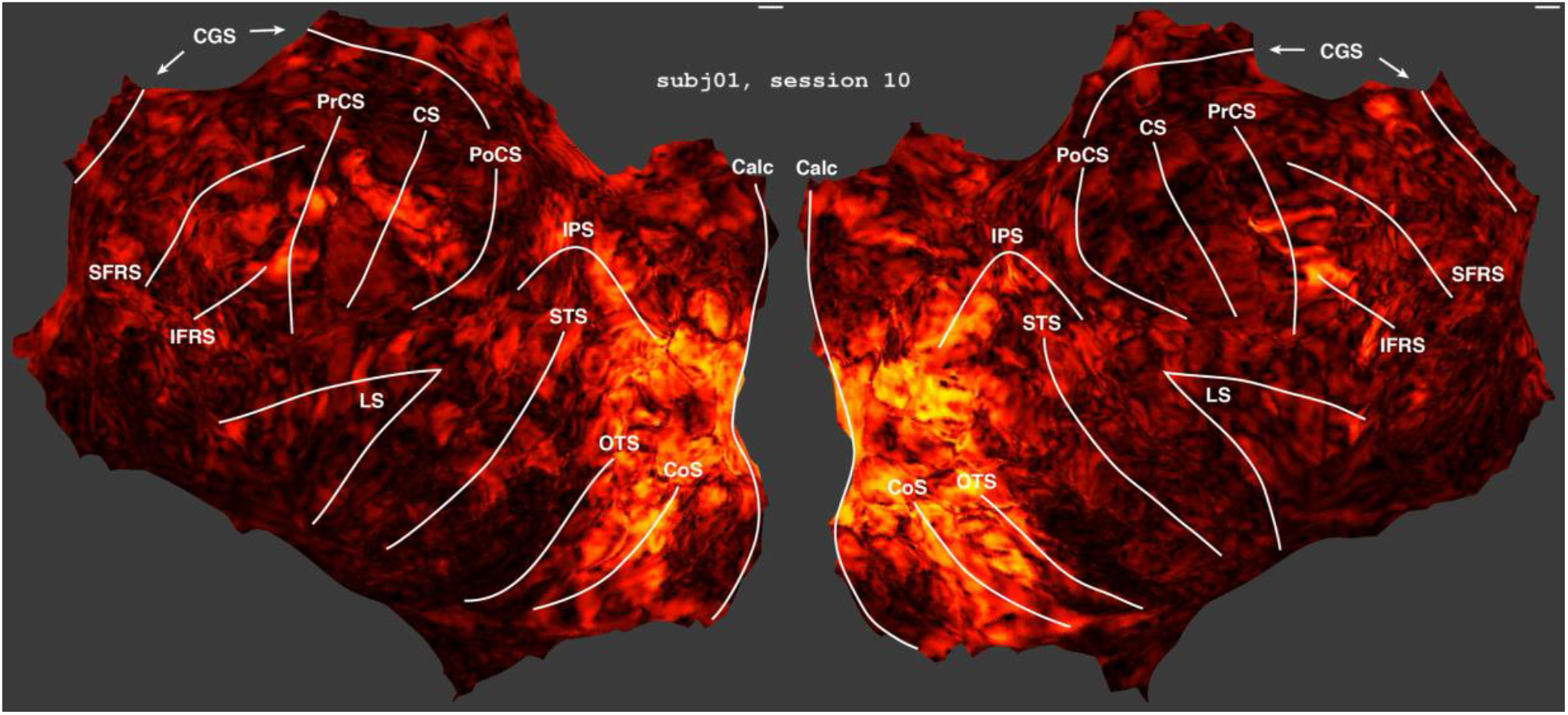
Inspection of BOLD signal strength across scan sessions (surface visualization). Video available online (https://osf.io/gu9wx/). This is similar in spirit to **Supplementary Video 8**, except that the variance explained volumes have been projected onto each subject’s cortical surface and then transferred to the *fsaverage* surface.

**Supplementary Video 10.**
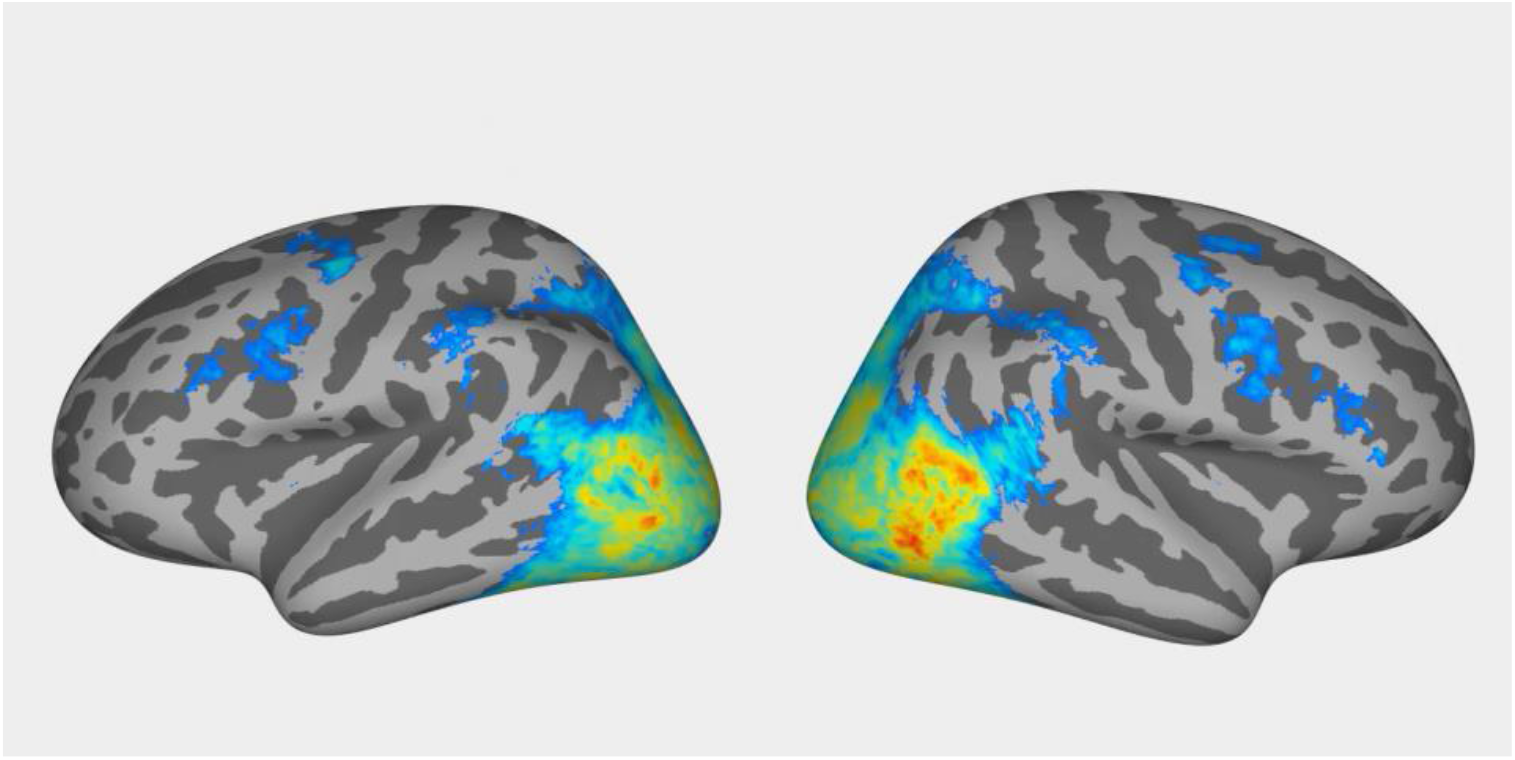
Inflated surface visualization of noise ceilings. Video available online (https://osf.io/z3wxn/). This video shows the group-average b3 noise ceiling (see **Figure 5F**) on a rotating, inflated *fsaverage* surface. Values below 15% are thresholded away in order to show the underlying curvature. This video is useful for identifying brain regions whose activity is strongly related to the sensory content presented in the NSD experiment.

## References

Ades-Aron, B., Veraart, J., Kochunov, P., McGuire, S., Sherman, P., Kellner, E., Novikov, D.S., Fieremans, E., 2018. Evaluation of the accuracy and precision of the diffusion parameter EStImation with Gibbs and NoisE removal pipeline. Neuroimage 183, 532–543. https://doi.org/10.1016/j.neuroimage.2018.07.066

Albrecht, D.G., Hamilton, D.B., 1982. Striate cortex of monkey and cat: contrast response function. Journal of neurophysiolog y 48, 217–237.

Aliko, S., Huang, J., Gheorghiu, F., Meliss, S., Skipper, J.I., 2020. A naturalistic neuroimaging database for understanding the brain using ecological stimuli. Sci Data 7, 347. https://doi.org/10.1038/s41597-020-00680-2

Arcaro, M.J., Pinsk, M.A., Kastner, S., 2015. The Anatomical and Functional Organization of the Human Visual Pulvinar. J. Neurosci. 35, 9848–9871. https://doi.org/10.1523/JNEUROSCI.1575-14.2015

Arora, S., Liang, Y., Ma, T., 2017. A Simple but Tough-to-Beat Baseline for Sentence Embeddings. Presented at the ICLR 2017.

Avants, B.B., Tustison, N.J., Song, G., Cook, P.A., Klein, A., Gee, J.C., 2011. A reproducible evaluation of ANTs similarity metric performance in brain image registration. Neuroimage 54, 2033–2044. https://doi.org/10.1016/j.neuroimage.2010.09.025

Avesani, P., McPherson, B., Hayashi, S., Caiafa, C.F., Henschel, R., Garyfallidis, E., Kitchell, L., Bullock, D., Patterson, A., Olivetti, E., Sporns, O., Saykin, A.J., Wang, L., Dinov, I., Hancock, D., Caron, B., Qian, Y., Pestilli, F., 2019. The open diffusion data derivatives, brain data upcycling via integrated publishing of derivatives and reproducible open cloud services. Scientific Data 6, 69. https://doi.org/10.1038/s41597-019-0073-y

Bassett, D.S., Sporns, O., 2017. Network neuroscience. Nature Neuroscience 20, 353–364. https://doi.org/10.1038/nn.4502

Bellec, P., Boyle, J.A., 2019. Bridging the gap between perception and action: the case for neuroimaging, AI and video games (preprint). PsyArXiv. https://doi.org/10.31234/osf.io/3epws

Benson, N.C., Jamison, K.W., Arcaro, M.J., Vu, A.T., Glasser, M.F., Coalson, T.S., Van Essen, D.C., Yacoub, E., Ugurbil, K., Winawer, J., Kay, K., 2018. The Human Connectome Project 7 Tesla retinotopy dataset: Description and population receptive field analysis. J Vis 18. https://doi.org/10.1167/18.13.23

Berron, D., Vieweg, P., Hochkeppler, A., Pluta, J.B., Ding, S.-L., Maass, A., Luther, A., Xie, L., Das, S.R., Wolk, D.A., Wolbers, T., Yushkevich, P.A., Düzel, E., Wisse, L.E.M., 2017. A protocol for manual segmentation of medial temporal lobe subregions in 7 Tesla MRI. Neuroimage Clin 15, 466–482. https://doi.org/10.1016/j.nicl.2017.05.022

Biswal, B., Yetkin, F.Z., Haughton, V.M., Hyde, J.S., 1995. Functional connectivity in the motor cortex of resting human brain using echo-planar MRI. Magn Reson Med 34, 537–541. https://doi.org/10.1002/mrm.1910340409

Boynton, G.M., Demb, J.B., Glover, G.H., Heeger, D.J., 1999. Neuronal basis of contrast discrimination. Vision research 39, 257– 269.

Brady, T.F., Konkle, T., Alvarez, G.A., Oliva, A., 2008. Visual long-term memory has a massive storage capacity for object details. PNAS 105, 14325–14329. https://doi.org/10.1073/pnas.0803390105

Brainard, D.H., 1997. The Psychophysics Toolbox. Spat Vis 10, 433–436.

Breedlove, J.L., St-Yves, G., Olman, C.A., Naselaris, T., 2020. Generative Feedback Explains Distinct Brain Activity Codes for Seen and Mental Images. Curr Biol 30, 2211-2224.e6. https://doi.org/10.1016/j.cub.2020.04.014

Bullock, D., Takemura, H., Caiafa, C.F., Kitchell, L., McPherson, B., Caron, B., Pestilli, F., 2019. Associative white matter connecting the dorsal and ventral posterior human cortex. Brain Struct Funct 224, 2631–2660. https://doi.org/10.1007/s00429-019-01907-8

Caesar, H., Uijlings, J., Ferrari, V., 2018. COCO-Stuff: Thing and Stuff Classes in Context. Presented at the Proceedings of the IEEE Conference on Computer Vision and Pattern Recognition, pp. 1209–1218.

Chang, N., Pyles, J.A., Marcus, A., Gupta, A., Tarr, M.J., Aminoff, E.M., 2019. BOLD5000, a public fMRI dataset while viewing 5000 visual images. Sci Data 6, 49. https://doi.org/10.1038/s41597-019-0052-3

Charest, I., Kriegeskorte, N., Kay, K.N., 2018. GLMdenoise improves multivariate pattern analysis of fMRI data. NeuroImage 183, 606–616. https://doi.org/10.1016/j.neuroimage.2018.08.064

Cichy, R.M., Roig, G., Oliva, A., 2019. The Algonauts Project. Nature Machine Intelligence 1, 613–613. https://doi.org/10.1038/s42256-019-0127-z

Cox, D.D., Dean, T., 2014. Neural Networks and Neuroscience-Inspired Computer Vision. Current Biology 24, R921–R929. https://doi.org/10.1016/j.cub.2014.08.026

Daducci, A., Canales-Rodríguez, E.J., Zhang, H., Dyrby, T.B., Alexander, D.C., Thiran, J.-P., 2015. Accelerated Microstructure Imaging via Convex Optimization (AMICO) from diffusion MRI data. NeuroImage 105, 32–44. https://doi.org/10.1016/j.neuroimage.2014.10.026

de Vries, S.E.J., Lecoq, J.A., Buice, M.A., Groblewski, P.A., Ocker, G.K., Oliver, M., Feng, D., Cain, N., Ledochowitsch, P., Millman, D., Roll, K., Garrett, M., Keenan, T., Kuan, L., Mihalas, S., Olsen, S., Thompson, C., Wakeman, W., Waters, J., Williams, D., Barber, C., Berbesque, N., Blanchard, B., Bowles, N., Caldejon, S.D., Casal, L., Cho, A., Cross, S., Dang, C., Dolbeare, T., Edwards, M., Galbraith, J., Gaudreault, N., Gilbert, T.L., Griffin, F., Hargrave, P., Howard, R., Huang, L., Jewell, S., Keller, N., Knoblich, U., Larkin, J.D., Larsen, R., Lau, C., Lee, E., Lee, F., Leon, A., Li, L., Long, F., Luviano, J., Mace, K., Nguyen, T., Perkins, J., Robertson, M., Seid, S., Shea-Brown, E., Shi, J., Sjoquist, N., Slaughterbeck, C., Sullivan, D., Valenza, R., White, C., Williford, A., Witten, D.M., Zhuang, J., Zeng, H., Farrell, C., Ng, L., Bernard, A., Phillips, J.W., Reid, R.C., Koch, C., 2020. A large-scale standardized physiological survey reveals functional organization of the mouse visual cortex. Nat Neurosci 23, 138–151. https://doi.org/10.1038/s41593-019-0550-9

Deng, J., Dong, W., Socher, R., Li, L., Kai Li, Li Fei-Fei, 2009. ImageNet: A large-scale hierarchical image database, in: 2009 IEEE Conference on Computer Vision and Pattern Recognition. Presented at the 2009 IEEE Conference on Computer Vision and Pattern Recognition, pp. 248–255. https://doi.org/10.1109/CVPR.2009.5206848

Duchaine, B., Nakayama, K., 2006. The Cambridge Face Memory Test: Results for neurologically intact individuals and an investigation of its validity using inverted face stimuli and prosopagnosic participants. Neuropsychologia 44, 576–585. https://doi.org/10.1016/j.neuropsychologia.2005.07.001

Esteban, O., Markiewicz, C.J., Blair, R.W., Moodie, C.A., Isik, A.I., Erramuzpe, A., Kent, J.D., Goncalves, M., DuPre, E., Snyder, M., Oya, H., Ghosh, S.S., Wright, J., Durnez, J., Poldrack, R.A., Gorgolewski, K.J., 2019. fMRIPrep: a robust preprocessing pipeline for functional MRI. Nat Methods 16, 111–116. https://doi.org/10.1038/s41592-018-0235-4

Filevich, E., Lisofsky, N., Becker, M., Butler, O., Lochstet, M., Martensson, J., Wenger, E., Lindenberger, U., Kühn, S., 2017. Day2day: investigating daily variability of magnetic resonance imaging measures over half a year. BMC Neuroscience 18, 65. https://doi.org/10.1186/s12868-017-0383-y

Fong, R.C., Scheirer, W.J., Cox, D.D., 2018. Using human brain activity to guide machine learning. Scientific Reports 8, 5397. https://doi.org/10.1038/s41598-018-23618-6

Frey, M., Nau, M., Doeller, C.F., 2020. MR-based camera-less eye tracking using deep neural networks. bioRxiv 2020.11.30.401323. https://doi.org/10.1101/2020.11.30.401323

Fukutomi, H., Glasser, M.F., Zhang, H., Autio, J.A., Coalson, T.S., Okada, T., Togashi, K., Van Essen, D.C., Hayashi, T., 2018. Neurite imaging reveals microstructural variations in human cerebral cortical gray matter. NeuroImage, Microstructural Imaging 182, 488–499. https://doi.org/10.1016/j.neuroimage.2018.02.017

Garyfallidis, E., Brett, M., Amirbekian, B., Rokem, A., Van Der Walt, S., Descoteaux, M., Nimmo-Smith, I., 2014. Dipy, a library for the analysis of diffusion MRI data. Front. Neuroinform. 8. https://doi.org/10.3389/fninf.2014.00008

Geisler, W.S., 2008. Visual perception and the statistical properties of natural scenes. Annual review of psychology 59, 167–192.

Glasser, M.F., Coalson, T.S., Robinson, E.C., Hacker, C.D., Harwell, J., Yacoub, E., Ugurbil, K., Andersson, J., Beckmann, C.F., Jenkinson, M., Smith, S.M., Van Essen, D.C., 2016. A multi-modal parcellation of human cerebral cortex. Nature 536, 171–178. https://doi.org/10.1038/nature18933

Gomez, J., Natu, V., Jeska, B., Barnett, M., Grill-Spector, K., 2018. Development differentially sculpts receptive fields across early and high-level human visual cortex. Nature Communications 9, 788. https://doi.org/10.1038/s41467-018-03166-3

Gonzalez-Castillo, J., Saad, Z.S., Handwerker, D.A., Inati, S.J., Brenowitz, N., Bandettini, P.A., 2012. Whole-brain, time-locked activation with simple tasks revealed using massive averaging and model-free analysis. PNAS 109, 5487–5492. https://doi.org/10.1073/pnas.1121049109

Gordon, E.M., Laumann, T.O., Gilmore, A.W., Newbold, D.J., Greene, D.J., Berg, J.J., Ortega, M., Hoyt-Drazen, C., Gratton, C., Sun, H., Hampton, J.M., Coalson, R.S., Nguyen, A.L., McDermott, K.B., Shimony, J.S., Snyder, A.Z., Schlaggar, B.L., Petersen, S.E., Nelson, S.M., Dosenbach, N.U.F., 2017. Precision Functional Mapping of Individual Human Brains. Neuron 95, 791-807.e7. https://doi.org/10.1016/j.neuron.2017.07.011

Gorgolewski, K.J., Auer, T., Calhoun, V.D., Craddock, R.C., Das, S., Duff, E.P., Flandin, G., Ghosh, S.S., Glatard, T., Halch enko, Y.O., Handwerker, D.A., Hanke, M., Keator, D., Li, X., Michael, Z., Maumet, C., Nichols, B.N., Nichols, T.E., Pellman, J., Poline, J.-B., Rokem, A., Schaefer, G., Sochat, V., Triplett, W., Turner, J.A., Varoquaux, G., Poldrack, R.A., 2016. The brain imaging data structure, a format for organizing and describing outputs of neuroimaging experiments. Sci Data 3, 1–9. https://doi.org/10.1038/sdata.2016.44

Gratton, C., Dworetsky, A., Coalson, R.S., Adeyemo, B., Laumann, T.O., Wig, G.S., Kong, T.S., Gratton, G., Fabiani, M., Barch, D.M., Tranel, D., Miranda-Dominguez, O., Fair, D.A., Dosenbach, N.U.F., Snyder, A.Z., Perlmutter, J.S., Petersen, S.E., Campbell, M.C., 2020. Removal of high frequency contamination from motion estimates in single-band fMRI saves data without biasing functional connectivity. Neuroimage 217, 116866. https://doi.org/10.1016/j.neuroimage.2020.116866

Grill-Spector, K., Malach, R., 2004. The human visual cortex. Annual review of neuroscience 27, 649–677.

Güçlü, U., van Gerven, M.A.J., 2015. Deep Neural Networks Reveal a Gradient in the Complexity of Neural Representations across the Ventral Stream. J. Neurosci. 35, 10005–10014. https://doi.org/10.1523/JNEUROSCI.5023-14.2015

Hagmann, P., Cammoun, L., Gigandet, X., Meuli, R., Honey, C.J., Wedeen, V.J., Sporns, O., 2008. Mapping the Structural Core of Human Cerebral Cortex. PLOS Biology 6, e159. https://doi.org/10.1371/journal.pbio.0060159

Han, K., Wen, H., Shi, J., Lu, K.-H., Zhang, Y., Fu, D., Liu, Z., 2019. Variational autoencoder: An unsupervised model for encoding and decoding fMRI activity in visual cortex. NeuroImage 198, 125–136. https://doi.org/10.1016/j.neuroimage.2019.05.039

Handwerker, D.A., Gonzalez-Castillo, J., D’Esposito, M., Bandettini, P.A., 2012. The continuing challenge of understanding and modeling hemodynamic variation in fMRI. NeuroImage 62, 1017–1023. https://doi.org/10.1016/j.neuroimage.2012.02.015

Hoerl, A.E., Kennard, R.W., 1970. Ridge Regression: Biased Estimation for Nonorthogonal Problems. null 12, 55–67. https://doi.org/10.1080/00401706.1970.10488634

Horikawa, T., Kamitani, Y., 2017. Generic decoding of seen and imagined objects using hierarchical visual features. Nature Communications 8, 15037. https://doi.org/10.1038/ncomms15037

Huth, A.G., Nishimoto, S., Vu, A.T., Gallant, J.L., 2012. A continuous semantic space describes the representation of thousands of object and action categories across the human brain. Neuron 76, 1210–1224. https://doi.org/10.1016/j.neuron.2012.10.014

Jensen, J.H., Helpern, J.A., 2010. MRI quantification of non-Gaussian water diffusion by kurtosis analysis. NMR in Biomedicine 23, 698–710. https://doi.org/10.1002/nbm.1518

Jezzard, P., 2012. Correction of geometric distortion in fMRI data. NeuroImage 62, 648–651. https://doi.org/10.1016/j.neuroimage.2011.09.010

Jonathan Pillow, null Sahani, M., 2019. Editorial overview: Machine learning, big data, and neuroscience. Curr Opin Neurobiol 55, iii–iv. https://doi.org/10.1016/j.conb.2019.05.002

Kang, X., Yund, E.W., Herron, T.J., Woods, D.L., 2007. Improving the resolution of functional brain imaging: analyz ing functional data in anatomical space. Magnetic resonance imaging 25, 1070–1078.

Kay, K., Jamison, K.W., Vizioli, L., Zhang, R., Margalit, E., Ugurbil, K., 2019. A critical assessment of data quality and ve nous effects in sub-millimeter fMRI. NeuroImage 189, 847–869. https://doi.org/10.1016/j.neuroimage.2019.02.006

Kay, K., Jamison, K.W., Zhang, R.-Y., Uğurbil, K., 2020. A temporal decomposition method for identifying venous effects in task-based fMRI. Nat Methods 17, 1033–1039. https://doi.org/10.1038/s41592-020-0941-6

Kay, K.N., Naselaris, T., Prenger, R.J., Gallant, J.L., 2008. Identifying natural images from human brain activity. Nature 452, 352– 355. https://doi.org/10.1038/nature06713

Kay, K.N., Rokem, A., Winawer, J., Dougherty, R.F., Wandell, B., 2013a. GLMdenoise: a fast, automated technique for denoising task-based fMRI data. Front Neurosci 7, 247. https://doi.org/10.3389/fnins.2013.00247

Kay, K.N., Weiner, K.S., Grill-Spector, K., 2015. Attention reduces spatial uncertainty in human ventral temporal cortex. Curr Biol 25, 595–600. https://doi.org/10.1016/j.cub.2014.12.050

Kay, K.N., Winawer, J., Mezer, A., Wandell, B., 2013b. Compressive spatial summation in human visual cortex. Journal of neurophysiology 110, 481–494. https://doi.org/10.1152/jn.00105.2013

Kay, K.N., Yeatman, J.D., 2017. Bottom-up and top-down computations in word- and face-selective cortex. Elife 6, e22341. https://doi.org/10.7554/eLife.22341

Khaligh-Razavi, S.-M., Kriegeskorte, N., 2014. Deep supervised, but not unsupervised, models may explain IT cortical representation. PLoS computational biology 10, e1003915. https://doi.org/10.1371/journal.pcbi.1003915

Kingma, D.P., Ba, J., 2017. Adam: A Method for Stochastic Optimization. arXiv:1412.6980 [cs].

Kriegeskorte, N., Mur, M., 2012. Inverse MDS: Inferring Dissimilarity Structure from Multiple Item Arrangements. Front Psychol 3, 245. https://doi.org/10.3389/fpsyg.2012.00245

Kriegeskorte, N., Mur, M., Bandettini, P., 2008. Representational similarity analysis - connecting the branches of systems neuroscience. Frontiers in systems neuroscience 2, 4.

Krizhevsky, A., 2009. Learning Multiple Layers of Features from Tiny Images. University of Toronto.

Krizhevsky, A., Sutskever, I., Hinton, G.E., 2012. ImageNet Classification with Deep Convolutional Neural Networks 1097–1105.

Krizhevsky, A., Sutskever, I., Hinton, G.E., 2017. ImageNet classification with deep convolutional neural networks. Commun. ACM 60, 84–90. https://doi.org/10.1145/3065386

Lage-Castellanos, A., Valente, G., Formisano, E., De Martino, F., 2019. Methods for computing the maximum performance of computational models of fMRI responses. PLoS Comput Biol 15, e1006397. https://doi.org/10.1371/journal.pcbi.1006397

Laumann, T.O., Gordon, E.M., Adeyemo, B., Snyder, A.Z., Joo, S.J., Chen, M.-Y., Gilmore, A.W., McDermott, K.B., Nelson, S.M., Dosenbach, N.U.F., Schlaggar, B.L., Mumford, J.A., Poldrack, R.A., Petersen, S.E., 2015. Functional System and Areal Organization of a Highly Sampled Individual Human Brain. Neuron 87, 657–670. https://doi.org/10.1016/j.neuron.2015.06.037

Lin, T.-Y., Maire, M., Belongie, S., Hays, J., Pietro Perona, Ramanan, D., Dollár, P., Zitnick, C.L., 2014. Microsoft COCO: Common Objects in Context, in: Computer Vision – ECCV 2014. Springer, Cham, Cham, pp. 740–755. https://doi.org/10.1007/978-3-319-10602-1_48

Lynch, C.J., Silver, B.M., Dubin, M.J., Martin, A., Voss, H.U., Jones, R.M., Power, J.D., 2020. Prevalent and sex-biased breathing patterns modify functional connectivity MRI in young adults. Nature Communications 11, 5290. https://doi.org/10.1038/s41467-020-18974-9

Maaten, L. van der, Hinton, G., 2008. Visualizing Data using t-SNE. Journal of Machine Learning Research 9, 2579–2605.

Markram, H., Muller, E., Ramaswamy, S., Reimann, M.W., Abdellah, M., Sanchez, C.A., Ailamaki, A., Alonso-Nanclares, L., Antille, N., Arsever, S., Kahou, G.A.A., Berger, T.K., Bilgili, A., Buncic, N., Chalimourda, A., Chindemi, G., Courcol, J.-D., Delalondre, F., Delattre, V., Druckmann, S., Dumusc, R., Dynes, J., Eilemann, S., Gal, E., Gevaert, M.E., Ghobril, J.-P., Gidon, A., Graham, J.W., Gupta, A., Haenel, V., Hay, E., Heinis, T., Hernando, J.B., Hines, M., Kanari, L., Keller, D., Kenyon, J., Khazen, G., Kim, Y., King, J.G., Kisvarday, Z., Kumbhar, P., Lasserre, S., Le Bé, J.-V., Magalhães, B.R.C., Merchán-Pérez, A., Meystre, J., Morrice, B.R., Muller, J., Muñoz-Céspedes, A., Muralidhar, S., Muthurasa, K., Nachbaur, D., Newton, T.H., Nolte, M., Ovcharenko, A., Palacios, J., Pastor, L., Perin, R., Ranjan, R., Riachi, I., Rodríguez, J.-R., Riquelme, J.L., Rössert, C., Sfyrakis, K., Shi, Y., Shillcock, J.C., Silberberg, G., Silva, R., Tauheed, F., Telefont, M., Toledo-Rodriguez, M., Tränkler, T., Van Geit, W., Díaz, J.V., Walker, R., Wang, Y., Zaninetta, S.M., DeFelipe, J., Hill, S.L., Segev, I., Schürmann, F., 2015. Reconstruction and Simulation of Neocortical Microcircuitry. Cell 163, 456–492. https://doi.org/10.1016/j.cell.2015.09.029

Marks, D.F., 1973. Visual Imagery Differences in the Recall of Pictures. British Journal of Psychology 64, 17–24. https://doi.org/10.1111/j.2044-8295.1973.tb01322.x

Naselaris, T., Allen, E., Kay, K., 2021. Extensive sampling for complete models of individual brains. Current Opinion in Beha vioral Sciences 40, 45–51. https://doi.org/10.1016/j.cobeha.2020.12.008

Naselaris, T., Bassett, D.S., Fletcher, A.K., Kording, K., Kriegeskorte, N., Nienborg, H., Poldrack, R.A., Shohamy, D., Kay, K., 2018. Cognitive Computational Neuroscience: A New Conference for an Emerging Discipline. Trends Cogn Sci 22, 365–367. https://doi.org/10.1016/j.tics.2018.02.008

Naselaris, T., Kay, K.N., Nishimoto, S., Gallant, J.L., 2011. Encoding and decoding in fMRI. NeuroImage 56, 400 –410. https://doi.org/10.1016/j.neuroimage.2010.07.073

Nastase, S.A., Liu, Y.-F., Hillman, H., Norman, K.A., Hasson, U., 2020. Leveraging shared connectivity to aggregate heterogeneous datasets into a common response space. Neuroimage 217, 116865. https://doi.org/10.1016/j.neuroimage.2020.116865

Nili, H., Wingfield, C., Walther, A., Su, L., Marslen-Wilson, W., Kriegeskorte, N., 2014. A toolbox for representational similarity analysis. PLoS computational biology 10, e1003553. https://doi.org/10.1371/journal.pcbi.1003553

Pedregosa, F., Varoquaux, G., Gramfort, A., Michel, V., Thirion, B., Grisel, O., Blondel, M., Prettenhofer, P., Weiss, R., Dubourg, V., Vanderplas, J., Passos, A., Cournapeau, D., Brucher, M., Perrot, M., Duchesnay, É., 2011. Scikit-learn: Machine Learning in Python. Journal of Machine Learning Research 12, 2825–2830.

Pelli, D.G., 1997. The VideoToolbox software for visual psychophysics: transforming numbers into movies. Spat Vis 10, 437–442.

Pierpaoli, C., Jezzard, P., Basser, P.J., Barnett, A., Di Chiro, G., 1996. Diffusion tensor MR imaging of the human brain. Radiology 201, 637–648. https://doi.org/10.1148/radiology.201.3.8939209

Pinho, A.L., Amadon, A., Ruest, T., Fabre, M., Dohmatob, E., Denghien, I., Ginisty, C., Becuwe-Desmidt, S., Roger, S., Laurier, L., Joly-Testault, V., Médiouni-Cloarec, G., Doublé, C., Martins, B., Pinel, P., Eger, E., Varoquaux, G., Pallier, C., Dehaene, S., Hertz-Pannier, L., Thirion, B., 2018. Individual Brain Charting, a high-resolution fMRI dataset for cognitive mapping. Sci Data 5, 180105. https://doi.org/10.1038/sdata.2018.105

Poldrack, R.A., Laumann, T.O., Koyejo, O., Gregory, B., Hover, A., Chen, M.-Y., Gorgolewski, K.J., Luci, J., Joo, S.J., Boyd, R.L., Hunicke-Smith, S., Simpson, Z.B., Caven, T., Sochat, V., Shine, J.M., Gordon, E., Snyder, A.Z., Adeyemo, B., Petersen, S. E., Glahn, D.C., Reese Mckay, D., Curran, J.E., Göring, H.H.H., Carless, M.A., Blangero, J., Dougherty, R., Leemans, A., Handwerker, D.A., Frick, L., Marcotte, E.M., Mumford, J.A., 2015. Long-term neural and physiological phenotyping of a single human. Nat Commun 6, 8885. https://doi.org/10.1038/ncomms9885

Polimeni, J.R., Renvall, V., Zaretskaya, N., Fischl, B., 2018. Analysis strategies for high-resolution UHF-fMRI data. NeuroImage 168, 296–320. https://doi.org/10.1016/j.neuroimage.2017.04.053

Power, J.D., Barnes, K.A., Snyder, A.Z., Schlaggar, B.L., Petersen, S.E., 2012. Spurious but systematic correlations in functional connectivity MRI networks arise from subject motion. Neuroimage 59, 2142–2154. https://doi.org/10.1016/j.neuroimage.2011.10.018

Power, J.D., Silver, B.M., Silverman, M.R., Ajodan, E.L., Bos, D.J., Jones, R.M., 2019. Customized head molds reduce motion during resting state fMRI scans. NeuroImage 189, 141–149. https://doi.org/10.1016/j.neuroimage.2019.01.016

Rokem, A., Kay, K., 2020. Fractional ridge regression: a fast, interpretable reparameterization of ridge regression. GigaScience 9. https://doi.org/10.1093/gigascience/giaa133

Roth, Z.N., Ryoo, M., Merriam, E.P., 2020. Task-related activity in human visual cortex. PLoS Biol 18, e3000921. https://doi.org/10.1371/journal.pbio.3000921

Satterthwaite, T.D., Elliott, M.A., Ruparel, K., Loughead, J., Prabhakaran, K., Calkins, M.E., Hopson, R., Jackson, C., Keefe, J., Riley, M., Mentch, F.D., Sleiman, P., Verma, R., Davatzikos, C., Hakonarson, H., Gur, R.C., Gur, R.E., 2014. Neuroimaging of the Philadelphia Neurodevelopmental Cohort. NeuroImage 86, 544–553. https://doi.org/10.1016/j.neuroimage.2013.07.064

Schira, M.M., Tyler, C.W., Breakspear, M., Spehar, B., 2009. The foveal confluence in human visual cortex. J. Neurosci. 29, 9050– 9058. https://doi.org/10.1523/JNEUROSCI.1760-09.2009

Seeliger, K., Ambrogioni, L., Güçlütürk, Y., Bulk L.M. van den, Güçlü, U., Gerven, M.A.J. van, 2021. End-to-end neural system identification with neural information flow. PLOS Computational Biology 17, e1008558. https://doi.org/10.1371/journal.pcbi.1008558

Seeliger, K., Sommers, R.P., Güçlü, U., Bosch, S.E., Gerven, M.A.J. van, 2019. A large single-participant fMRI dataset for probing brain responses to naturalistic stimuli in space and time. bioRxiv 687681. https://doi.org/10.1101/687681

Shahid, A., Wilkinson, K., Marcu, S., Shapiro, C.M., 2012. Stanford Sleepiness Scale (SSS), in: Shahid, A., Wilkinson, K., Marcu, S., Shapiro, C.M. (Eds.), STOP, THAT and One Hundred Other Sleep Scales. Springer, New York, NY, pp. 369–370. https://doi.org/10.1007/978-1-4419-9893-4_91

Siegle, J.H., Jia, X., Durand, S., Gale, S., Bennett, C., Graddis, N., Heller, G., Ramirez, T.K., Choi, H., Luviano, J.A., Groblewski, P.A., Ahmed, R., Arkhipov, A., Bernard, A., Billeh, Y.N., Brown, D., Buice, M.A., Cain, N., Caldejon, S., Casal, L., Cho, A., Chvilicek, M., Cox, T.C., Dai, K., Denman, D.J., de Vries, S.E.J., Dietzman, R., Esposito, L., Farrell, C., Feng, D., Galbraith, J., Garrett, M., Gelfand, E.C., Hancock, N., Harris, J.A., Howard, R., Hu, B., Hytnen, R., Iyer, R., Jessett, E., Johnson, K., Kato, I., Kiggins, J., Lambert, S., Lecoq, J., Ledochowitsch, P., Lee, J.H., Leon, A., Li, Y., Liang, E., Long, F., Mace, K., Melchior, J., Millman, D., Mollenkopf, T., Nayan, C., Ng, L., Ngo, K., Nguyen, T., Nicovich, P.R., North, K., Ocker, G.K., Ollerenshaw, D., Oliver, M., Pachitariu, M., Perkins, J., Reding, M., Reid, D., Robertson, M., Ronellenfitch, K., Seid, S., Slaughterbeck, C., Stoecklin, M., Sullivan, D., Sutton, B., Swapp, J., Thompson, C., Turner, K., Wakeman, W., Whitesell, J.D., Williams, D., Williford, A., Young, R., Zeng, H., Naylor, S., Phillips, J.W., Reid, R.C., Mihalas, S., Olsen, S.R., Koch, C., 2021. Survey of spiking in the mouse visual system reveals functional hierarchy. Nature. https://doi.org/10.1038/s41586-020-03171-x

Sinz, F.H., Pitkow, X., Reimer, J., Bethge, M., Tolias, A.S., 2019. Engineering a Less Artificial Intelligence. Neuron 103, 967–979. https://doi.org/10.1016/j.neuron.2019.08.034

Smith, R.E., Tournier, J.-D., Calamante, F., Connelly, A., 2012. Anatomically-constrained tractography: Improved diffusion MRI streamlines tractography through effective use of anatomical information. NeuroImage 62, 1924–1938. https://doi.org/10.1016/j.neuroimage.2012.06.005

Smith, R.E., Tournier, J.-D., Calamante, F., Connelly, A., 2015. SIFT2: Enabling dense quantitative assessment of brain white matter connectivity using streamlines tractography. NeuroImage 119, 338–351. https://doi.org/10.1016/j.neuroimage.2015.06.092

Son, J., Ai, L., Lim, R., Xu, T., Colcombe, S., Franco, A.R., Cloud, J., LaConte, S., Lisinski, J., Klein, A., Craddock, R.C., Milham, M., 2020. Evaluating fMRI-Based Estimation of Eye Gaze During Naturalistic Viewing. Cerebral Cortex 30, 1171–1184. https://doi.org/10.1093/cercor/bhz157

Spaniol, J., Davidson, P.S.R., Kim, A.S.N., Han, H., Moscovitch, M., Grady, C.L., 2009. Event-related fMRI studies of episodic encoding and retrieval: meta-analyses using activation likelihood estimation. Neuropsychologia 47, 1765–1779. https://doi.org/10.1016/j.neuropsychologia.2009.02.028

St-Yves, G., Naselaris, T., 2017. The feature-weighted receptive field: an interpretable encoding model for complex feature spaces. NeuroImage. https://doi.org/10.1016/j.neuroimage.2017.06.035

Stansbury, D.E., Naselaris, T., Gallant, J.L., 2013. Natural scene statistics account for the representation of scene categories in human visual cortex. Neuron 79, 1025–1034. https://doi.org/10.1016/j.neuron.2013.06.034

Stigliani, A., Weiner, K.S., Grill-Spector, K., 2015. Temporal Processing Capacity in High-Level Visual Cortex Is Domain Specific. J. Neurosci. 35, 12412–12424. https://doi.org/10.1523/JNEUROSCI.4822-14.2015

Stringer, C., Pachitariu, M., Steinmetz, N., Carandini, M., Harris, K.D., 2019. High-dimensional geometry of population responses in visual cortex. Nature 571, 361–365. https://doi.org/10.1038/s41586-019-1346-5

Tardif, J., Watson, M., Giaschi, D., Gosselin, F., 2016. Measuring the Contrast Sensitivity Function in just three clicks. Journal of Vision 16, 966–966. https://doi.org/10.1167/16.12.966

Taylor, J.R., Williams, N., Cusack, R., Auer, T., Shafto, M.A., Dixon, M., Tyler, L.K., Cam-Can, null, Henson, R.N., 2017. The Cambridge Centre for Ageing and Neuroscience (Cam-CAN) data repository: Structural and functional MRI, MEG, and cognitive data from a cross-sectional adult lifespan sample. Neuroimage 144, 262–269. https://doi.org/10.1016/j.neuroimage.2015.09.018

Toneva, M., Wehbe, L., 2019. Interpreting and improving natural-language processing (in machines) with natural language-processing (in the brain). arXiv:1905.11833 [cs, q-bio].

Torgesen, J.K., Wagner, R., Rashotte, C., 2012. Test of word reading efficiency:(TOWRE-2). Pearson Clinical Assessment.

Tournier, J.-D., Calamante, F., Connelly, A., 2007. Robust determination of the fibre orientation distribution in diffusion MRI: non-negativity constrained super-resolved spherical deconvolution. NeuroImage 35, 1459–1472. https://doi.org/10.1016/j.neuroimage.2007.02.016

Tournier, J.-D., Smith, R., Raffelt, D., Tabbara, R., Dhollander, T., Pietsch, M., Christiaens, D., Jeurissen, B., Yeh, C.-H., Connelly, A., 2019. MRtrix3: A fast, flexible and open software framework for medical image processing and visualisation. NeuroImage 202, 116137. https://doi.org/10.1016/j.neuroimage.2019.116137

van de Ven, G.M., Siegelmann, H.T., Tolias, A.S., 2020. Brain-inspired replay for continual learning with artificial neural networks. Nature Communications 11, 4069. https://doi.org/10.1038/s41467-020-17866-2

Van Essen, D.C., Lewis, J.W., Drury, H.A., Hadjikhani, N., Tootell, R.B., Bakircioglu, M., Miller, M.I., 2001. Mapping visual cortex in monkeys and humans using surface-based atlases. Vision Res 41, 1359–1378. https://doi.org/10.1016/s0042-6989(01)00045-1

Van Essen, D.C., Smith, S.M., Barch, D.M., Behrens, T.E.J., Yacoub, E., Ugurbil, K., WU-Minn HCP Consortium, 2013. The WU-Minn Human Connectome Project: an overview. NeuroImage 80, 62–79. https://doi.org/10.1016/j.neuroimage.2013.05.041

Vu, M.-A.T., Adali, T., Ba, D., Buzsáki, G., Carlson, D., Heller, K., Liston, C., Rudin, C., Sohal, V.S., Widge, A.S., Mayberg, H.S., Sapiro, G., Dzirasa, K., 2018. A Shared Vision for Machine Learning in Neuroscience. J Neurosci 38, 1601–1607. https://doi.org/10.1523/JNEUROSCI.0508-17.2018

Wagner, A.D., Shannon, B.J., Kahn, I., Buckner, R.L., 2005. Parietal lobe contributions to episodic memory retrieval. Trends Cogn Sci 9, 445–453. https://doi.org/10.1016/j.tics.2005.07.001

Wang, A.Y., Wehbe, L., Tarr, M.J., 2019. Neural Taskonomy: Inferring the Similarity of Task-Derived Representations from Brain Activity. bioRxiv 708016. https://doi.org/10.1101/708016

Wang, L., Mruczek, R.E.B., Arcaro, M.J., Kastner, S., 2015. Probabilistic Maps of Visual Topography in Human Cortex. Cereb. Cortex 25, 3911–3931. https://doi.org/10.1093/cercor/bhu277

Watanabe, M., Bartels, A., Macke, J.H., Murayama, Y., Logothetis, N.K., 2013. Temporal jitter of the BOLD signal reveals a reliable initial dip and improved spatial resolution. Curr Biol 23, 2146–2150. https://doi.org/10.1016/j.cub.2013.08.057

Wheeler, M.E., Petersen, S.E., Buckner, R.L., 2000. Memory’s echo: Vivid remembering reactivates sensory-specific cortex. PNAS 97, 11125–11129. https://doi.org/10.1073/pnas.97.20.11125

Winawer, J., Witthoft, N., 2017. Identification of the ventral occipital visual field maps in the human brain. F1000Res 6, 1526. https://doi.org/10.12688/f1000research.12364.1

Yamins, D.L.K., Hong, H., Cadieu, C.F., Solomon, E.A., Seibert, D., DiCarlo, J.J., 2014. Performance-optimized hierarchical models predict neural responses in higher visual cortex. Proceedings of the National Academy of Sciences of the United States of America 111, 8619–8624. https://doi.org/10.1073/pnas.1403112111

Yeatman, J.D., Dougherty, R.F., Myall, N.J., Wandell, B.A., Feldman, H.M., 2012. Tract Profiles of White Matter Properties: Automating Fiber-Tract Quantification. PLOS ONE 7, e49790. https://doi.org/10.1371/journal.pone.0049790

Yushkevich, P.A., Piven, J., Hazlett, H.C., Smith, R.G., Ho, S., Gee, J.C., Gerig, G., 2006. User-guided 3D active contour segmentation of anatomical structures: Significantly improved efficiency and reliability. NeuroImage 31, 1116–1128. https://doi.org/10.1016/j.neuroimage.2006.01.015

Zhang, H., Schneider, T., Wheeler-Kingshott, C.A., Alexander, D.C., 2012. NODDI: Practical in vivo neurite orientation dispersion and density imaging of the human brain. NeuroImage 61, 1000–1016. https://doi.org/10.1016/j.neuroimage.2012.03.072

Zheng, Z., Lauritzen, J.S., Perlman, E., Robinson, C.G., Nichols, M., Milkie, D., Torrens, O., Price, J., Fisher, C.B., Sharifi, N., Calle-Schuler, S.A., Kmecova, L., Ali, I.J., Karsh, B., Trautman, E.T., Bogovic, J.A., Hanslovsky, P., Jefferis, G.S.X.E., Kazhdan, M., Khairy, K., Saalfeld, S., Fetter, R.D., Bock, D.D., 2018. A Complete Electron Microscopy Volume of the Brain of Adult Drosophila melanogaster. Cell 174, 730-743.e22. https://doi.org/10.1016/j.cell.2018.06.019

Zhuang, C., Yan, S., Nayebi, A., Schrimpf, M., Frank, M.C., DiCarlo, J.J., Yamins, D.L.K., 2021. Unsupervised neural network models of the ventral visual stream. Proc Natl Acad Sci U S A 118. https://doi.org/10.1073/pnas.2014196118

